# Dissecting mechanisms of human islet differentiation and maturation through epigenome profiling

**DOI:** 10.1101/613026

**Authors:** Juan R. Alvarez-Dominguez, Julie Donaghey, Jennifer H. R. Kenty, Niloofar Rasouli, Aharon Helman, Jocelyn Charlton, Juerg R. Straubhaar, Alexander Meissner, Douglas A. Melton

**Affiliations:** Department of Stem Cell and Regenerative Biology, Harvard Stem Cell Institute, Harvard University, Cambridge, MA 02138, USA; Department of Genome Regulation, Max Planck Institute for Molecular Genetics Berlin, 14195 Germany

## Abstract

Investigating pancreatic islet differentiation from human stem cells in vitro provides a unique opportunity to dissect mechanisms that operate during human development in utero. We developed methods to profile DNA methylation, chromatin accessibility, and histone modifications from pluripotent stem cells to mature pancreatic islet cells, uncovering widespread epigenome remodeling upon endocrine commitment. Key lineage-defining loci are epigenetically primed before activation, foreshadowing cell fate commitment, and we show that priming of α-cell-specific enhancers steers polyhormonal cells toward an α-cell fate. We further dissect pioneer factors and core regulatory circuits across islet cell differentiation and maturation stages, which identify LMX1B as a key regulator of in vitro-derived endocrine progenitors. Finally, by contrasting maturing stem cell-derived to natural β-cells, we discover that circadian metabolic cycles trigger rhythmic control of insulin synthesis and release and promote mature insulin responsiveness via an increased glucose threshold. These findings form a basis for understanding mechanisms orchestrating human islet cell specification and maturation.

## INTRODUCTION

The pancreatic islets of Langerhans are responsible for physiological glucose homeostasis. They comprise α cells, which secrete glucagon in response to low blood glucose to promote glucose release by the liver; β cells, which secrete insulin in response to high blood glucose to promote glucose uptake by other tissues, and lesser amounts of δ, ε, and PP cells which secrete somatostatin, ghrelin, and pancreatic polypeptide, respectively. β-cell loss or dysfunction is accompanied by defects in other islet cell types and underlies diabetes (Ashcroft and Rorsman, 2012; Gromada et al., 2018; Rorsman and Huising, 2018), which affects >400 million people worldwide (Organization, 2016). Although many drugs exist to improve glycemic control in diabetics, none matches the precision of endogenous islet function. These patients could be cured through transplantation of new islets, a therapy that is presently limited by the scarcity and quality of islets from cadaveric sources (McCall and Shapiro, 2012).

To generate a limitless supply of islet cells, our group and others have devised strategies for directing hPSC differentiation into the pancreatic endocrine lineage (Pagliuca et al., 2014; Rezania et al., 2014; Russ et al., 2015). These were made possible by the recognition of key genes and signals regulating endoderm commitment and pancreatic progenitor specification (Pagliuca and Melton, 2013; Shih et al., 2013). Better control of deriving all islet cell types is limited by an incomplete understanding of the mechanisms driving endocrine lineage specification. In vitro preparations usually contain 20-30% β-like and 10-20% polyhormonal (insulin^+^ glucagon^+^) cells, as well as lesser amounts of α- and δ-like cells. While these stem cell-derived islet preparations do show glucose-stimulated insulin release and can cure diabetic rodents, they lack the precision and kinetics of glucose responsiveness characteristic of mature islets. Mature glucose responsiveness develops postnatally (Otonkoski et al., 1988; Rorsman et al., 1989) and involves increases in the glucose threshold for insulin secretion and in insulin secretory capacity between birth and weaning (Aguayo-Mazzucato et al., 2006; Blum et al., 2012; Stolovich-Rain et al., 2015). While factors that influence islet maturation have been characterized (MAFA, NEUROD1), the underlying mechanisms remain unclear (Liu and Hebrok, 2017).

Assessing epigenome and gene expression dynamics during development offers a powerful tool to dissect regulatory mechanisms that drive cell state transitions. For example, delineating regulatory elements and the factors that bind them has exposed key effectors driving hPSC differentiation into cardiac (Paige et al., 2012; Wamstad et al., 2012) and neural (Rada-Iglesias et al., 2012; Ziller et al., 2015) lineages. These key cell state-specific effectors gain H3K4me1 and loose DNA methylation at their regulatory sites before transcriptional activation, indicative of epigenetic priming (Creyghton et al., 2010; Wamstad et al., 2012; Zhang et al., 2012), and upon activation form extended or super enhancer clusters (Gaulton et al., 2010; Parker et al., 2013; Whyte et al., 2013) at gene loci important for cell state-specific processes, including their own, thus forming interconnected auto-regulatory loops that reflect the core transcriptional regulatory circuity for that cell state (Boyer et al., 2005; Lin et al., 2016; Neph et al., 2012; Saint-Andre et al., 2016; Tsankov et al., 2015).

To better understand mechanisms responsible for islet cell development, we exploited a protocol for the stepwise differentiation of islet cells from ∼10^8^ hPSC (Millman et al., 2016; Pagliuca et al., 2014). We developed methods to purify large quantities of islet developmental intermediates, including endocrine progenitors, stem cell-derived β (SC-β) and polyhormonal (PH) cells, which enable genome-wide profiling of DNA methylation, chromatin accessibility, histone modification, and RNA transcription. We elucidated the landscape of regulatory domains, putative pioneer factors that establish them, and their state dynamics throughout human islet development. We find that endocrine specification involves widespread enhancer resetting and is foreshadowed by the priming of lineage-specifying loci. Accordingly, we show that priming of α-cell-specific enhancers in PH cells precedes resolution toward an α-cell fate. We further infer core regulatory circuits for each islet differentiation stage, capturing both known and unexpected factors including LMX1B, which we validate as key for in vitro-derived endocrine progenitors. Finally, we contrast regulatory landscapes in SC-β with their natural counterparts to uncover a role for circadian rhythms in fostering mature insulin responsiveness. Metabolically synchronized cadaveric/SC-islets gain cyclic transcription of genes controlling insulin synthesis/release and rhythmic insulin responses with an increased glucose threshold, a hallmark of functional maturity. These data form a basis for understanding genetic and epigenetic mechanisms controlling human islet cell fate and function.

## RESULTS

### In vitro isolation of human islet developmental intermediates

We used the directed differentiation of human pluripotent stem cells (hPSC) into functional islet cells (Millman et al., 2016; Pagliuca et al., 2014) as a model to study human islet development (Figures 1A and S1A). Based on marker genes, we used flow cytometry to isolate cells at defined differentiation stages representing key intermediates during islet lineage progression (Figures S1B and S1C): undifferentiated hPSC, SOX17^+^ definitive endoderm (DE) cells, PDX1^+^ early pancreatic progenitors (PP1), PDX1^+^ NKX6.1^+^ late progenitors (PP2), NGN3^+^ endocrine progenitors (EN), and monohormonal (INS^+^ GCG^-^) SC-β as well as polyhormonal (INS^+^ GCG^+^) PH cells. To isolate live SC-β and PH, we developed a strategy that combines endocrine cell enrichment by the TSQ zinc dye (Jindal et al., 1992) with detection of DPP4/CD26, which specifically labels glucagon-expressing islet cells (Blodgett et al., 2015). SC-β and PH cells were enriched in the TSQ^+^ DPP4^-^ and TSQ^+^ DPP4^+^ populations with >90% and >85% purity, respectively (Figure S1D; Extended Experimental Procedures), thus validating our strategy.These methods enabled comprehensive epigenome profiling throughout islet cell differentiation.

**Figure 1.**
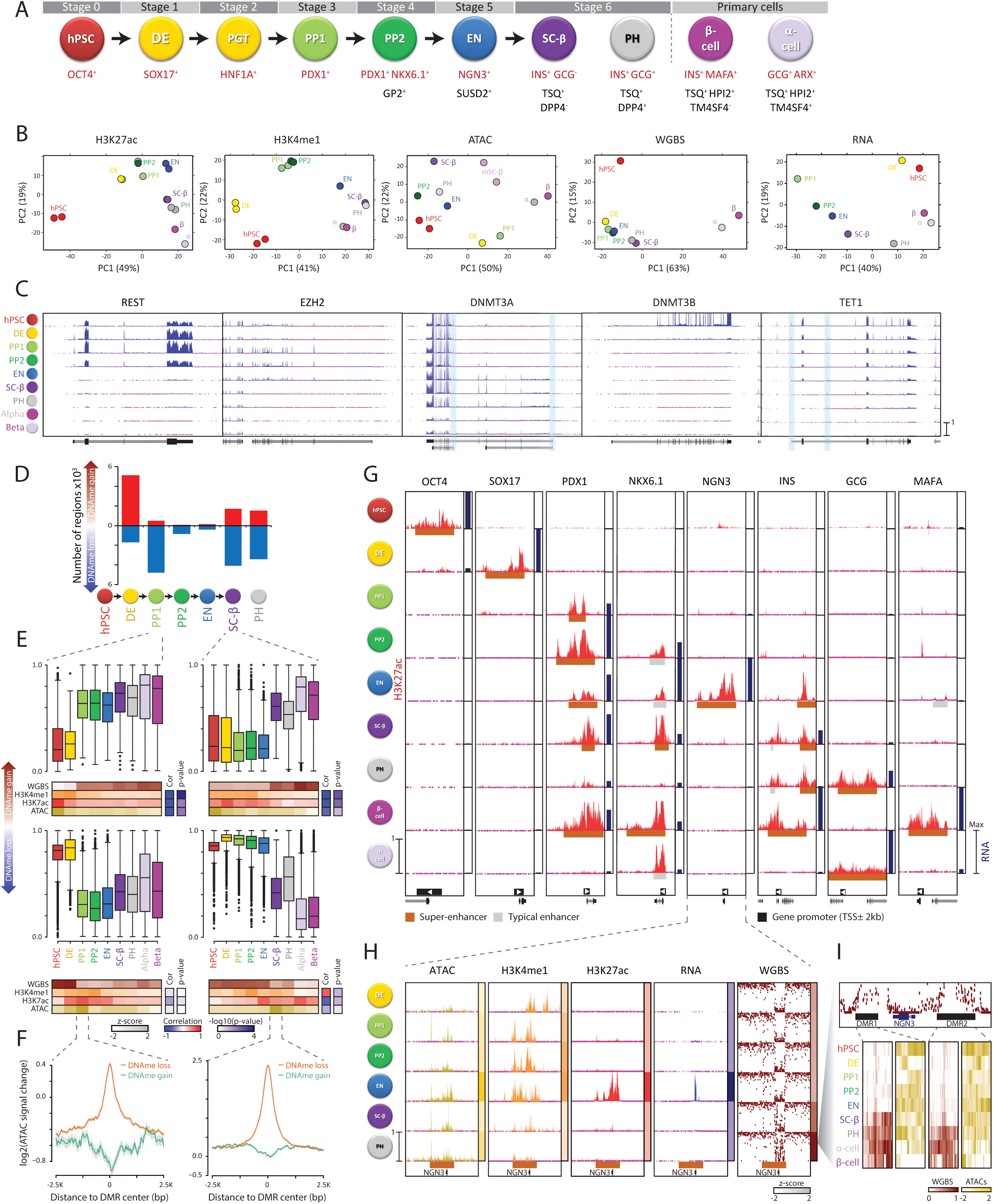
Transcriptional and epigenome dynamics during human islet differentiation. (**A**) Stages of directed differentiation from hPSC to hormone-producing islet cell types. (**B**) Stepwise developmental trajectory of islet cells from hPSC reconstructed by principal component (PC) analysis of ATAC, H3K4me1, H3K27ac, RNA, and WGBS landscapes. (**C**) Dynamic expression and isoform usage patterns of key epigenome regulators. Tracks display normalized RNA-seq signal across the gene models shown below. Alternative *DNMT3A* and *TET1* promoters are highlighted. (**D**) DNAme dynamics during directed differentiation. The number of regions that gain or lose DNAme is highest upon endoderm or pancreatic/endocrine specification, respectively. (**E**) Stable epigenetic silencing upon DNAme gain (top panels) vs. transient activation patterns upon DNAme loss (bottom panels). Boxplots show distribution of DNAme levels at all stages for regions differentially methylated during PP1 (left) and SC-β (right) specification. Heatmaps below display their median WGBS, H3K4me1, H3K27ac, and ATAC levels at each stage, with correlations between WGBS and H3K4me1/H3K27ac/ATAC patterns and their significance quantified to the right. (**F**) DNAme gain/loss coincides with focal ATAC loss/gain. Shown is the ATAC signal change around differentially methylated regions from DE to PP1 (left) and from EN to SC-β (right). (**G**) Stage-specific H3K27ac and RNA dynamics across islet development. Tracks show normalized H3K27ac signal over a region 1.5× greater than the super-enhancer domain at the stage marked by genes highlighted in (A), with relative gene expression quantified to the right. (**H**) Coordinated *NGN3* ATAC, H3K4me1, H3K27ac, RNA, and WGBS dynamics. Tracks as in (G); heatmaps to the right display relative signal over the domain shown below. (**I**) Concerted chromatin closing and DNAme gain within *NGN3*-flanking DMRs. Heatmaps show WGBS/ATAC levels at individual CpGs.

### Epigenome states during islet lineage progression

To study how gene expression is coordinated, including transcription and epigenome states, throughout islet lineage progression, we subjected hPSC-derived lineage intermediates and purified primary α/β cells (Figure S1E) to whole-genome bisulfite sequencing (WGBS), assay for transposase-accessible chromatin by sequencing (ATAC-seq), chromatin immunoprecipitation sequencing (ChIP-seq) for two histone marks (H3K27ac and H3K4me1), and directional total RNA sequencing (RNA-seq). In total, we generated 184 data sets with ∼12 billion reads aligned to the human genome (Table S1). These epigenome and transcriptome measurements showed high reproducibility and consistency (Figure S2), and independently recapitulated the stepwise developmental trajectory of islet cells from hPSC (Figure 1B).

Global mRNA expression profiling verified that in vitro-purified populations recapitulate developmental intermediates during in vivo islet lineage progression (Figures S3A-S3D; Table S2). In addition, our high-resolution total RNA maps reveal transcription across regulatory elements, non-coding genes, and splicing patterns. For example, we identify 1,056 long non-coding RNAs (lncRNAs) dynamically expressed across developmental stages (Figure S3E and Table S2). lncRNAs are recognized as vital for cell development and function (Flynn and Chang, 2014; Hu et al., 2012), including in pancreatic islets (Singer and Sussel, 2018), but have not been studied during islet cell specification/maturation. We detect >500 lncRNAs specifically induced in endocrine progenitors and maturing islet cells, which often correlate in expression with neighbor genes enriched for roles in endocrine cell development (e.g. *NGN3, ISL1, IRX2*) (Figures S3F and S3G), suggesting that some may regulate their neighboring gene (Rinn and Chang, 2012). We also identify 1,918 genes that show differential isoform use across stage transitions (Figures S3H and S3I; Table S2). These include a switch towards *DNMT3A* and *TET1* isoforms generated from alternative promoters upon endocrine commitment (Figures 1C, S3J and S3K), which reportedly confer distinct subnuclear localization/biochemical activities (Chen et al., 2002; Gu et al., 2018; Zhang et al., 2016).

We next used ChIP-seq to profile H3K27ac, which marks active promoters/enhancers, and H3K4me1, which marks either active or poised promoters/enhancers (see below) (Creyghton et al., 2010; Heintzman et al., 2009; Rada-Iglesias et al., 2011), and identified 361,576 sites enriched for H3K27ac/H3K4me1 in at least one stage. Despite consistent genomic coverage and histone modification levels during differentiation (Figure S2A, S4A-S4C), >75% of H3K27ac/ H3K4me1 regions are detected in at most 3 stages (Figure S4D). These dynamic chromatin regions are mostly distal to promoters and are overrepresented in EN and α/β cells (Figures S4D and S4E), consistent with widespread epigenetic remodeling upon endocrine commitment.

At the level of DNA methylation (DNAme), we find α/β cells to be globally hypomethylated and thus cluster separately from hPSC-derived lineage intermediates (Figures 1B and S2C), in line with hypermethylation of in vitro-derived vs. primary adult cells (FigureS5A) (Lister et al., 2011; Ziller et al., 2013). In total, we detect 126,346 differentially methylated regions (DMRs) across stage transitions (Figure 1D and Table S3), comprising narrow (median size 382bp), distal intergenic regions with focal ATAC and H3K27ac/H3K4me1 enrichment (Figures S5B and S5C). Notably, DNAme gain at DMRs was highest during exit from pluripotency, whereas DNAme loss was widespread upon pancreatic commitment and endocrine cell specification (Figure 1D). DNAme gain during these major cell-fate decisions occurs at sites transitioning from hypomethylated in all preceding stages to hypermethylated in all subsequent ones (Figures 1D and S5D, top), suggesting stable silencing of stem cell/progenitor-specific gene regulatory elements. Indeed, binding sites for OCT4/NANOG or for factors such as NEUROD1 are among the most enriched at DMRs hypermethylated in DE or in SC-β/PH, respectively, and DNAme gain at these sites is concomitant with stable loss of ATAC and H3K27ac/H3K4me1 signal (Figures 1E, 1F, S5D-S5F, top).

Decreased DNAme involves sites that re-gain DNAme upon subsequent differentiation (Figures 1E, S5D, bottom), suggesting stage-specific activity of regulatory elements. Indeed, DNAme loss coincides with localized chromatin opening and H3K27ac/H3K4me1 marking (Figures 1F and S5F), exhibits greater stage specificity than DNAme gain (Figure S5G), and occurs at sites enriched for binding of TFs known to regulate the respective stage (Figure S5E). Accordingly, recognition sites for GATA4 and FOXA1/P1/P2 are among the most enriched at PP1-hypomethylated DMRs, ONECUT1/2/3 are enriched at PP2-hypomehtylated DMRs, and MAFA is enriched at DMRs demethylated in SC-β. Supporting the functional relevance of these observations, we further find that DMR hyper/hypo methylation is preferentially linked to repression/induction of the nearest gene (Figure S5H), which include the expected stage-specific TFs (Figure S5I). These findings are well illustrated at the *NGN3* locus, wherein DNAme gain at distal flanking regions is concomitant with chromatin closing and transcriptional silencing (Figure 1I). In all, these data chart DNAme dynamics throughout islet cell differentiation and link them to the control of stage-specific regulatory elements directing stage-specific gene expression programs.

### Enhancer transitions during islet cell development

Distal enhancers are essential for coordinating stage-specific gene expression during lineage branching. Using H3K27ac enrichment outside gene promoter-proximal (TSS ±2kb) regions, we identified 51,568 putative enhancer sites which as expected show localized ATAC/H3K4me1 enrichment (Figure S6A). We then linked clustered sites to define 34,781 enhancer domains active during islet lineage progression (Table S4; Extended Experimental Procedures). These comprise broad H3K27ac regions that often overlap the gene with which they were associated and show striking stage specificity (Figures 1G, S6B-S6E). For example, the enhancer domain engulfing *SOX17* is specific to DE, the *NGN3* domain is specific to EN, and the *MAFA* domain is specific to mature β cells. Overall, 85% of enhancer domains (29,617) change activity across developmental transitions, concurring with changes in chromatin accessibility, H3K4me1 deposition, and RNA production (Figure 2A). Importantly, differential enhancer activity coincides with differential expression of the overlapping/nearest gene (Figures 1G and 2B).

**Figure 2.**
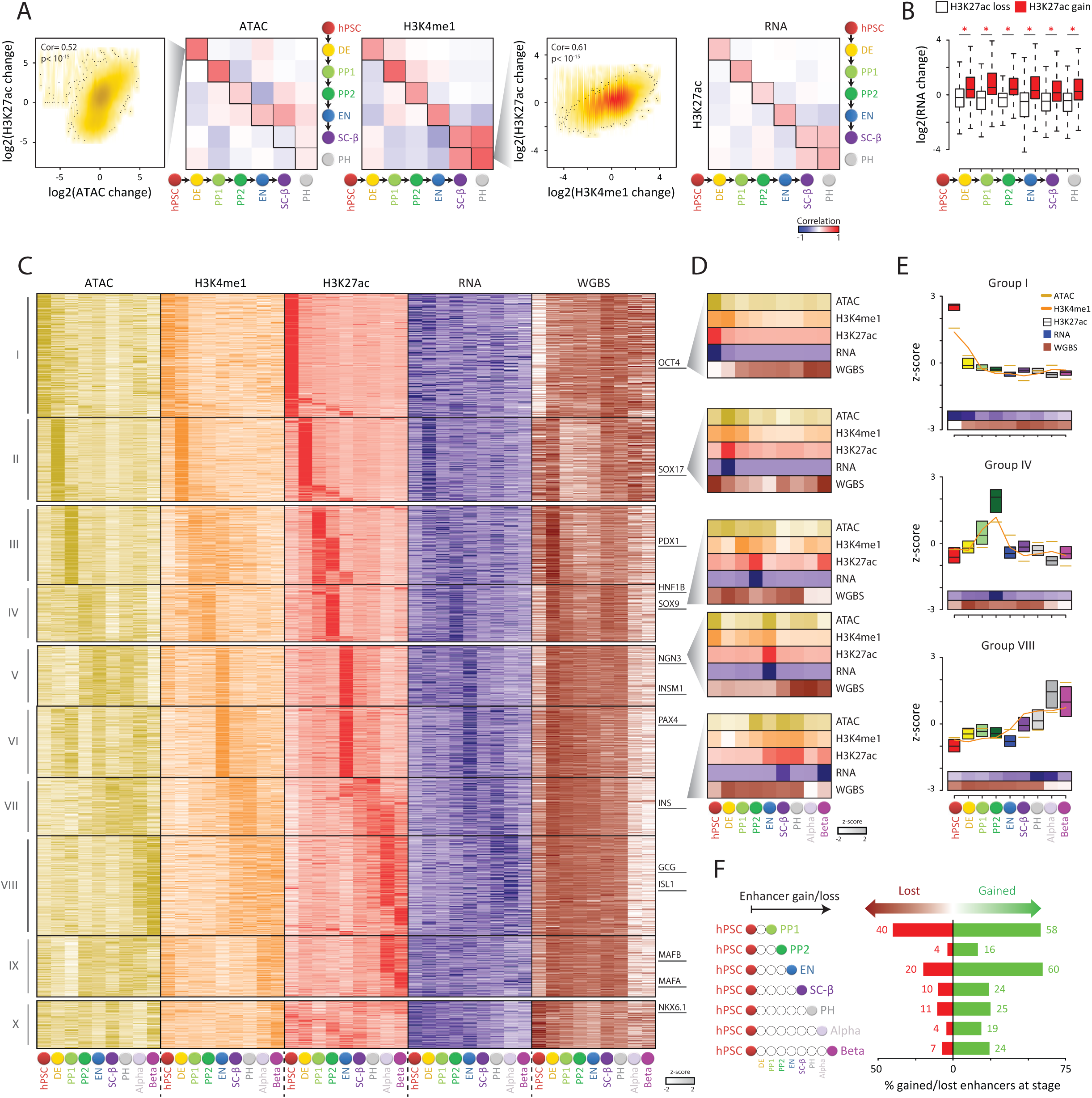
Enhancer transitions during islet cell development. (**A-B**) Synchronous changes in chromatin accessibility, modification, and transcription during directed differentiation. Correlations between H3K27ac and ATAC/H3K4me1/RNA changes across all enhancer domains for each pair of differentiation stages shown in (A). Concordance between enhancer H3K27ac loss/gain and differential expression of the overlapping/nearest gene across subsequent developmental transitions shown in (B); *p <10^-24^ (Wilcoxon test). (**C-E**) Transitioning enhancer states throughout the islet lineage. Clustering of enhancer domains based on their ATAC, H3K4me1, H3K27ac, RNA, and WGBS profiles (C). Profiles for domains associated with the genes highlighted to the right shown in (D). Epigenetic and transcriptional patterns for select enhancer clusters representing hPSC (I), pancreatic progenitors (IV), and hormone-producing islet cells (VIII) shown in (E). (**F**) Widespread enhancer turnover at lineage branching points. For enhancers gained/lost between hPSC and each stage, the fraction specifically gained/lost at that stage is shown.

Dissecting enhancer formation during islet lineage development reveals that only 10% of enhancers active in developmental intermediates are preset in stem cells; the rest are established de novo during differentiation. Enhancer establishment, marked by H3K4me1 deposition, co-occurs with focal chromatin opening and DNAme loss (Figures S7A and S7B), consistent with targeting by sequence-specific TFs (Felsenfeld et al., 1996) (Stadler et al., 2011). A stepwise acquisition of the different enhancer marks can be observed at the loci of key lineage-specifying TFs. For instance, *NGN3* distal enhancers show H3K4me1 and ATAC marking and DNAme loss at the PP1 stage, just prior to gaining H3K27ac and *NGN3* expression in endocrine-committed progenitors (Figure 1H). Similar conclusions can be drawn by examining enhancers for master regulators of other differentiation stages such as *SOX17, PDX1*, and *NKX6.1* (Figure S6E), indicating that stepwise epigenetic changes can forecast and reflect transcriptional programs that drive developmental state transitions.

Clustering enhancer domains by their ATAC, H3K4me1, H3K27ac, RNA, and WGBS profiles reveals 10 groups: I-II share enhancers established before pancreatic specification; III-IV cluster enhancers established in committed pancreatic progenitors; V-VIII comprise enhancers established upon endocrine specification and activated in progenitor/terminally differentiated endocrine cells. We also found sets of enhancers (group IX) that are lost from hPSC and re-form upon islet cell maturation, including loci specifically active in β cells (e.g. *MAFA, UCN3, SIX3*) (Figure S7C). We then asked, for each enhancer gained/lost between hPSC and differentiated progeny, the stage of gain/loss along the differentiation path (Figures 2F and S7D). Surprisingly, we find that most (∼60%) enhancers in pancreatic or endocrine progenitors are established in the first progenitor of the lineage (PP1 or EN), with reduced enhancer formation upon subsequent differentiation. Similarly, enhancer loss is higher in the first pancreatic/endocrine committed progenitor vs. subsequent progeny. Enhancer activation/deactivation follows an analogous path (Figure S7E), consistent with enhancer turnover occurring mainly at the lineage branching point. These data delineate enhancer dynamics from pluripotent to postmitotic islet cells, revealing that enhancer repertoires are largely set upon lineage commitment.

### Epigenetic priming identifies polyhormonal cells as α-cell precursors

The competence to differentiate into specific cell lineages in response to inductive cues can be linked to a gain of H3K4me1 before H3K27ac marking at lineage-specific enhancers (Creyghton et al., 2010; Wamstad et al., 2012; Zhang et al., 2012). We find that only a minority (10-20%) of enhancers specifically activated across stage transitions gain H3K4me1 in the preceding stage (Figure S8A), yet they enrich for effectors critical for executing the subsequent one (Figures 2D and S8B). Strikingly, up to 60% of newly-activated enhancers are H3K4me1-marked during earlier stages and show dynamic patterns of H3K4me1 gain/loss prior to activation (Figures 3A, 3B, S8A and S8D), revealing substantial turnover of enhancer potential. For example, *BHLHE23* and *GUSBP1* distal enhancers newly activated in EN undergo transient H3K4me1 marking at the hPSC and DE stages, respectively (Figure S8E). Importantly, H3K4me1-premarked enhancers show stronger H3K7ac activity and RNA expression than de novo enhancers (Figures 3C and S8C), consistent with a transcriptionally permissive or primed epigenetic state.

**Figure 3.**
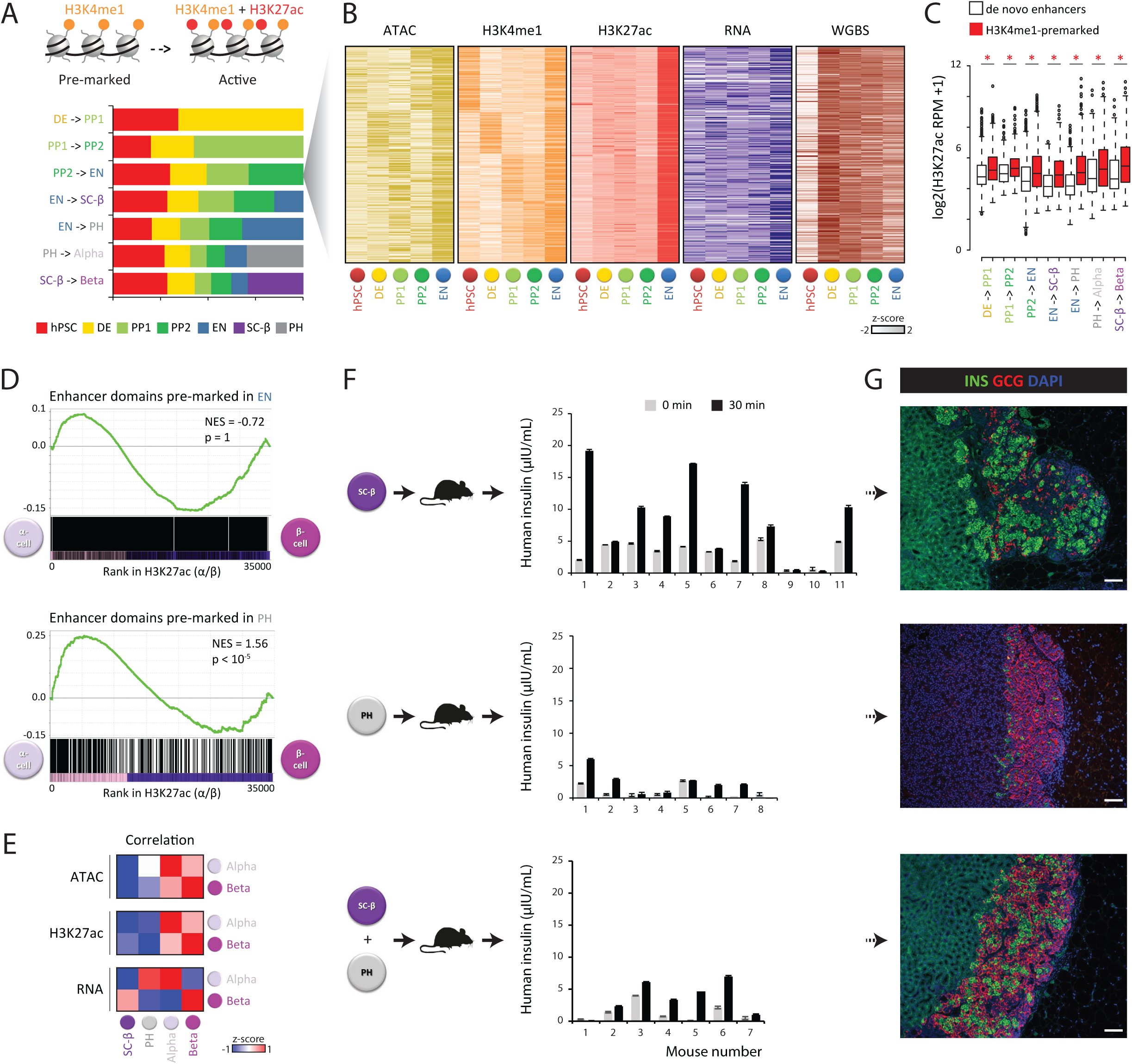
Epigenetic priming identifies polyhormonal cells as α-cell precursors. (**A-B**) Substantial turnover of enhancer H3K4me1 pre-marking. For enhancers activated at each stage transition that were previously H3K4me1-marked, the fraction whose maximal H3K4me1 deposition occurred at each previous stage is shown (A). Epigenome/RNA dynamics for 2,097 enhancers activated in EN that were H3K4me1-marked in previous stages shown in (B). (**C**) H3K4me1-premarked enhancers gain higher H3K27ac levels than de novo enhancers across stage transitions; *p <10^-8^ (Wilcoxon test). (**D**) Selective priming of α-cell-specific enhancers in PH cells. Enrichment analysis of enhancers inactive but H3K4me1-marked in EN (top) and PH (bottom) by their H3K27ac level in α vs. β cells. NES (normalized enrichment score). (**E**) PH resemble α vs. β cells. Correlations between ATAC (top), H3K27ac (middle), and RNA (bottom) levels across all enhancer domains between SC-derived and primary α/β cells. (**F-G**) PH cells resolve towards α-cells *in vivo*. Fasted mice transplanted with FACS-purified SC-β, PH, or unsorted re-aggregated cells were assayed for serum human insulin before/30min after a glucose injection 4-6 weeks post-transplantation (F). Hormonal resolution evidenced by staining retrieved grafts (G) for insulin (green)/glucagon (red). Scale bar=100μm.

We next investigated the extent to which islet lineage potential is reflected by enhancer priming in lineage intermediates. Primed-to-active enhancer transitions between EN and α/β cells are comparably likely (Figure 3D), consistent with EN multipotency, whereas priming in SC-β is biased toward β-cell-specific enhancers (Figure S9A). Intriguingly, PH cells appear selectively poised for activation of α-cell-specific enhancers (Fig. 3D), suggesting a bias toward the α-cell fate (Rezania et al., 2011). Consistent with this, PH cells closer resemble α vs. β cells by global epigenome and transcriptome metrics and preferentially express TFs restricted to α vs. β cells (Figures 3E, S2 and S3C).

To test whether PH resolve to α cells, we capitalized on our ability to purify PH and SC-β cells and transplanted them alone or in combination under the kidney capsule of immunocompromised mice (Figures 3F and S9B-S9E). Measuring human insulin in the serum of glucose-challenged mice 4-6 weeks post-transplantation showed the expected glucose-stimulated insulin secretion (GSIS) in animals receiving SC-β. By contrast, those receiving comparable numbers of PH lacked robust GSIS, while those transplanted with both SC-β ad PH showed an intermediate phenotype. Consistently, this trend persisted by the end of >2 months of observation post-transplantation (Figures S9C and S9E). To investigate hormonal resolution, engrafted kidneys were then retrieved and stained for insulin/glucagon (Figures 3G and S9D). Unlike SC-β grafts, comprised mainly of insulin^+^ monohormonal cells, PH grafts largely contained insulin^-^ glucagon^+^ cells, demonstrating α-cell resolution. We thus conclude that an epigenetic program steers in vitro-derived PH cells toward an α-cell fate.

### Identification of pioneer factors during islet cell differentiation and maturation

The acquisition of developmental competence relies on chromatin-opening pioneer transcription factors (Iwafuchi-Doi and Zaret, 2014). We searched newly-opened chromatin sites across stage transitions for TF binding motifs, and found specific enrichment for known pioneers (Figures 4A, 4B and S10A). Gained accessibility at these sites co-occurs with focal DNAme loss (Figures S10B-S10D), as illustrated at FOXA1/2 and GATA4/6 binding sites in DE, in line with pioneer activity. Beyond pioneer FOXA/GATA and HNF/MEIS factors in PP1/PP2, RFX/NEUROD factors in EN, and MAF factors in islet cells, our data highlights several unexpected TFs, including NHLH1/2 in EN and IRF1, ELF3, and AP-1 components (FOS, FOSL2, JUNB) in β cells (Figure 4C and Table S5). Notably, stage-specific gene expression patterns (Figure S10E) support roles for these factors in the hierarchy of regulators that orchestrate islet development.

**Figure 4.**
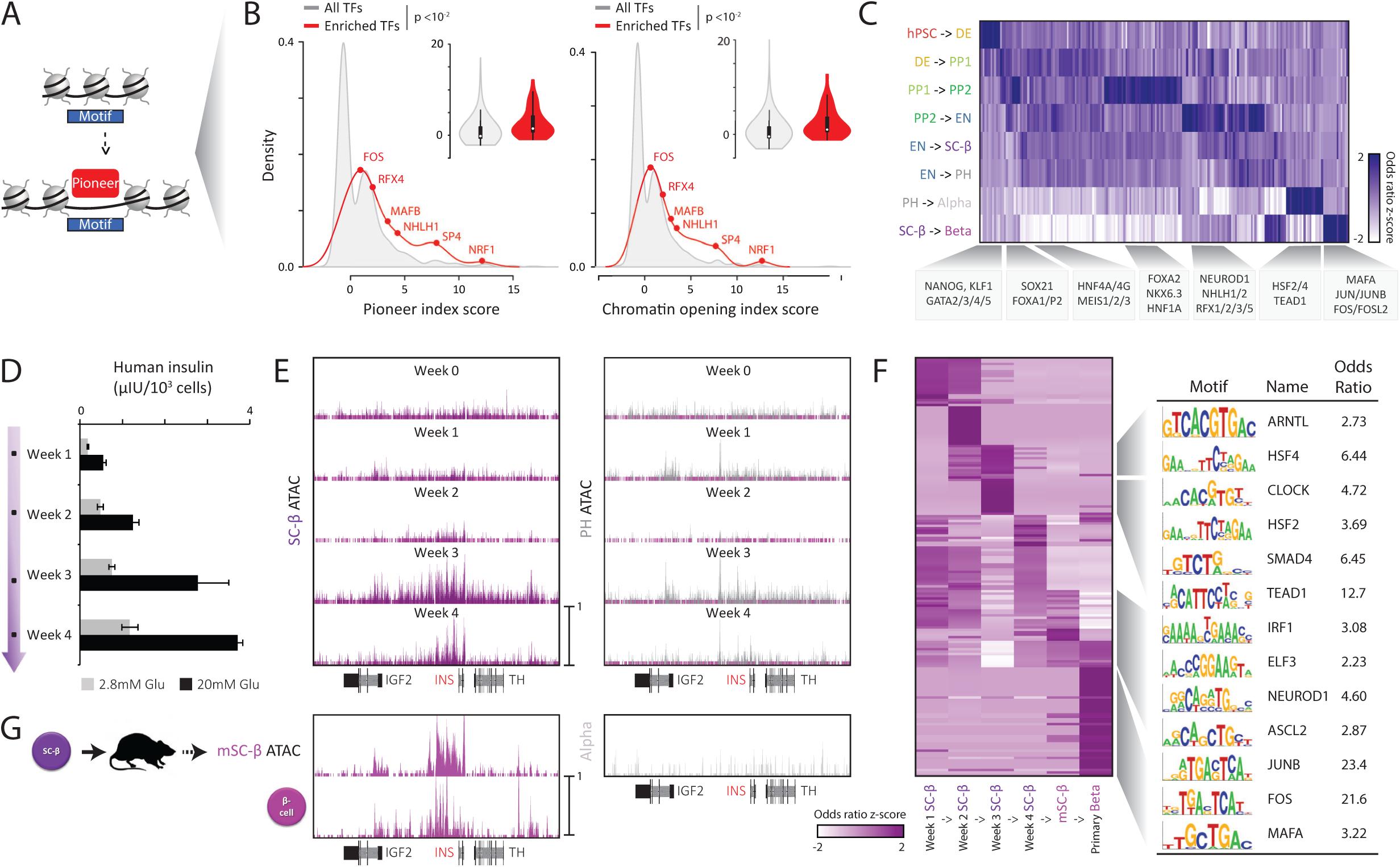
Identification of pioneer factors during islet cell differentiation and maturation. (**A-B**) Recognition motifs within newly-opened chromatin sites (A) enrich for TFs with strong pioneer (B, left) and chromatin opening (B, right) index scores (Sherwood et al., 2014). (**C**) Enrichment patterns highlight known and unexpected pioneer factors throughout islet lineage progression. Heatmap shows relative enrichment odds ratio of motifs for all expressed TFs within newly-opened regions across stage transitions. (**D**) Refinement of GSIS function during extended culture of terminal-stage SC-β preparations. Data are mean ±SEM of N=3 preparations with n=3 replicate measurements each. (**E**) Concomitant opening of regions around the insulin locus as SC-β gain robust GSIS function. Tracks display normalized ATAC signal from SC-β (left) and PH (right) cells across the regions shown below. (**F**) Pioneer factors throughout β-cell maturation. Left: TF motif enrichments as in (C) across subsequent in vitro/in vivo SC-β maturation stages, as well as between SC-β and primary β. Right: select TF motifs among the most enriched during key maturation stage transitions. (**G**) Sustained insulin openness in transplanted SC-β/primary β cells. Tracks a in (E) display ATAC-signal for in vivo-matured SC-β/primary β (left) as well as primary α (right).

To uncover pioneer factors underlying β-cell maturation, we leveraged the in vitro system to dissect changes in chromatin openness as SC-β gain glucose responsiveness. We conducted ATAC-seq in cells purified from terminal-stage cultures across a 4-week timecourse (Figure 4D). Robust function (GSIS >2 μIU/10^3^ cells and >1.5-fold stimulation) was detected by week 3, reflecting increases in both insulin content and secretory capacity (Figures 4D, S11A, and S11B). Accordingly, parallel ATAC-seq profiles reveal gained openness at sites proximal/overlapping both insulin and genes regulating its processing and secretion, in SC-β but not PH, at the 3-week timepoint (Figures 4E and S11C). The latter include calcium channel and exocyst complex components as well as calcium-dependent secretory machinery (e.g. *CADPS, SYT4, STX2*). Surprisingly, newly-opened sites during this crucial transition (weeks 2 to 3) feature CLOCK and ARNTL/BMAL1, the core circadian clock activators, among the top-enriched TF-binding motifs, which also include heat-shock factors HSF2/4 and signaling-linked TFs SMAD4 and TEAD1 (Figure 4F; Table S5). Well-known controllers such as MAFA and NEUROD1, by contrast, only begin to be enriched by week 4, suggesting incomplete maturation. Indeed, both insulin content and secretion of week-4 SC-β remain below those of primary β cells (Figures S11A and S11B).

We thus sought to profile further changes upon in vivo maturation. To this end, we transplanted preparations containing 5×10^7^ SC-β cells into immunocompromised rats, which enabled recovery of enough engrafted cells for ATAC-seq (Figures 4G, S12A and S12B; Extended Experimental Procedures). GSIS was detected 2-4 weeks post-transplantation and persisted after 2-4 months of observation, verifying in vivo SC-β function. Subsequent ATAC profiling confirms that transplanted SC-β closer resemble natural β cells (Figures 1B and S2B), revealing 41,150 newly-opened sites overrepresented at TF (e.g. *ISL1, PAX6*) and secretory (e.g. *UCN3, SLC30A8*) genes (Figures 4G, S12B, S12C; Table S5). Unlike those gained in vitro, these sites enrich for binding motifs of late-onset factors both well-known (e.g. NEUROD1, MAFA) and uncharacterized (e.g. IRF1, ELF3), many of which are shared with primary β cells (Figures 4F, S12C-S12E). For instance, NEUROD1/MAFA/IRF1/ ELF3 target sites at the *IAPP* and *UCN3* loci become active specifically in transplanted SC-β/primary β cells, suggesting roles in activating these markers. Conversely, AP-1 sites appear specifically opened in primary β cells. These data identify pioneer TFs throughout islet lineage progression, highlighting potential regulators of the onset and refinement of β-cell function.

### Core regulatory circuits in the human islet lineage

Master TFs are thought to program stable cell states using autoregulatory loops that involve cooperative formation of extended or super enhancer clusters (Gaulton et al., 2010; Parker et al., 2013; Whyte et al., 2013). A cell’s core regulatory circuit (CRC) can thus be reconstructed by finding SE-driven TFs in interconnected autoregulatory loops (Boyer et al., 2005; Lin et al., 2016; Saint-Andre et al., 2016). We implemented this logic via a three-tier system (Figure 5A). First, we ranked enhancer domains by H3K27ac density to identify SEs, which as expected are associated with genes important for cell state-specific processes (Figures 1G and S13). We then used binding motifs to predict SE-driven TFs that bind their own SE and are thus autoregulated. Finally, we identified groups of autoregulated SE-driven TFs that bind each other’s SE to form fully-interconnected autoregulatory loops. The resulting CRCs indeed capture master TFs of each differentiation stage (Figures 5B and S14; Table S6). For example, PP2 CRCs comprise PDX1, SOX9, and ONECUT1/HNF6, while possible β-cell CRCs comprise MAFA, NKX6.1, and PAX6. Importantly, binding relationships among CRC TFs are supported by ChIP-seq (Figures S14A-S14F), and depleting a given TF triggers selective depletion of the others in its CRC (Figure S14G), supporting strong interconnectivity (Figure S14H).

**Figure 5.**
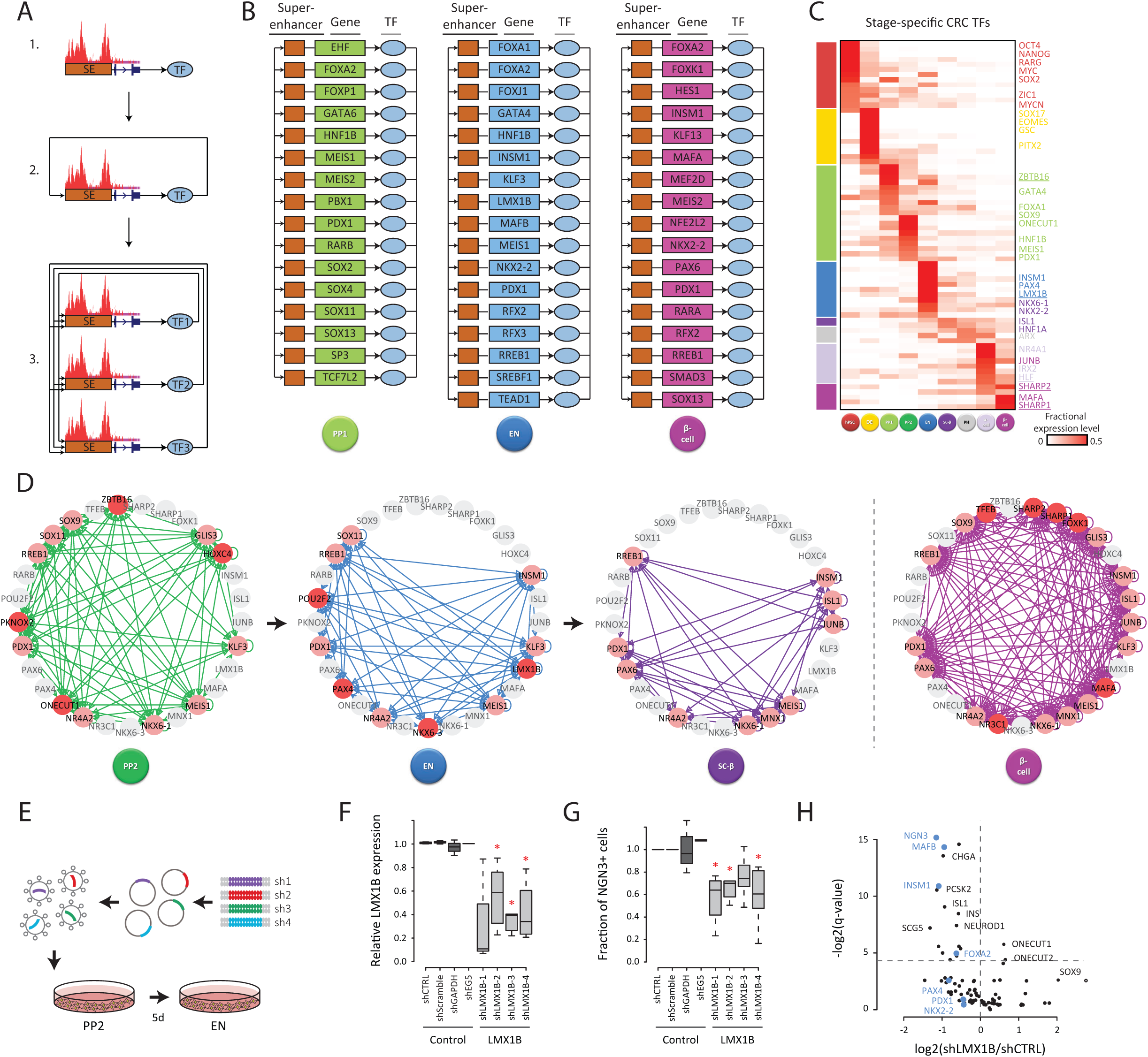
Core regulatory circuits identify LMX1B as a critical regulator of human endocrine progenitors. (**A-B**) Reconstructing CRCs (Saint-Andre et al., 2016) across the human pancreatic lineage. 1) SE-driven TFs are identified; 2) autoregulated TFs are identified as TFs predicted to bind their own SE.; 3) CRCs are defined as groups of autoregulated TFs predicted to bind each other’s SE to form fully-interconnected autoregulatory loops (A). Representative CRC models for PP1, EN, and primary β shown in (B). (**C**) Lineage-specifying CRC TFs are distinguished by their stage-specific expression patterns (see Figure S15 for all TFs). Known lineage-specifying TFs are highlighted to the right; newly implicated TFs are underlined. (**D**) Dynamic CRC rewiring through the stages of pancreatic development. Nodes represent stage-specific CRC TFs from (C) that partake (pink) or not (gray) in PP2, EN, SC-β, or primary β-cell CRCs. Edges represent predicted transcriptional regulatory relationships between TFs within the same CRC. TFs unique to each stage are highlighted in red. (**E-H**) LMX1B is a critical regulator of in vitro-derived human EN progenitors. shRNA studies (E) show that *LMX1B* knockdown in shRNA-transduced EN-stage cells (F) depletes the fraction of NGN3+ progenitors (G) and blocks induction of EN CRC TFs (H, highlighted in blue). Boxplots: data from N=2-9 experiments with n=1-3 replicate measurements each. *p <0.05 relative to control shRNA (t-test). Volcano plot: data averaged over 4 distinct shRNAs with N=2-4 experiments each.

How are CRCs remodeled during development? We reasoned that lineage-inductive cues could trigger activation of new groups of CRC-forming TFs that in turn reprogram cell sate. Examining expression specificity reveals that key CRC lineage-specifiers are indeed cell state-specific (Figures 5C and S15). For example, TFs specifying endoderm (GSC, EOMES, SOX17) and pancreatic (PDX1, HNF6, SOX9) fates are highly specific to EN and PP1, respectively, confirming numerous earlier studies (Oliver-Krasinski and Stoffers, 2008). This analysis also casts unforeseen TFs as lineage-specifiers (Figures 5B and 5C), such as LMX1B for EN and the circadian factors HLF for α as well as BHLHE40/SHARP2 and BHLHE41/SHARP1for β cells. These TFs show high stage-specific interconnectivity (Figures 5D and S14I), including with key TFs that partake in CRCs at multiple stages (e.g. PDX1, NKX6.1, PAX6).

We tested whether LMX1B, specifically active in NGN3^+^ progenitors (Figures 5C, S16A, and S16B), regulates the EN state, by expressing *LMX1B*-targeted shRNAs during endocrine differentiation (Figure 5E). Depleting *LMX1B* by ∼50% reduced the fraction of NGN3^+^ cells by ∼50% (Figures 5F and G), reflecting proportional NGN3 depletion at the single-cell level (Figures S16C and S16D). Accordingly, the core endocrine gene program was disrupted, as evidenced by selective downregulation of EN CRC TFs (*INSM1, MAFB*) and suppressed induction of pan-endocrine TFs (*ISL1, NEUROD1*), secretory proteins (*CHGA, PCSK2*), and hormones (insulin, glucagon) (Figures 5H and S16E). Conversely, the PP2 CRC TFs *ONECUT1* and *SOX9* remain active in *LMX1B*-depleted cells, indicating a block in differentiation. We thus conclude that LMX1B acts as a key regulator of in vitro-derived human EN progenitors.

### Circadian rhythms trigger an islet maturation step

Circadian rhythms are vital for attuning insulin secretion to daily feeding/fasting cycles (Bass and Takahashi, 2010; Gamble et al., 2014). A role for circadian clock controllers in islet maturation is suggested by their expression patterns (Figures S3B and S10E), their enrichment at regulatory sites gained as SC-β raise their insulin production/secretion (Figure 4F), and their interconnections with key maturation-linked TFs in the α/β CRCs (Figures 5C and 5D). We thus asked if clock synchronization can promote functional islet maturation in vitro. First, we verified that insulin responsiveness in human cadaveric and stem cell-derived islet (SC-islet) preparations can be chemically synchronized (Figures S17A and S17B), extending prior work in rodents (Perelis et al., 2015). Synchronized cultures grown in constant media show time-of-day-dependent variation in insulin content and secretion in response to glucose/KCL, with peak production/secretion at the 12h circadian timepoint. Enhanced GSIS vs. parallel mock-treated cultures can be detected at this time by dynamic release and calcium influx assays (Figure S17B). We then discovered that prior synchronization by recurring shocks of various stimuli (glucose, arginine, forskolin, insulin) can further amplify insulin responsiveness, as much as 6-fold after recovery from forskolin/glucose shocks (Figures 6A, 6C, S17C, and S17D). Strikingly, this reflects reduced basal responses to low (2.8mM) glucose without significant change in overall secretion under stimulatory (20mM glucose/30mM KCL) conditions or overall insulin content (Figures 6B and S17E). Synchronization can thus endow cultured islets with an increased glucose threshold for insulin secretion, a hallmark of functional maturity (Blum et al., 2012).

**Figure 6.**
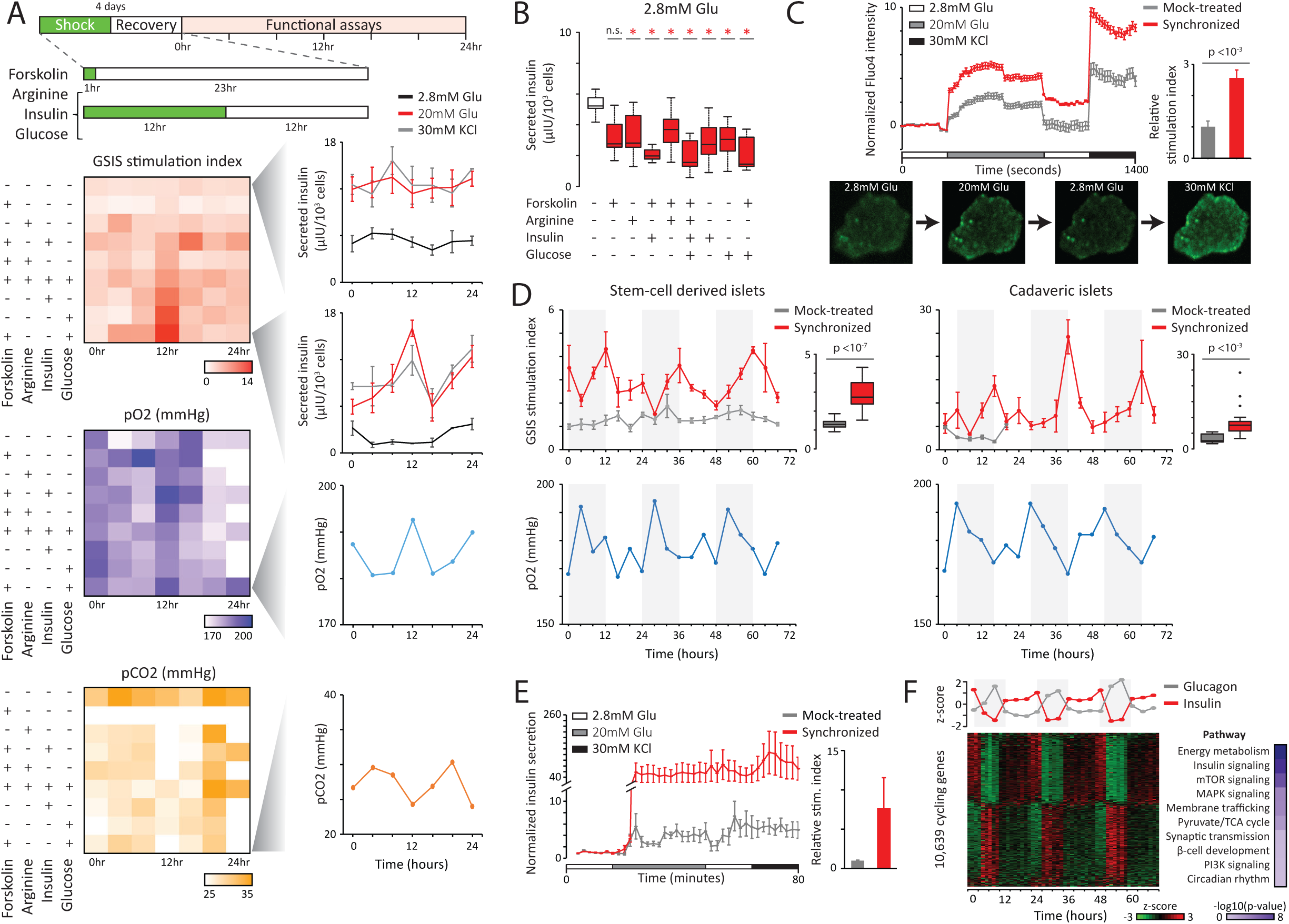
Circadian rhythms trigger an islet maturation step. (**A**) Human islet GSIS and respiration rhythms induced by synchronization with various stimuli. Schematic: timeline for recurrent shock/recovery cycles followed by functional assays across 24h. Panels: GSIS stimulation indexes (top), partial O2 pressure in the medium of unstimulated cultures (middle), and partial CO2 pressure (bottom), with patterns for mock-treated vs. synchronized cultures shown to the right. Data are mean ±SEM of n=3 replicate measurements. (**B**) Reduced insulin response to basal glucose levels in synchronized islet cultures. Boxplots summarize insulin secretion across 24h for the indicated conditions; *p <0.01 relative to mock-treated (t-test). (**C**) Synchronization enhances islet glucose responsiveness. Top: calcium signaling traces from mock-treated vs. synchronized islets during the indicated incubations, with relative stimulation indexes quantified to the right. Data are mean ±SEM of n=7 islets sampled from each culture at the 12h circadian timepoint, normalized to mean of the first incubation. Bottom: representative Fluo4-staining images. (**D**) Rhythmic GSIS responses in synchronized cadaveric/SC-islet cultures across 72h. Top: GSIS stimulation indexes for synchronized vs. mock-treated SC-islets (left) and cadaveric islets (right), summarized to the right of each. Data are mean ±SEM of n=3 replicate measurements. Bottom: partial O2 pressure in the medium of unstimulated synchronized SC-islets (left) and cadaveric islets (right). (**E**) Synchronization enhances SC-islet glucose responsiveness. Insulin secretion from mock-treated vs. synchronized SC-islets during the indicated incubations, with relative stimulation indexes quantified to the right. Data are mean ±SEM of N=3 preparations sampled at the 12h circadian timepoint with n=2 replicate measurements each, normalized to mean of the first incubation. (**F**) Rhythmic RNA expression in synchronized SC-islet cultures across 72h. Top: antiphasic insulin and glucose expression; data are mean of N=3 preparations. Bottom left: RNA expression patterns for all cycling genes; data pooled from N=3 preparations. Bottom right: enriched pathways within the cycling gene set.

We next tested our optimized forskolin/glucose synchronization strategy in SC-islets. The results in Figures 6D and S18A show sustained cycling of insulin production/secretion over 72h, causing rhythmic GSIS function. Remarkably, as with natural islets we detect lower basal insulin secretion and enhanced responsiveness, which at peak secretion can reach stimulation indexes 10-fold greater than mock-treated counterparts (Figures 6D, 6E, S18B). Of note, cadaveric/SC-islets that lack GSIS function do not respond to synchronization (data not shown), in line with roles in tuning but not eliciting insulin responsiveness. Interestingly, O_2_/CO_2_ levels in the medium of unstimulated synchronous cultures are also rhythmic (Figures 6A and 6D), linking GSIS rhythms to respiration cycles. Indeed, parallel RNA-seq profiling identifies energy metabolism enzymes among the 10,639 genes specifically induced to oscillate upon synchronization (Figures 6F, S19A-S19C; Table S7). These include glucose import/metabolism (e.g. *GCK, PDHA1/B/X*) and TCA cycle enzymes, as well as electron transport chain and ATP synthase components. Consistent with their cycling, pathway analysis reveals an overrepresentation of mTOR/PI3K/AKT and MAPK signaling factors among rhythmic RNAs.

Beyond energy metabolism, we detect antiphasic expression of insulin and glucagon (Figure 6F), as well as widespread cycling of factors that mediate insulin metabolism (*PCSK1, IDE*), signaling, and transport/secretion (Figure S19C). Surprisingly, the circadian transcriptome includes *PDX1, NKX6.1*, and maturity-linked factors (*NEUROD1, IAPP*), all of which show high-amplitude oscillations (Figure S19D). These data provide a molecular basis for circadian variation in insulin responsiveness, via the rhythmic control of its synthesis, transport, and release. Circadian regulation can thus recreate physiological changes reminiscent of postnatal maturation to restore/foster robust insulin responses, revealing a mechanistic component of the mature islet phenotype.

## DISCUSSSION

Realizing the promise of in vitro-derived islets for regenerative medicine will benefit from a complete understanding of the mechanisms governing islet development and function. Directed stem cell differentiation offers an unprecedented human platform to dissect these processes. Prior directed differentiation studies have described epigenetic processes influencing pancreatic specification (Cebola et al., 2015; Wang et al., 2015; Xie et al., 2013; Xu et al., 2014). Extending these studies to the specification of endocrine cell types has been hampered by their inefficient/incomplete derivation, which results in highly heterogenous preparations (Pagliuca et al., 2014; Rezania et al., 2014; Russ et al., 2015). Here, we circumvent these challenges with cell purification strategies that enable deep epigenome profiling throughout the specification, terminal differentiation, and functional maturation of α-like and β-like cell types. Our results reveal a chromatin-level determination of endocrine cell fates, providing insights into the dynamic regulation of cellular differentiation and functional specialization programs.

How chromatin changes coordinate genetic circuits in response to developmental cues to steer cell fate is little understood. Using combinations of a few specific growth factors and small molecules in separate steps to induce stem cells to islet cell fates, we describe the epigenome changes that accompany this directed differentiation and discover chromatin transitions that precede changes in transcription. The results show that lineage decisions involve widespread epigenome remodeling. Two major waves of enhancer gain/loss mark the pancreatic/endocrine commitment points, entailing concurrent turnover of chromatin accessibility, histone marking, and DNAme. These changes gate access of stage-specific TFs to their target sites, linking chromatin transitions to the rewiring of transcriptional circuits driving cell fate. This model implies that enhancer repertoires are not preset but rather set de novo upon lineage branching (Choukrallah et al., 2015; Gonzalez et al., 2015; Lara-Astiaso et al., 2014; Luyten et al., 2014).

Our data also reveal that pancreatic lineage-defining TFs gain a primed chromatin state before activation, foreshadowing cell fate commitment. Primed-to-active transitions are limited, yet they reliably mark stage-specific regulators across diverse lineages (Wamstad et al., 2012; Zhang et al., 2012; Ziller et al., 2015). Future studies will be necessary to dissect which developmental/environmental factors are responsible for priming events and how they do so. Such events may exist to facilitate robust concerted activation of critical fate-changing factors. Supporting this, we find that insulin^+^ glucagon^+^ cells primed for activating α-over β-selective factors indeed resolve towards an α-cell fate. Though preparations containing such polyhormonal cells reportedly yield mainly insulin^-^ α-like cells upon transplantation/extended culture (Rezania et al., 2011), in principle these could derive from other precursors in the preparations. Our transplantation studies resolve this controversy by directly tracking the fate of purified PH cells. Of note, they also show better GSIS function when SC-β are transplanted alone vs. with PH, suggesting potential suppressive effects of the latter.

The programming of cell fate has been recently linked to core regulatory circuits formed by super-enhancer-driven TFs (Lin et al., 2016; Saint-Andre et al., 2016). How these circuits are rearranged to program sequential cell fates during development is unclear. Our data shows that islet lineage CRCs are dynamically rewired by the stage-specific gain/loss of SE-driven TFs, suggesting that a few stage-specific factors can drive the reconfiguration of otherwise stable CRCs during differentiation (Adam et al., 2015; Goode et al., 2016). One such factor, LMX1B, is indeed critical for in vitro differentiation of endocrine progenitors. Interestingly, LMX1B controls dopaminergic/serotonergic neuron development downstream of NKX2.2 (Cheng et al., 2003; Smidt et al., 2000), and can trans-activate insulin ectopically (German et al., 1992), suggesting a related pathway for its activation and function during islet development.

Glucose responsiveness is the key feature of stem cell-derived islets, yet a detailed understanding of how this function develops has been lacking. We show that SC-islets refine their glucose responsiveness over time by increasing insulin synthesis and secretory capacity, and this is linked to the progressive activation of regulatory elements at both insulin and secretory genes by putative pioneer factors. Gaining control over these factors, which include circadian clock and TGF-β/Hippo signaling effectors that are readily tunable, will help improve the production of mature SC-islets, and harnessing of circadian control of insulin responsiveness marks a first step towards this goal.

Circadian variation in insulin responses has been linked to rhythmic control of insulin production/secretory capacity by islet-autonomous clocks that are established postnatally (Marcheva et al., 2010; Perelis et al., 2015; Peschke and Peschke, 1998; Polonsky et al., 1988; Rakshit et al., 2018). Accordingly, β-cell-specific *Arntl/Bmal1* deletion in rodents yields islets with impaired insulin secretory capacity, reminiscent of neonatal/functionally immature islets (Marcheva et al., 2010; Perelis et al., 2015; Rakshit et al., 2018). Our in vitro studies permit further dissection of islet function modulation by the clock in a controlled setting. We show that clock/metabolic synchronization triggers pulsatile insulin synthesis and release in human islets cultured ex vivo, restoring robust GSIS responses via an increased glucose threshold. The same mechanism operates and boosts the function of stem cell-derived islets, which otherwise increase basal insulin secretion as they expand their production/secretory capacity. Thus, insulin responsiveness is amenable to improvement by circadian/metabolic modulation.

Overall, our work provides insights that underpin human islet development and function. These insights may be valuable in developing improved β-cell programing strategies and understanding how disrupted genetic circuits contribute to metabolic diseases including diabetes.

## EXPERIMENTAL PROCEDURES

Detailed experimental and analytical methods can be found in the Extended Experimental Procedures.

### Cell culture and purification

Directed hPSC differentiation into islet cells and culture of primary adult islets from cadaveric donors was conducted essentially as described in (Pagliuca et al., 2014). Cell clusters sampled from these cultures were dispersed into single-cells by incubation in PBS +50% Accutase (StemCell Technologies) and purified using MoFlo flow cytometers (Beckman Coulter) according to the strategies outlined in Figure S1.

### ChIP-seq and ATAC-seq

ChIP-seq was conducted essentially as described in (Gifford et al., 2013). Sequencing libraries were prepared using a NEBNext Ultra II DNA Library Prep Kit (New England Biolabs; E7103). ATAC-seq was conducted as described in (Buenrostro et al., 2013). Libraries were sequenced on a HiSeq 2500 instrument (Illumina), and aligned to the human genome using bowtie2 (Langmead and Salzberg, 2012).

### WGBS

WGBS was conducted as described in (Donaghey et al., 2018). Sequencing libraries were prepared using an EZ DNA Methylation-Gold kit (Zymo Research), sequenced on a HiSeq 4000 instrument, and aligned to the human genome using BSMap (Xi and Li, 2009).

### RNA-seq

RNA-seq was conducted essentially as described in (Alvarez-Dominguez et al., 2017). Sequencing libraries were prepared using a stranded RNA-seq library preparation kit (KAPA Biosystems), sequenced on a HiSeq 2500 instrument, and aligned to the human genome using Tophat2 (Kim et al., 2013).

### Animal transplantation

Cell transplantation into the kidney capsule of immunodeficient SCID-Beige mice (Jackson Laboratory) or Rowett Nude rats (Charles River), and kidney retrieval, sectioning, and histology were conducted essentially as described in (Pagliuca et al., 2014). For ATAC-seq, grafts were first mechanically triturated and dissociated into single-cells by incubation in PBS +50% Accutase and then sorted using MoFlo flow cytometers as outlined in Figure S12B.

### Glucose-stimulated insulin secretion

GSIS measurements from cadaveric/SC-islets and serum human insulin detection in fasted transplanted animals following an intraperitoneal glucose challenge were conducted as described in (Pagliuca et al., 2014).

### Lentiviral transduction

Cell clusters sampled from suspension cultures were dispersed, transduced with concentrated shRNA-expressing lentiviruses harvested from LentiX-293T cells (Takara Bio; 632180), and cultured in BD Matrigel Matrix High Concentration (BD Biosciences) with daily feeding.

### NanoString and qPCR

Total RNA from target cells was isolated using a miRNeasy Mini Kit (Qiagen) and hybridized to a custom probe set for expression analysis on the nCounter Digital Analyzer (NanoString Technologies) or cDNA-converted for detection on the 7900HT Fast Real-time PCR System (Applied Biosystems).

### Calcium influx

Intracellular calcium was measured by imaging cadaveric/SC-islets stained with Fluo4-AM (Life Technologies) on an AxioZoom V16 microscope (Carl Zeiss) as in (Pagliuca et al., 2014).

### Islet synchronization

Cadaveric/SC-islet suspension cultures were treated as outlined in Figure 6A. After synchronization treatments, cultures were kept in constant media and cell/supernatant samples were taken every 4h for downstream assays.

## AUTHOR CONTRIBUTIONS

J.R.A.-D., J.D., J.H.R.K., N.R., and A.H. performed experiments. J.R.A.-D., J.D., J.C., and J.R.S. conducted bioinformatics analyses. J.R.A.-D., J.D., J.H.R.K., A.M., and D.A.M. designed the research, interpreted results, and wrote the manuscript.

## ACKNOWLEDGEMENTS

We thank Deanne Watson, Dani Swain, Jeff Davis, and Ramona Pop for reagents and assistance with experiments; the HSCI flow cytometry, genome technology, and histology cores for technical support and critical discussions; and N. Slavov, A. Yuan, and members of the Melton and Meissner labs for feedback on this manuscript. J.R.A-D. is a Howard Hughes Medical Institute Fellow of the Life Sciences Research Foundation. D.A.M. is an investigator of the Howard Hughes Medical Institute. This work was supported by the Max Planck Society and NIH grants DP3K111898 and P01GM099117 (A.M.) and by grants from the Juvenile Diabetes Research Foundation, Helmsley Charitable Trust, and the JPB Foundation.

---

**Figure S1.**
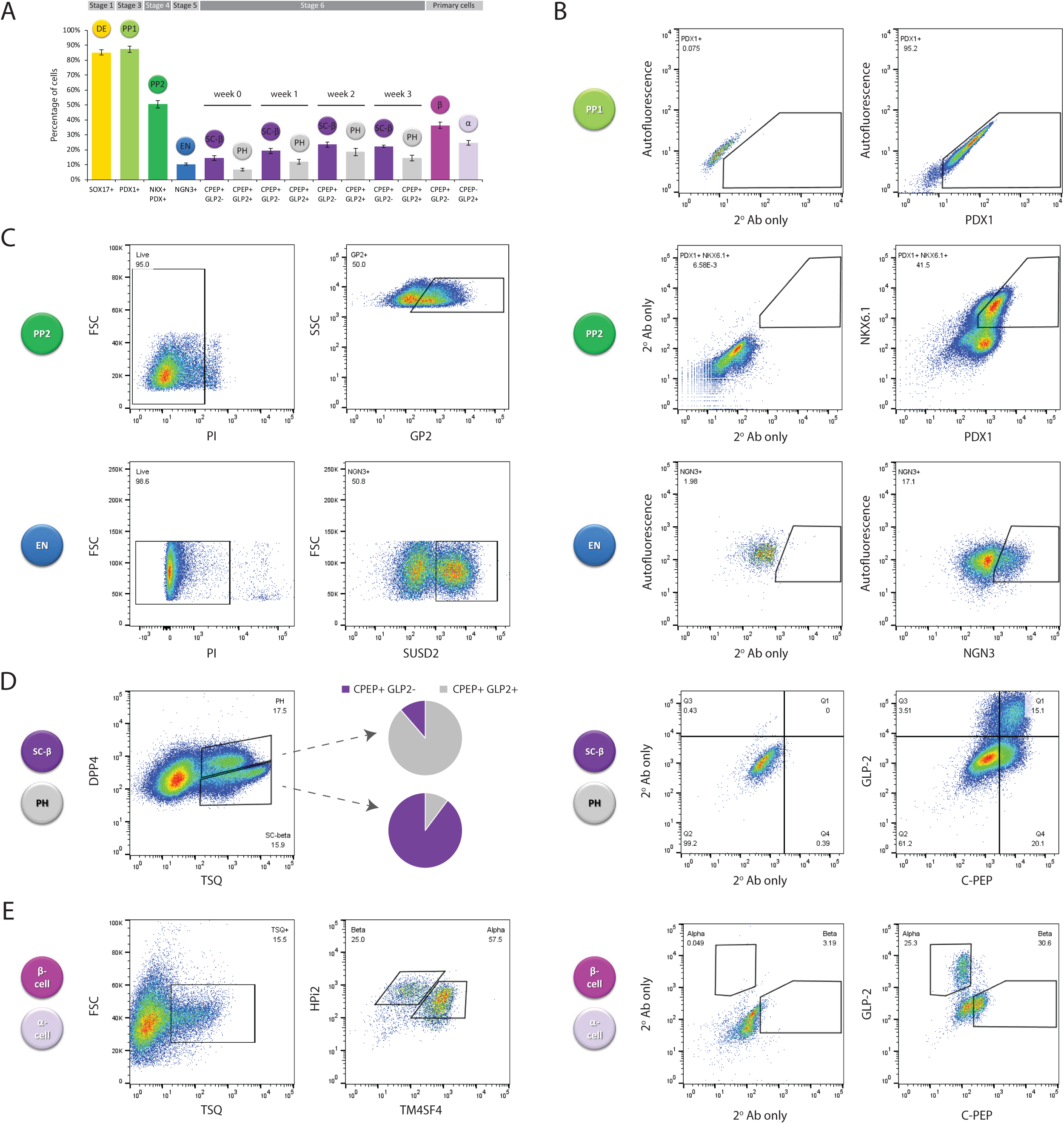
In vitro isolation of human islet developmental intermediates, related to Figure 1. (**A**) Target subpopulations in the directed differentiation from hPSC to hormone-producing islet cell types. (**B-C**). Sorting strategies for isolating fixed (B) or live (C) cells from the shown subpopulations by intracellular or surface staining for the indicated proteins, respectively. (**D**) Strategy for sorting SC-β from PH live cells based on the indicated markers, with representative population purities, assessed by post-sorting intracellular staining for the indicated proteins, shown to the right. (**E**) Sorting strategies for isolating live (left) and fixed (right) primary α/β cells, based on surface/intracellular staining for the indicated proteins.

**Figure S2.**
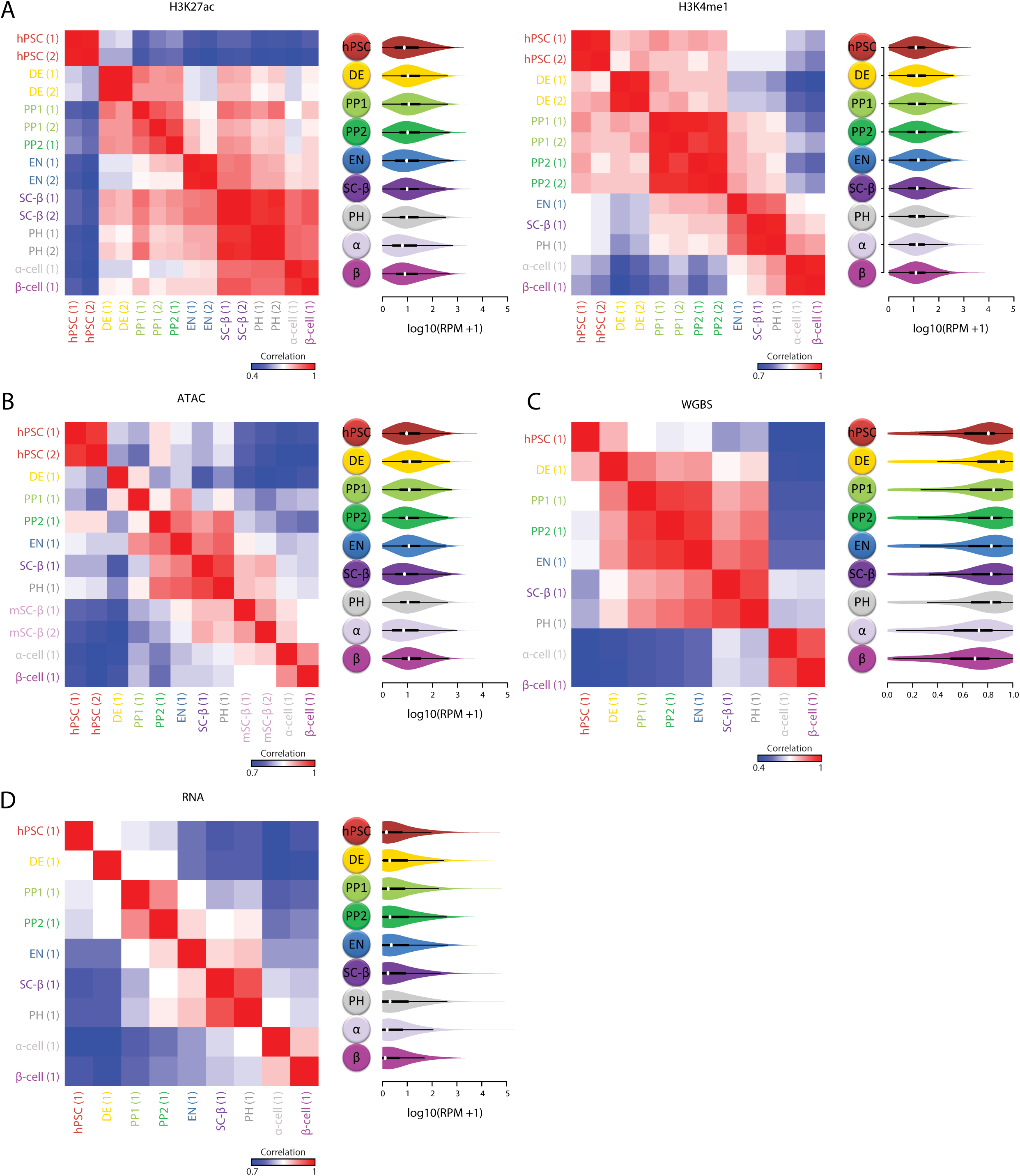
Consistent and reproducible transcriptional and epigenome states in the developing islet lineage, related to Figure 1. (**A-D**) Correlation of H3K27ac (A left), H3K4me1 (A right), ATAC (B), WGBS (C), and RNA (D) levels in 34,781 enhancer domains across all replicates and cell types analyzed in this study, with distributions summarized to the right of each heatmap.

**Figure S3.**
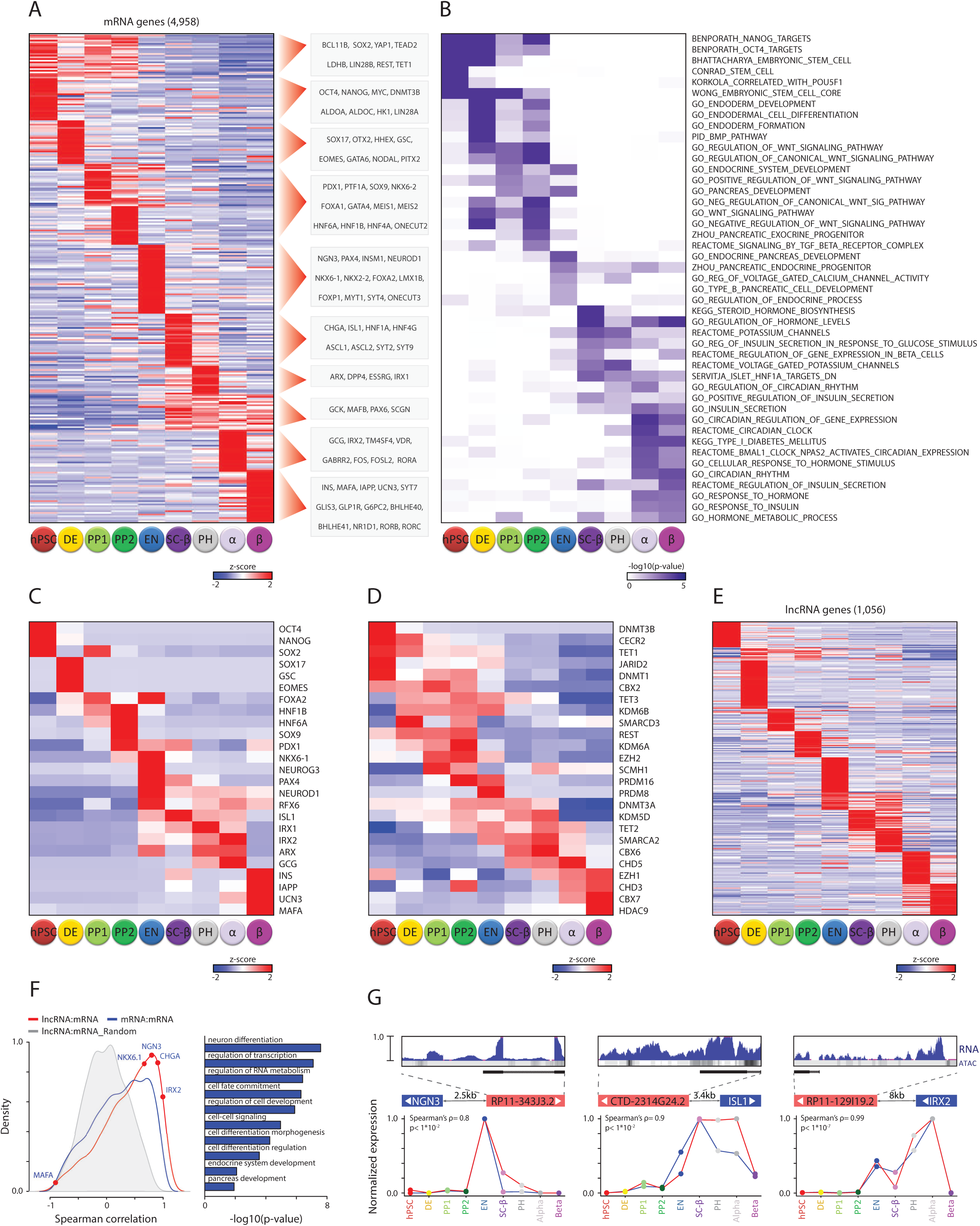
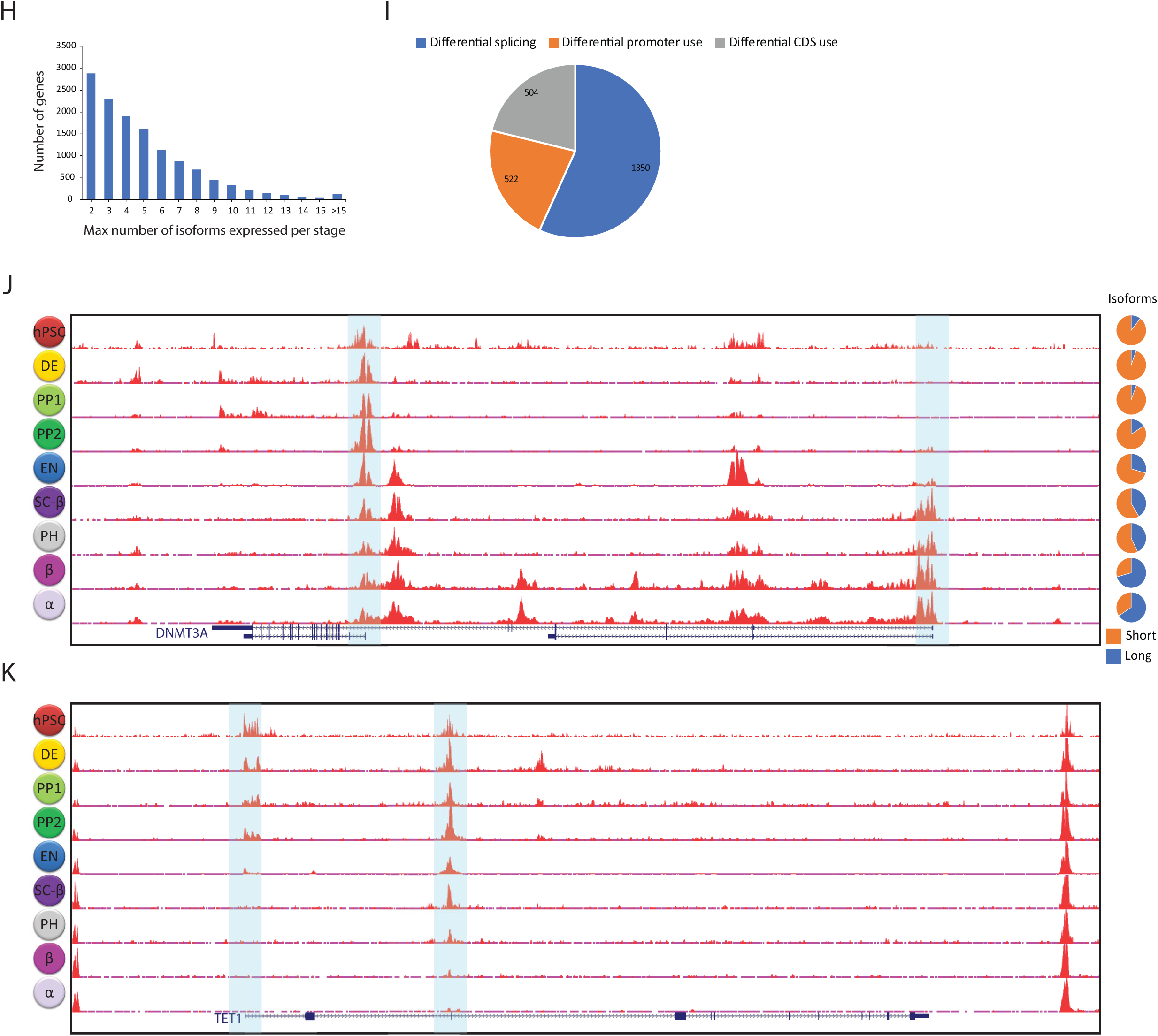
Transcriptional dynamics during islet lineage progression, related to Figure 1. (**A-B**) Differentially expressed mRNAs across developmental stages (A) and significantly enriched gene sets (B). (**C-E**) Dynamic expression of key TFs and markers (C), epigenome regulators (D), and lncRNAs across developmental stages. (**F**) Correlation of expression patterns between neighboring mRNA gene pairs (blue), neighboring lncRNA and mRNA gene pairs (red), or neighboring lncRNA and mRNA gene pairs where the mRNA gene expression was randomly shuffled (gray), with significantly enriched mRNA gene sets among neighbors of differentially expressed lncRNAs shown to the right. (**G**) Concurrent expression of endocrine development TF genes and their adjacent lncRNA. Top: normalized RNA-seq signal across the indicated lncRNA loci, with ATAC signal density shown below. Bottom: relative expression patterns and significance of correlation (t-test). (**H-I**) Alternative transcript isoform statistics (H) and distribution (I) among differential splicing, promoter, and CDS usage events. (**J-K**). Differential promoter usage of *DNTM3A* (J) and *TET1* (K). Tracks display normalized H3K27ac signal across the loci shown below, with alternative promoters highlighted and relative usage of annotated long/short isoforms across differentiation stages quantified to the right.

**Figure S4.**
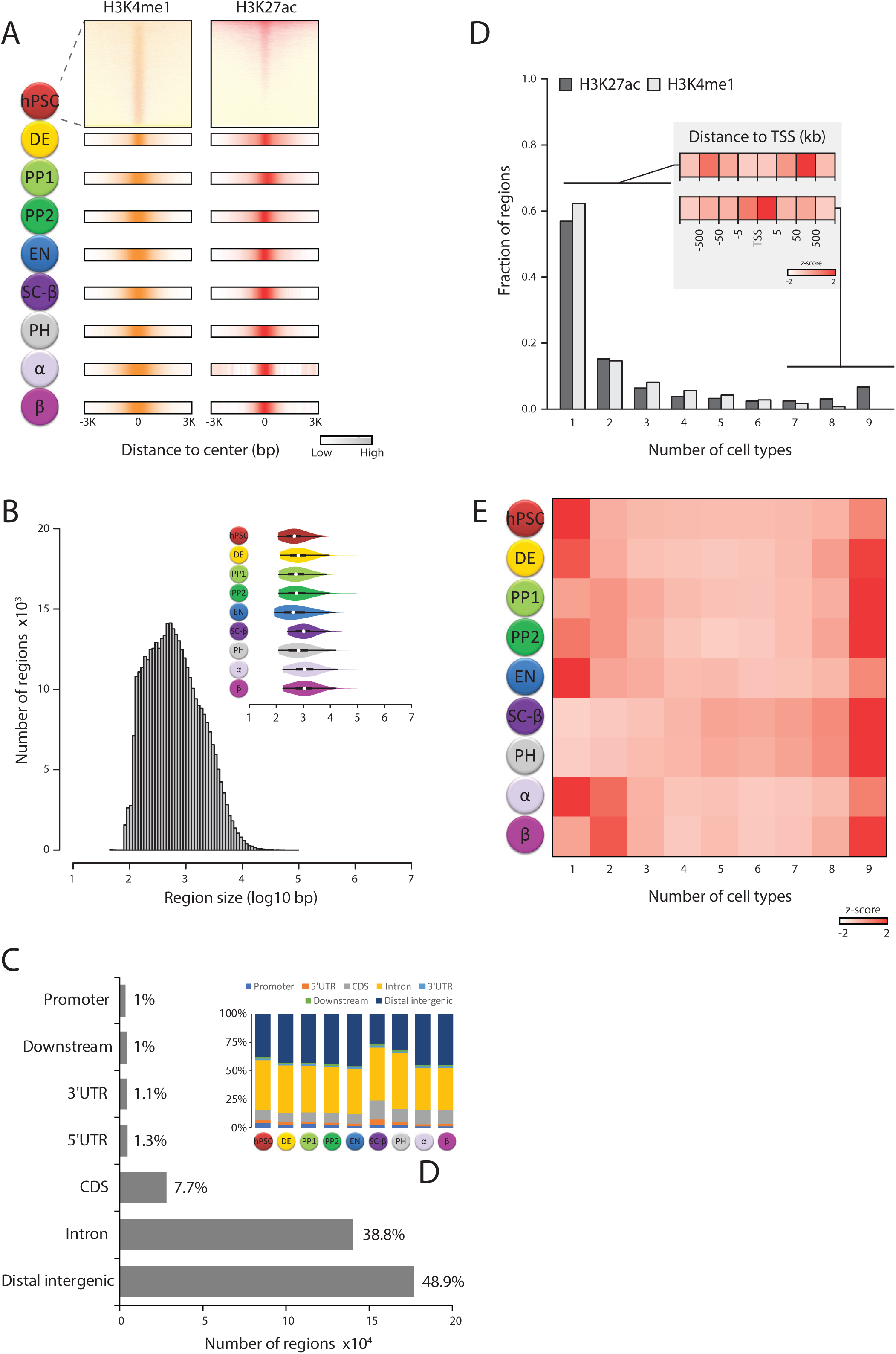
H3K4me1/H3K27ac states in the developing islet lineage, related to Figure 1. (**A**) Density of H3K4me1/H3K27ac signal enrichment over input ±3kb around the center of each region enriched for H3K27ac/H3K4me1 in at least one differentiation stage. (**B-C**) Consistent genomic coverage (**B**) and genomic location of H3K27ac/H3K4me1 enriched regions across differentiation stages. (**D-E**) High cell type-specificity of H3K27ac/H3K4me1 regions. Distribution of detection rates (D), with inset summarizing distance to the nearest TSS for regions detected in 1-3 vs. 7-9 cell types, and relative representation of regions detected in the indicated number of cell types in each differentiation stage (E)

**Figure S5.**
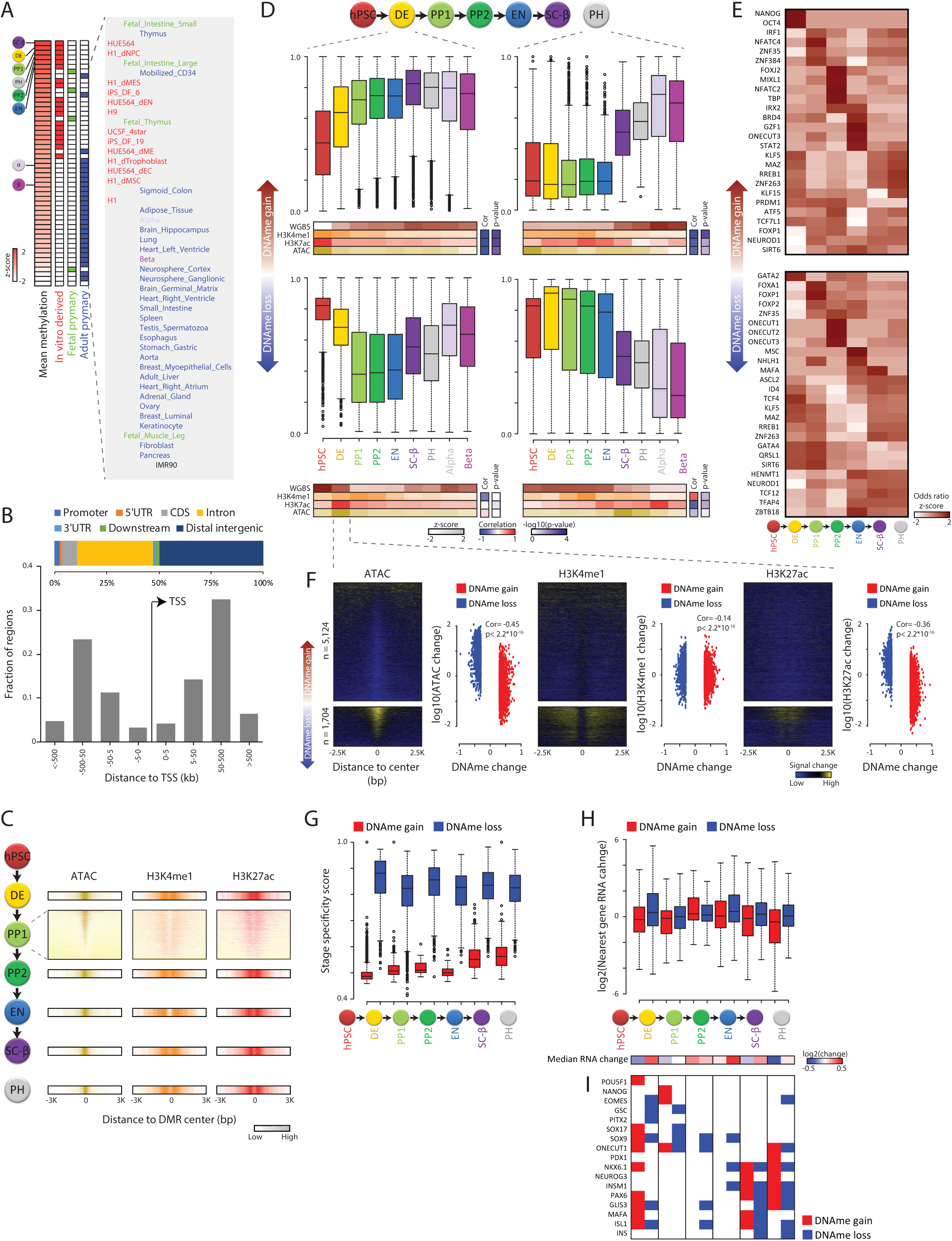
DNA methylation dynamics during islet lineage differentiation, related to Figure 1. (**A**) Hypermethylation of in vitro-derived vs. primary adult cells. Leftmost heatmap shows relative mean methylation levels across all CpGs assayed for the indicated human cell types by this study or by the NIH Roadmap Epigenomics Mapping Consortium (Roadmap Epigenomics et al., 2015). (**B-C**) DMRs comprise distal sites with focal ATAC and H3K27ac/H3K4me1 enrichment. Genomic location and distance to the nearest TSS summarized in (B), and density of ATAC, H3K4me1, and H3K27ac signal enrichment over input ±3kb around the midpoint of each region show in (C). (**D**) Patterns of epigenetic silencing upon DNAme gain (top panels) and activation upon DNAme loss (bottom panels). Boxplots show distribution of DNAme levels at all stages for regions differentially methylated during DE (left) and PH (right) specification. Heatmaps below display their median WGBS, H3K4me1, H3K27ac, and ATAC levels at each stage, with correlations between H3K4me1/H3K27ac/ATAC and WGBS patterns and their significance quantified to the right. (**E**) DMRs enrich for binding sites of known stage-specific regulators. Heatmap shows relative enrichment odds ratio of binding motifs for all expressed TFs within regions differentially methylated across stage transitions. (**F**) DNAme gain/loss coincides with focal loss/gain of ATAC, H3K4me1, and H3K27ac signal. Heatmaps show signal change ±2.5kb around the center of regions differentially methylated from hPSC to DE, quantified to the right of each heatmap. (**G**) Greater stage specificity of DNAme loss vs. gain within regions differentially methylated across stage transitions. (**H**) DNAme gain/loss within DMRs is preferentially associated with repression/induction of the nearest gene across stage transitions (top). Bottom: median of distributions summarized above. (**I**) Key stage-specific TFs and markers show DNAme loss at their respective stages. Heatmap shows DNAme change status across the stage transitions indicated in (H).

**Figure S6.**
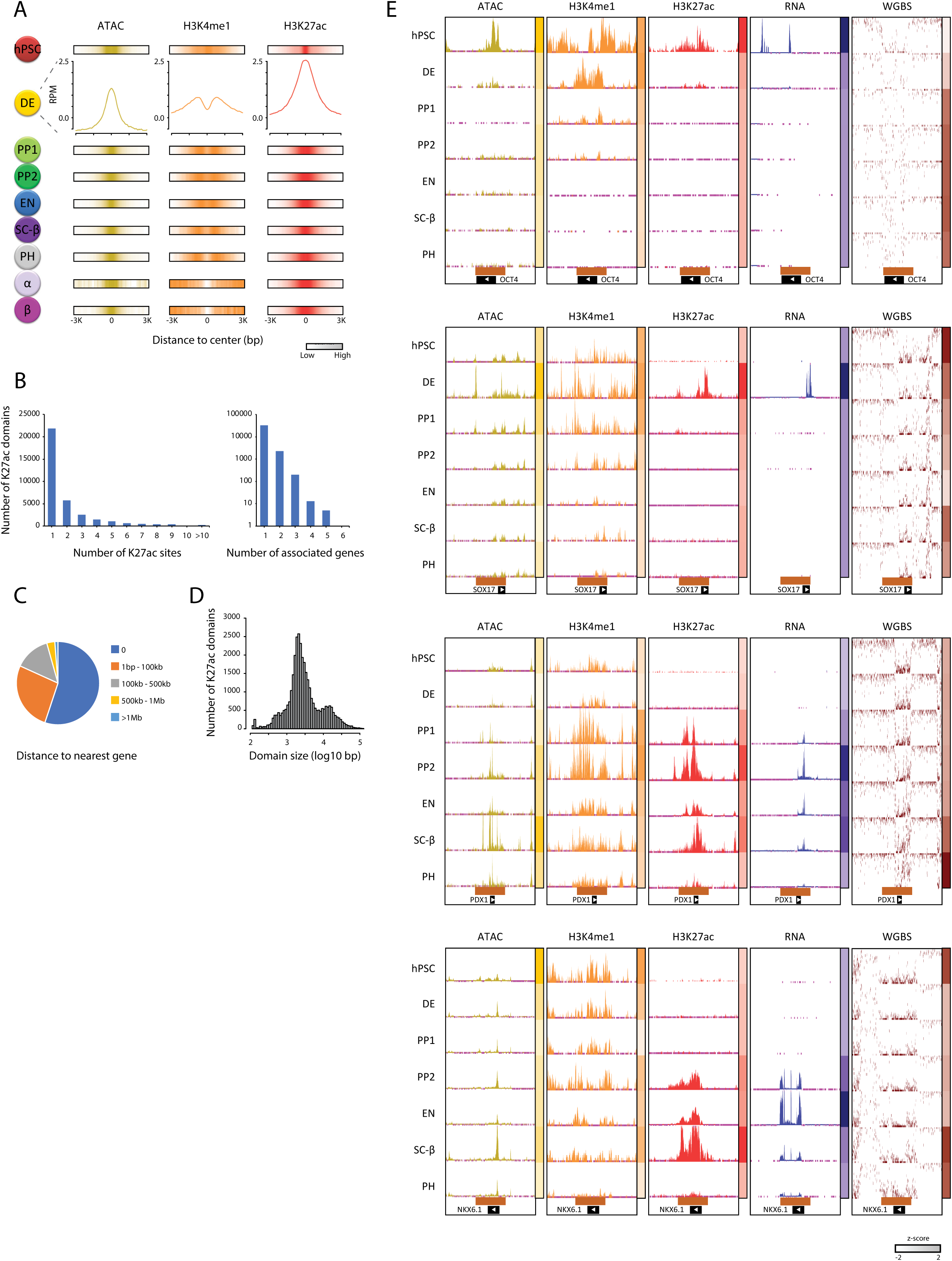
Identification of enhancers in the developing islet lineage, related to Figure 2. (**A**) H3K27ac-enriched sites outside gene TSS ±2kb regions show localized ATAC and H3K4me1 signal. Shown is the density of ATAC, H3K4me1, and H3K27ac signal enrichment over input ±3kb around the center of each site. (**B-D**) Characterization of enhancer domains defined by linking clustered enhancer sites: most (79%) comprise 1-2 enhancer sites (B left), 93% are associated with 1 gene (B right), and 55% overlap the gene that they are associated with (C). Their size distribution is summarized in (D). **(E)** Coordinated ATAC, H3K4me1, H3K27ac, RNA, and WGBS dynamics for *OCT4, SOX17, PDX1*, and *NKX6.1* along the directed differentiation timecourse. Tracks display normalized signal over a region 3x greater than the enhancer domain shown below each panel; heatmaps to the right of panels quantify relative signal over that domain.

**Figure S7.**
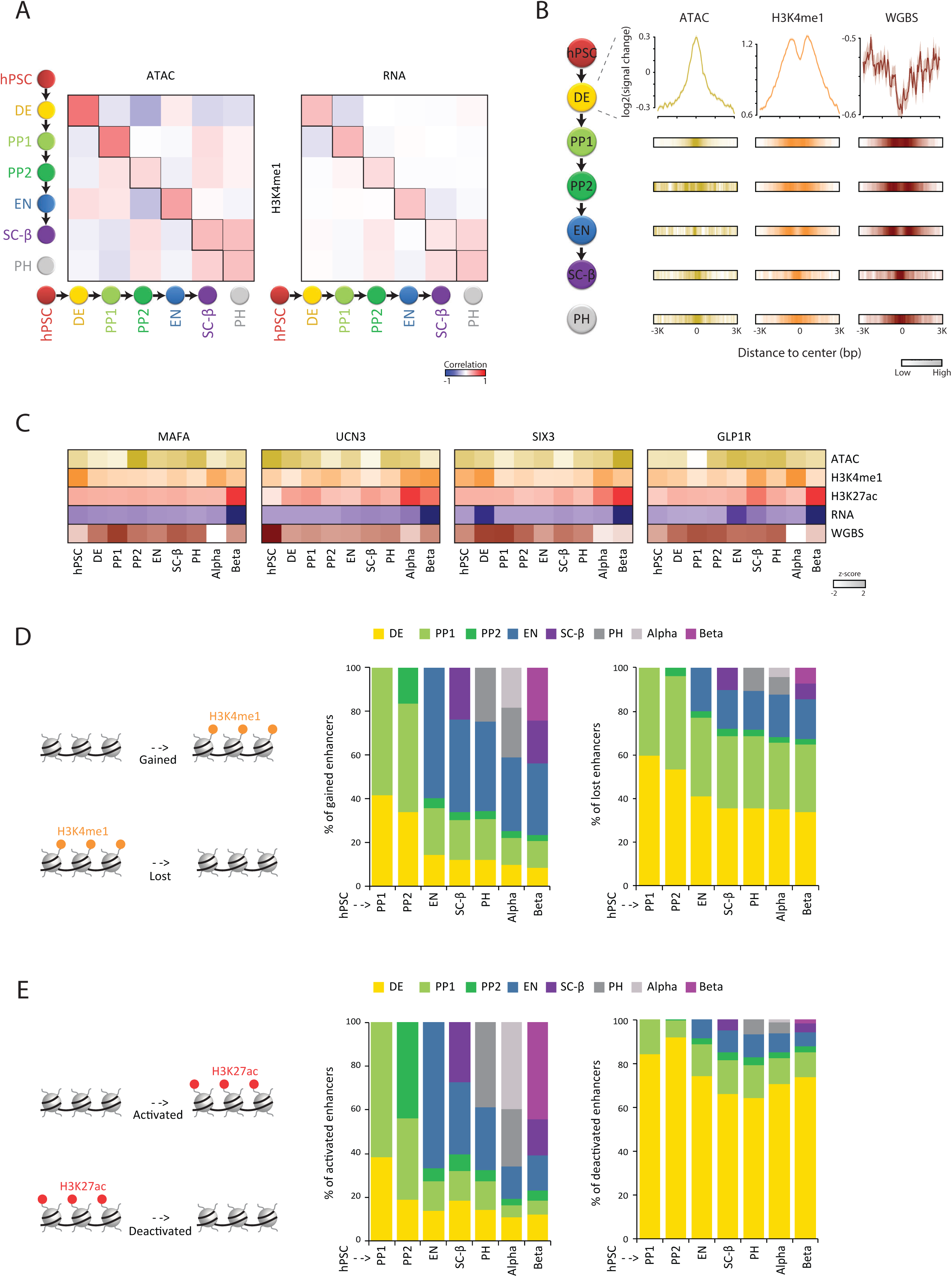
Enhancer dynamics during islet lineage progression, related to Figure 2. (**A**) Enhancer formation, denoted by H3K4me1 deposition, co-occurs with chromatin opening and RNA transcription across differentiation stages. Heatmaps show correlation between changes in H3K4me1 and changes in ATAC (left) or RNA (right) levels across all enhancer domains for each pair of differentiation stages. (**B**) Enhancer formation coincides with focal gain/loss of ATAC/WGBS signal, respectively. Shown is the change in ATAC, H3K4me1, and WGBS signal ±3kb around the center of each enhancer domain. (**C**) Epigenetic/RNA profiles for select enhancer domains, associated with the indicated genes, that are lost from hPSC and are re-formed/activated specifically in mature β cells. (**D-E**) Greater turnover of enhancer establishment/usage in the first pancreatic/endocrine committed progenitor vs. subsequent progeny. Plots show, for enhancers gained/lost (D) or activated/deactivated (E) between hPSC and each differentiation stage, the fraction gained/lost or activated/deactivated at that stage vs. previous stages, respectively.

**Figure S8.**
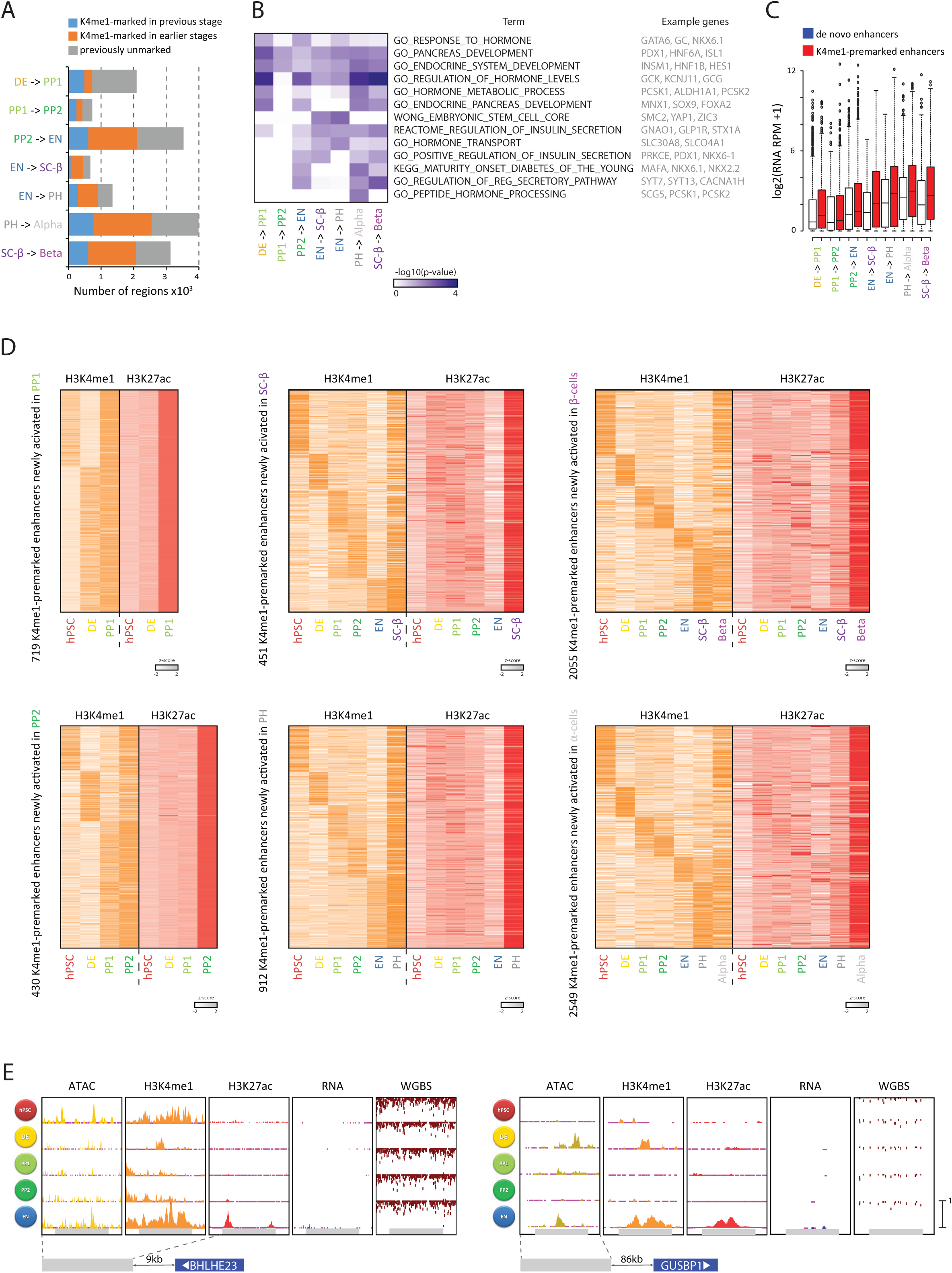
Epigenetic priming across the stages of islet lineage development, related to Figure 3. (**A**) H3K4me1 state transitions at enhancers specifically activated across stage transitions. (**B**) Enhancers that undergo primed-to-active state transitions enrich for loci encoding stage-specific regulators. Heatmap shows gene set annotations enriched at each stage transition, with member genes shown to the right. (**C**) H3K4me1-premarked enhancers gain higher RNA levels than de novo enhancers across stage transitions. (**D**) Chromatin dynamics of enhancers newly-activated in the indicated stages that were H3K4me1-marked in previous stages. Heatmaps show enhancer domain H3K4me1/H3K27ac signal sorted by H3K4me1 gaining step. (**E**) Transient H3K4me1 marking of enhancers neighboring *BHLHE23* (left) and *GUSBP1* (right) that are newly activated in EN. Tracks display normalized signal over a region 1.5x greater than the enhancer domain shown below each panel.

**Figure S9.**
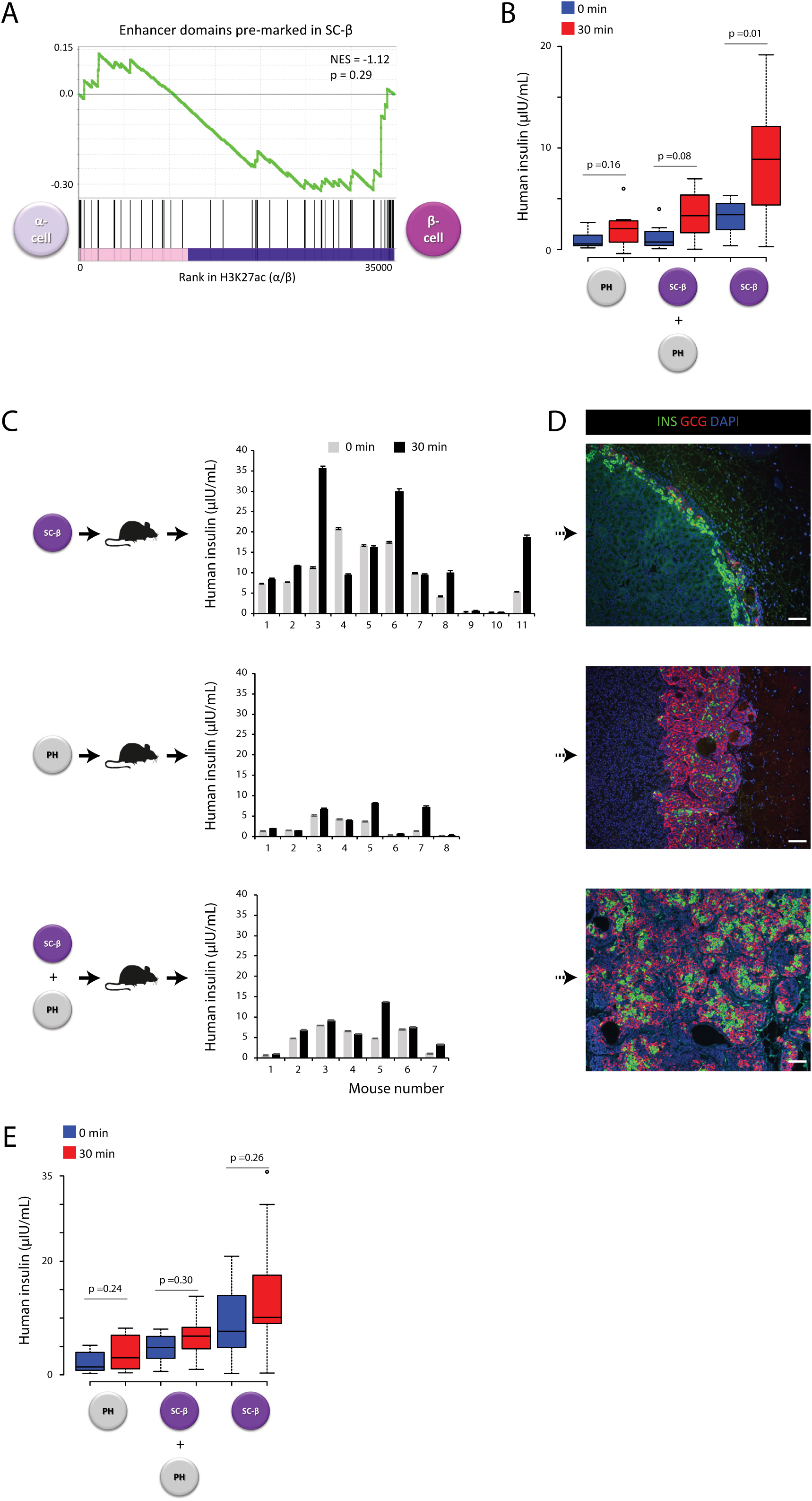
Epigenetic priming identifies polyhormonal cells as α-cell precursors, related to Figure 3. (**A**) Biased priming of β-cell-specific enhancers in SC-β cells. Enrichment analysis of enhancers inactive but H3K4me1-marked in SC-β by their H3K27ac level in α vs. β cells. NES (normalized enrichment score). (**B-C**) Human insulin levels in the serum of fasted mice transplanted with FACS-purified PH, SC-β, or unsorted re-aggregated cells assayed before and 30min after a glucose injection 4-6 weeks post-transplantation (B) and at the end of the post-transplantation observation period (>2 months) (C). (**D**) Additional examples of hormonal resolution evidenced by staining retrieved grafts for insulin (green)/glucagon (red). Scale bar=100μm. (**E**) Data from (C) quantified.

**Figure S10.**
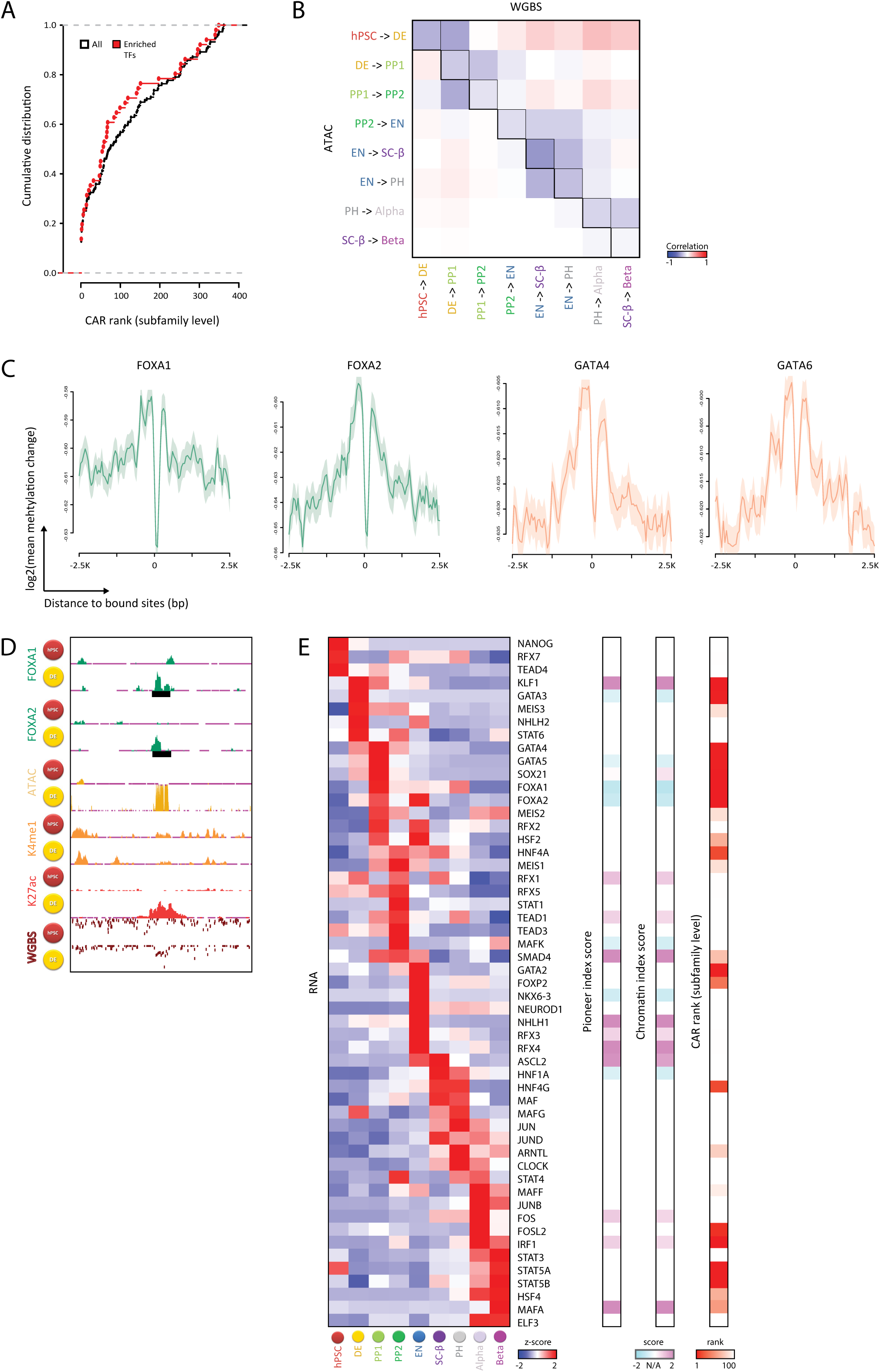
Pioneer factors across the islet differentiation path, related to Figure 4. (**A**) Newly-opened chromatin sites enrich for binding motifs of TFs with strong pioneer activity. Cumulative distribution of subfamily level chromatin accessibility regulator (CAR) ranks from (Lamparter et al., 2017), for all TF motifs analyzed and for TF motifs enriched at chromatin sites that become newly-opened across developmental transitions. (**B**) Increased chromatin openness co-occurs with DNAme loss. Heatmap shows correlation between changes in ATAC and changes in WGBS-based DNAme levels across all enhancer domains for each pair of differentiation stages. (**C**) Localized demethylation at sites bound by known pioneers FOXA1, FOXA2, GATA4, GATA6 in DE (Tsankov et al., 2015). Plots show relative decrease in mean CpG methylation level between hPSC and DE ±2.5kb from the center of each TF peak. (**D**) Example of localized demethylation at the binding sites of known pioneer factors FOXA1/2 in DE (Tsankov et al., 2015). Tracks display normalized signal (y-axis scale: 0-1) over a region 10x greater than the FOXA1/2 DE binding peak (shown in black). (**E**) Stage-specific expression patterns for select TFs enriched at chromatin sites newly opened across differentiation stages, with pioneer index and chromatin opening index log-odds scores from (Sherwood et al., 2014), as well as CAR ranks from (Lamparter et al., 2017) shown to the right.

**Figure S11.**
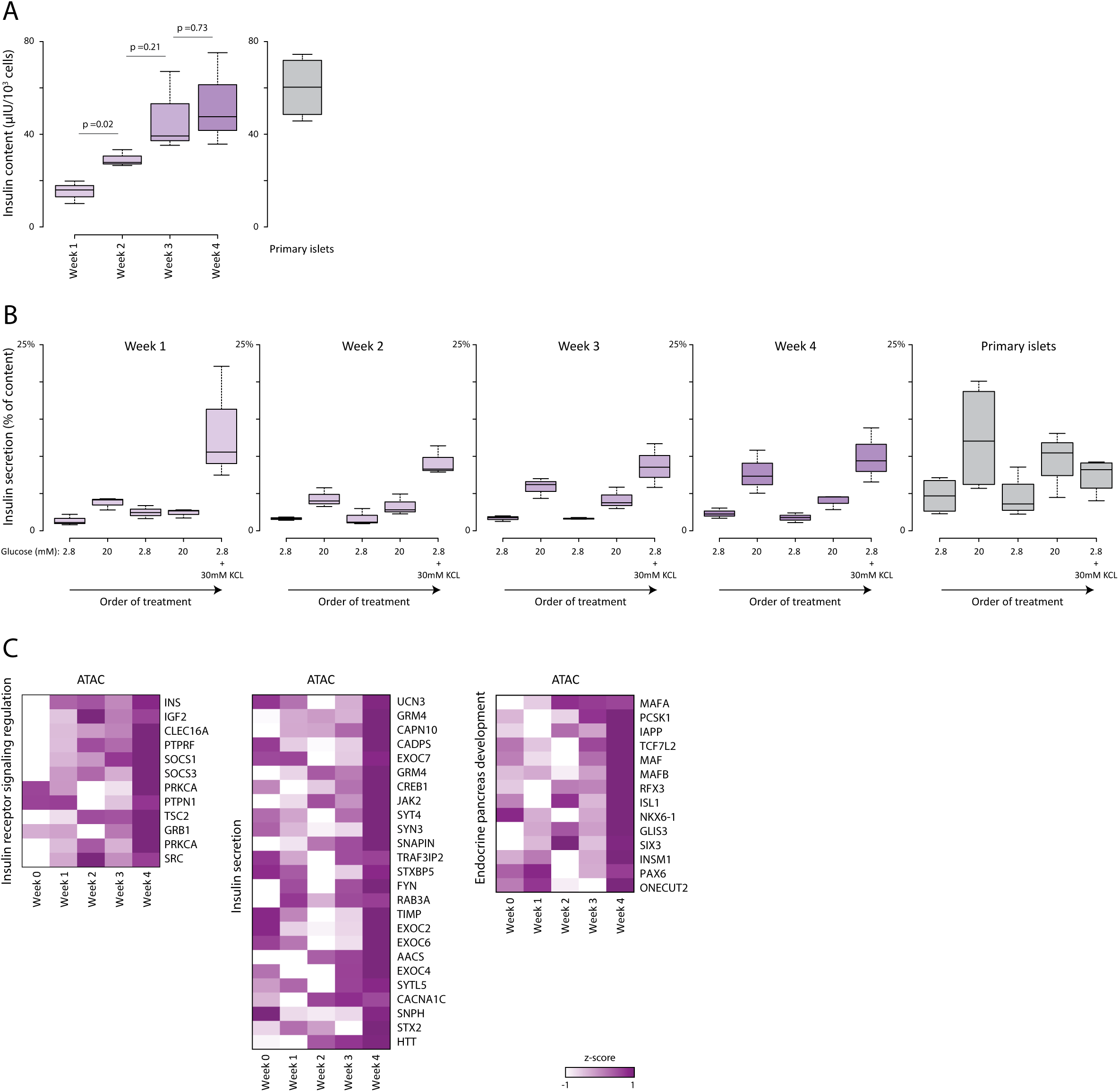
Refinement of GSIS function during extended culture of terminal-stage SC-β preparations, related to Figure 4. (**A-B**) Gradual increase of insulin production and secretory capacity. Boxplots summarize insulin content (A) and secretion over the indicated sequential 30min incubations (B), compared to cadaveric islets assayed in parallel. Data are pooled from N=3 SC-β and N=4 cadaveric islet preparations with n=3 replicate measurements each. (**C**) Gradual chromatin opening at genes regulating insulin processing and secretion. Heatmaps quantify relative ATAC signal over enhancer domains associated with the indicated genes across a 4-week timecourse.

**Figure S12.**
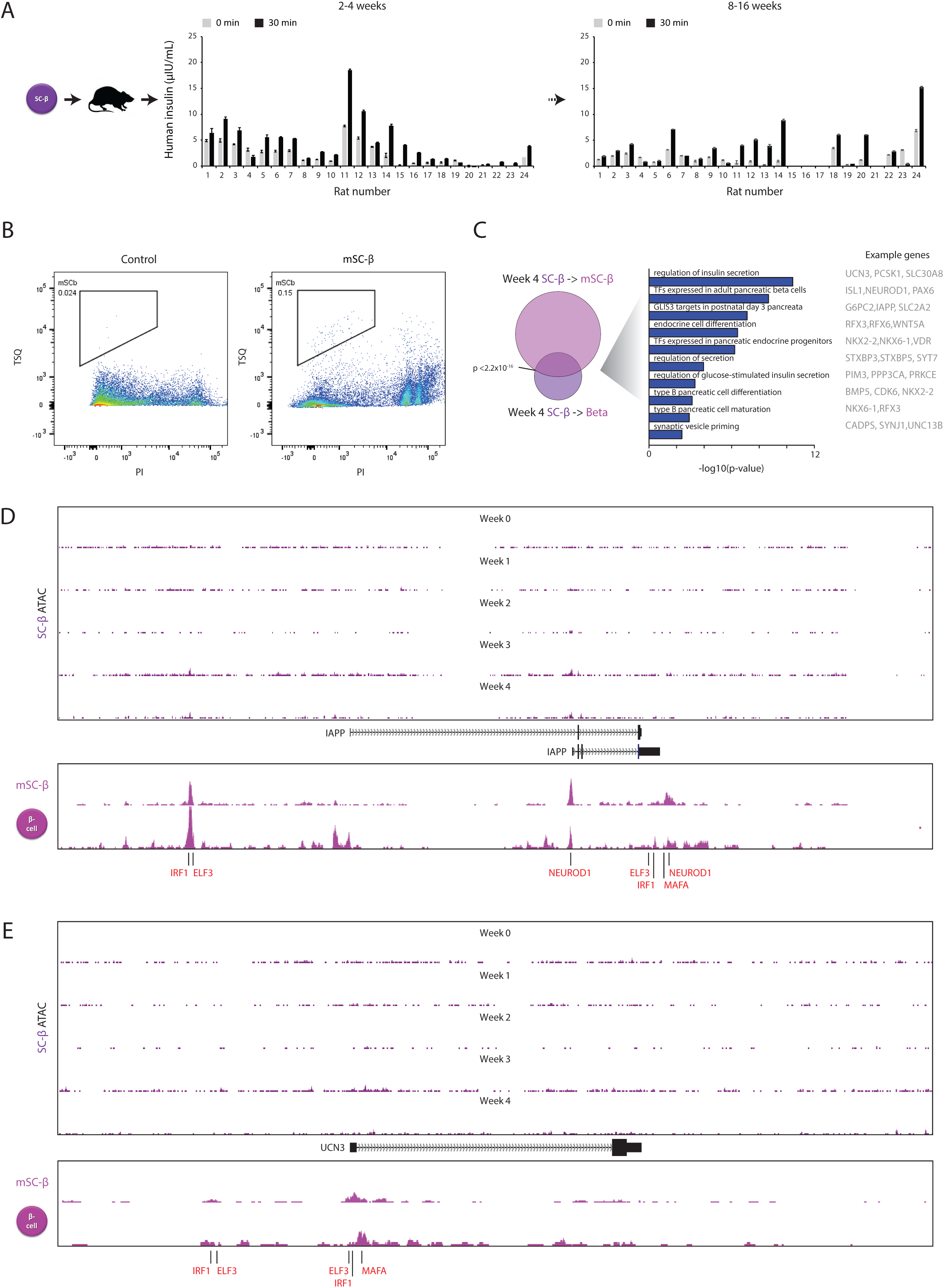
Chromatin accessibility dynamics during in vivo SC-β maturation, related to Figure 4. (**A**) Human insulin levels in the serum of fasted rats transplanted with preparations containing 5×10^7^ SC-β cells were assayed before and 30min after a glucose injection 2-4 weeks post-transplantation (left) and at the end of the post-transplantation observation period (2-4 months) (right). (**B**) Sorting strategy for isolating in vivo-matured SC-β from retrieved grafts. (**C**) Chromatin sites newly opened in transplanted SC-β enrich for maturation-associated TF and secretory genes. Overlap of sites newly opened in mSC-β vs. primary β relative to week-4 SC-β (left), with significantly enriched gene sets shown to the right. (**D**-**E**) Chromatin accessibility landscapes around *IAPP* (D) and *UCN3* (E) at the indicated stages. Tracks display normalized ATAC signal over the regions shown below, with select TF motifs at chromatin regions newly-accessible in mSC-β/primary β highlighted in red.

**Figure S13.**
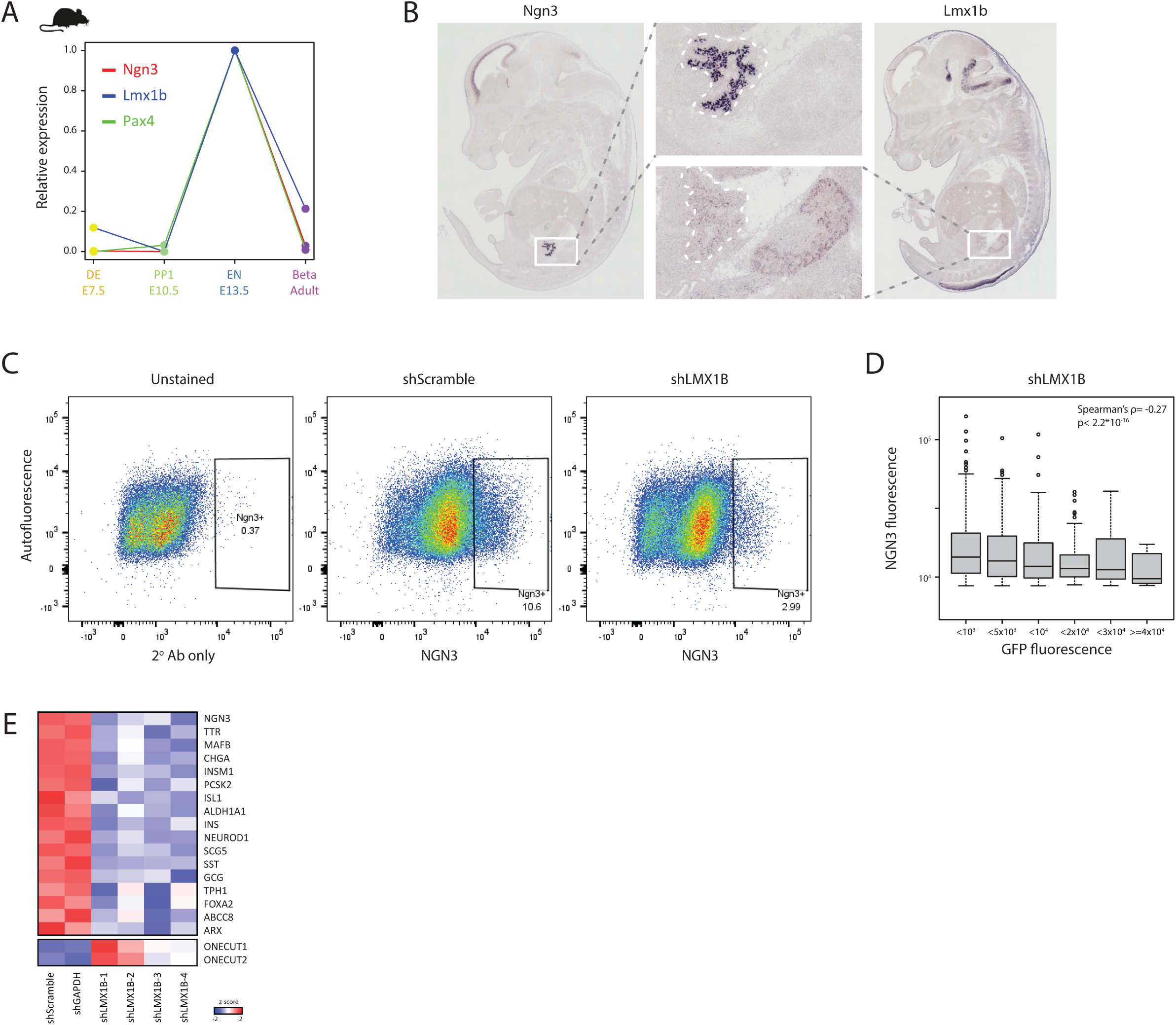
Identification of super-enhancers in the developing islet lineage, related to Figure 5. (**A**) Distribution of enhancer domain H3K27ac signal across differentiation stages identifies super-enhancers with exponentially higher levels than the rest, comprising lineage-and stage-specific regulators. (**B**) Top pathways enriched among genes associated with super-enhancers. Colors correspond to the stage of differentiation as shown in A. (**C**) Stage-specific H3K27ac activity at promoters of typical enhancer-and super enhancer-associated genes. Boxplots quantify stage specificity by the fractional promoter H3K27ac level across differentiation stages.

**Figure S14.**
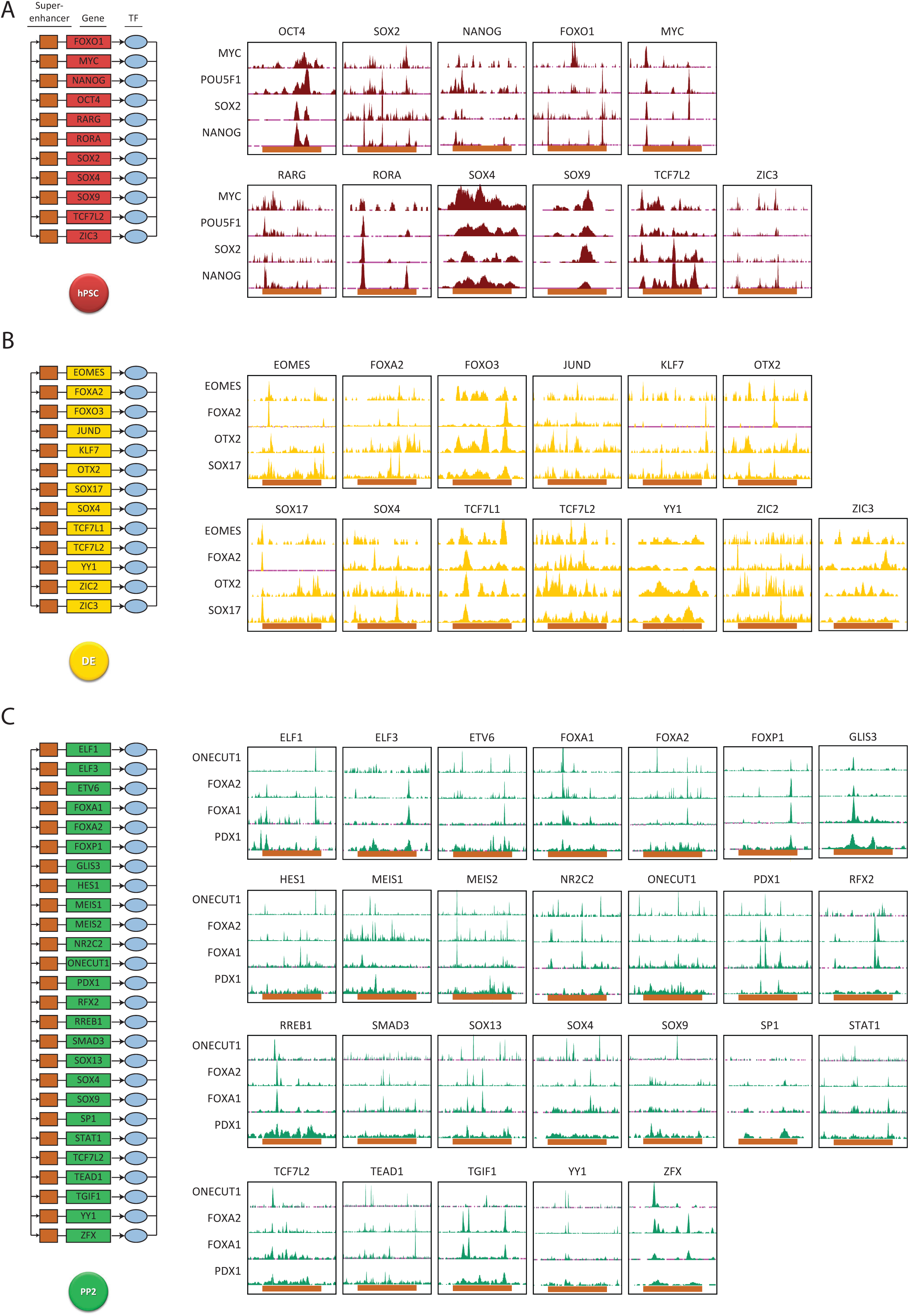
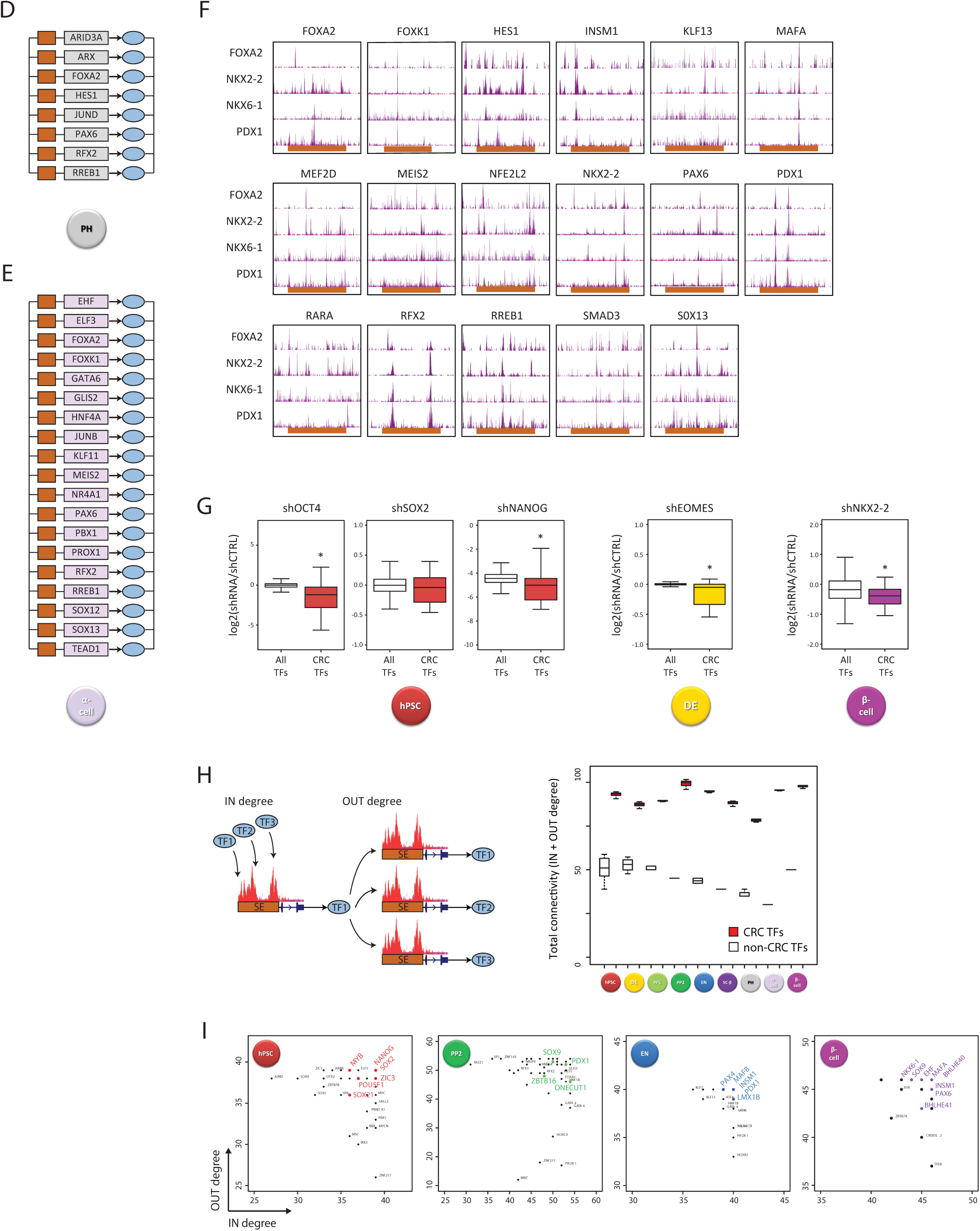
Identification of core regulatory circuits across the stages of islet lineage development, related to Figure 5. (**A-F**) Representative CRC models for the indicated stages. Panels show ChIP-seq for MYC/OCT4/SOX2/NANOG in hPSC (A), EOMES/FOXA2/OTX2/SOX17 in DE (Tsankov et al., 2015) (B), ONECUT1/FOXA2/FOXA1/PDX1 in PP2 (Cebola et al., 2015) (C), and FOXA2/NKX2-2/NKX6-1/PDX1 in cadaveric islets (Pasquali et al., 2014) (F) to the right of the hPSC, DE, PP2, and PH/α CRCs, respectively. Tracks display ChIP-seq signal over a region 1.5x greater than the SE domain shown below for each CRC TF. (**G**) Effect of shRNA-mediated depletion of the indicated factors in hPSC (Wang et al., 2012b), DE (Teo et al., 2011), and cadaveric islets (Gutierrez et al., 2017) on the respective CRC TFs, compared to the full set of TFs expressed at each stage. Boxplots display fold change in expression relative to control. *p <0.05, Wilcoxon test. (**H**) Interconnectivity among CRC vs. non-CRC TFs. The strategy for quantifying inward regulatory degree (IN degree) and outward regulatory (OUT degree) via motif binding is shown to the left. Boxplots to the right show total connectivity (IN + OUT degree) for CRC-forming vs. non-CRC-forming SE-driven TFs at each differentiation stage. (**I**) Scatter plots showing IN and OUT regulatory degree, as defined in (H), for CRC-forming vs. non-CRC-forming SE-driven TFs at the indicated stages.

**Figure S15.**
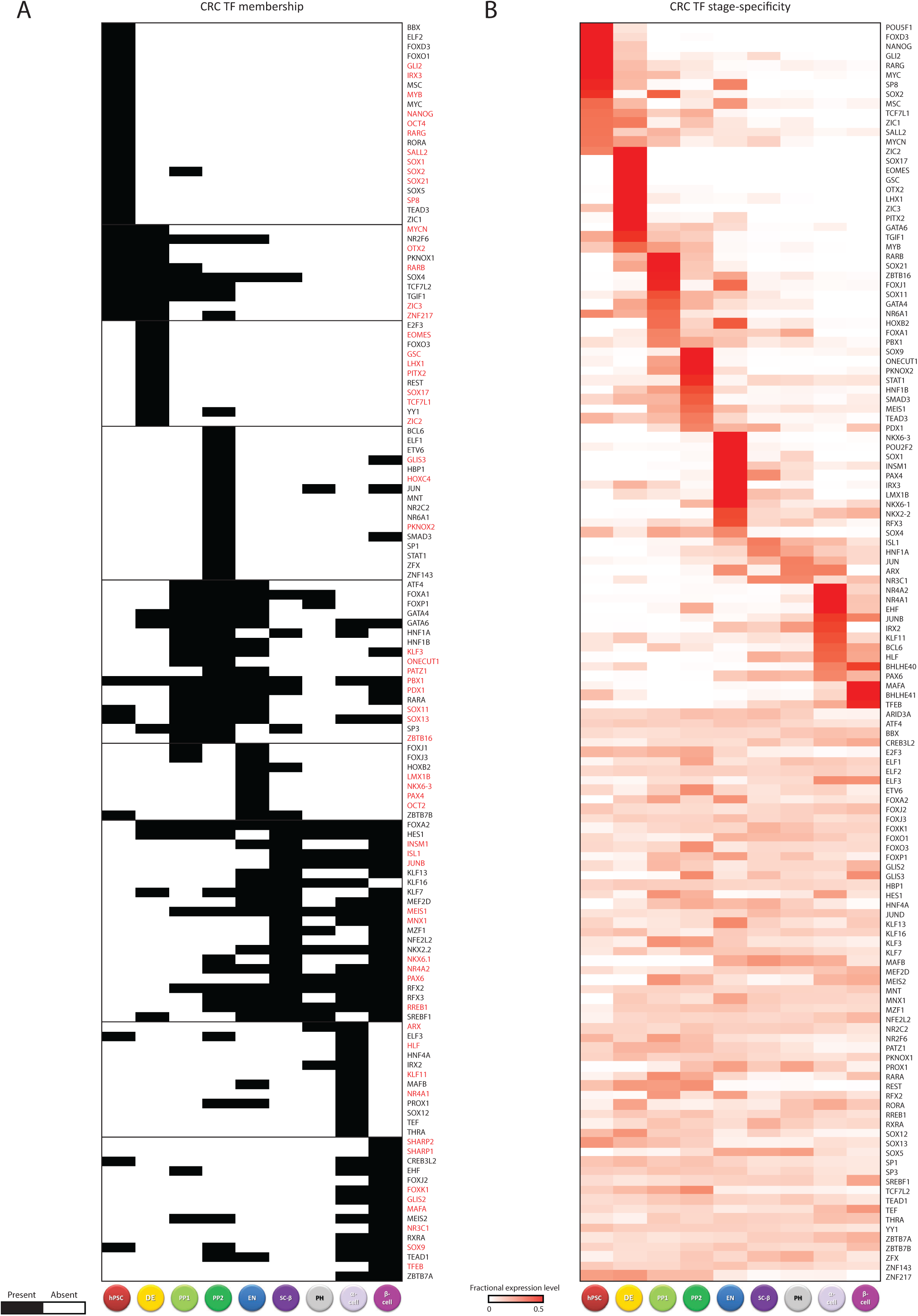
CRC TFs across the islet differentiation path, related to Figure 5. (**A**) K-means clustering (K=9) of CRC TF membership across pancreatic differentiation stages. Stage-specific factors are highlighted in red. (**B**) Stage specificity of CRC TFs in the human islet lineage.

**Figure S16.**
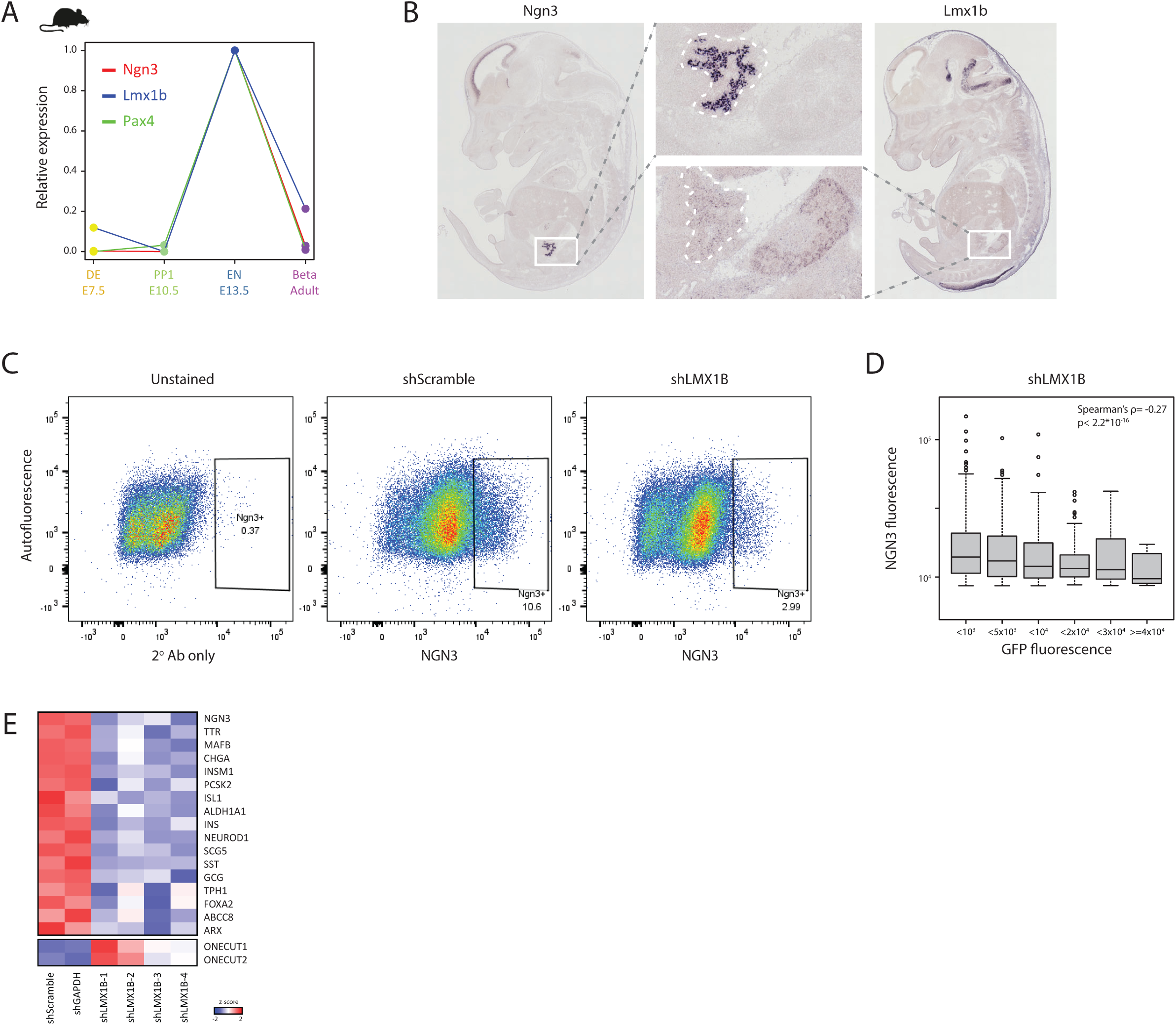
LMX1B is a critical regulator of human endocrine progenitors, related to Figure 5. (**A-B**) Lmx1b is specifically active in Ngn3+ progenitors in vivo. Lmx1b expression in the mouse pancreatic islet lineage follows that of endocrine regulators Ngn3 and Pax4 (Gu et al., 2004) (A). In situ detection of Ngn3 and Lmx1b mRNA in separate mouse E14.5 sections (Richardson et al., 2014) (B). Insets highlight embryonic pancreata. (**C-D**) *LMX1B* depletion triggers proportional NGN3 depletion. Quantification of NGN3 protein in fixed EN cells transduced with a GFP-expressing *LMX1B*-targeted shRNA vector by intracellular immunostaining (C), and correlation between NGN3 and GFP levels (D). (**E**) Reciprocal downregulation of endocrine lineage markers and upregulation of pancreatic progenitor markers upon LMX1B knockdown in sorted EN shRNA-transduced cells.

**Figure S17.**
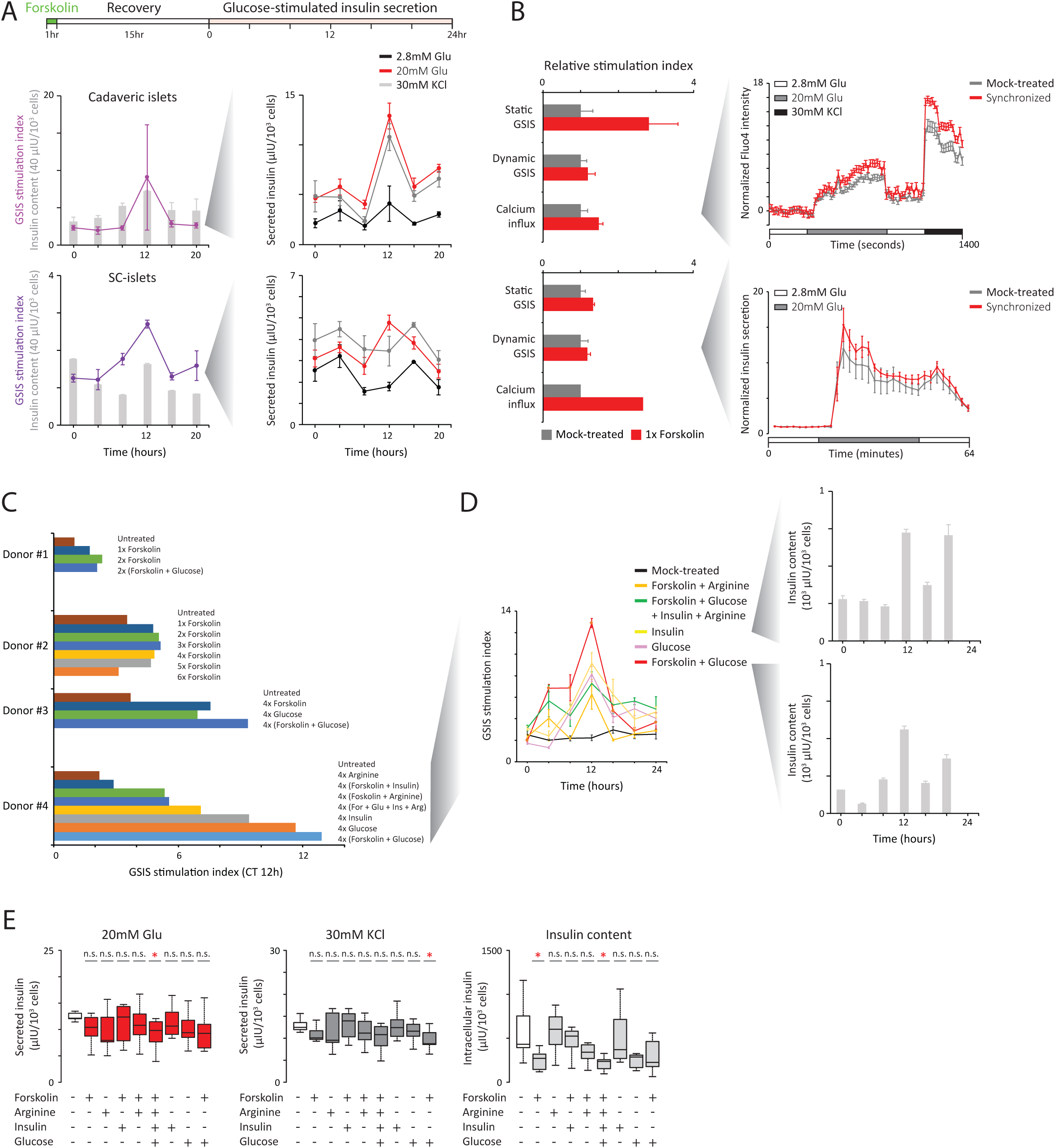
Circadian rhythms trigger an islet maturation step, related to Figure 6. (**A**) Rhythmic insulin production/responses in cadaveric/SC-isles synchronized by forskolin. Schematic: timeline for forskolin shock and recovery followed by functional assays. Left: GSIS stimulation indexes and insulin content across 20h for synchronized cadaveric (top) and SC-islets (bottom). Right: insulin secretion over the indicated sequential 30min incubations assayed every 4h for 20h. (**B**) Synchronization enhances glucose responsiveness. Shown are relative stimulation indexes for synchronized vs. mock-treated cadaveric (top) and SC-islets (bottom), as determined by static/dynamic GSIS or calcium influx assays at the 12h circadian timepoint. Representative calcium signaling / insulin release dynamics during the indicated sequential incubations are shown to the right. Data are mean of n=7 islets (calcium assay) or n=3 sets of islets (dynamic GSIS) sampled from each culture, normalized to mean of the first incubation. (**C-D**) Recurring shocks of various stimuli can further amplify islet GSIS function. Stimulation indexes determined by static GSIS assays at the 12h circadian timepoint following the indicated treatments for four cadaveric donor batches (C). Indexes across 24h for the indicated experiment are shown (D), with 24h insulin content profiles for select treatments shown to the right. (**E**) Insulin responses to stimulatory glucose/KCL incubations and insulin content for the cultures shown in (D). Boxplots summarize insulin secretion/content across 24h for the indicated conditions; *p <0.05 relative to mock-treated (t-test).

**Figure S18.**
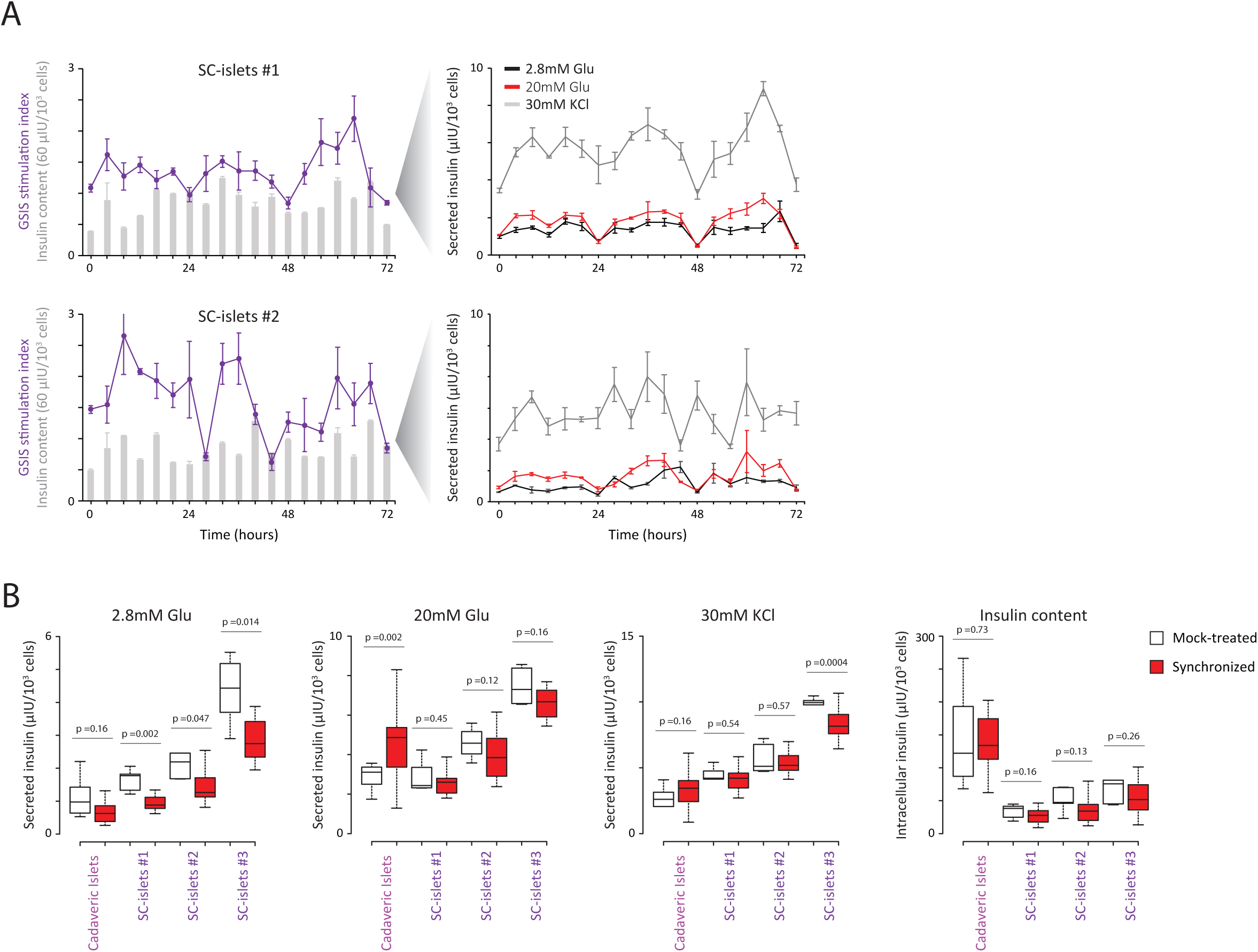
Circadian rhythms trigger an SC-islet maturation step, related to Figure 6. (**A**) Additional examples of rhythmic GSIS responses in synchronized SC-islet cultures across 72h. Left: GSIS stimulation indexes and insulin content profiles. Right: insulin secretion over the indicated sequential 30min incubations assayed every 4h for 72h. (**B**) Synchronized SC-islet cultures show decreased insulin responses to low (2.8mM) glucose incubations without significant change in overall insulin secretion under stimulatory (20mM glucose/30mM KCL) incubations or overall insulin content. Boxplots summarize insulin secretion/content across 72h.

**Figure S19.**
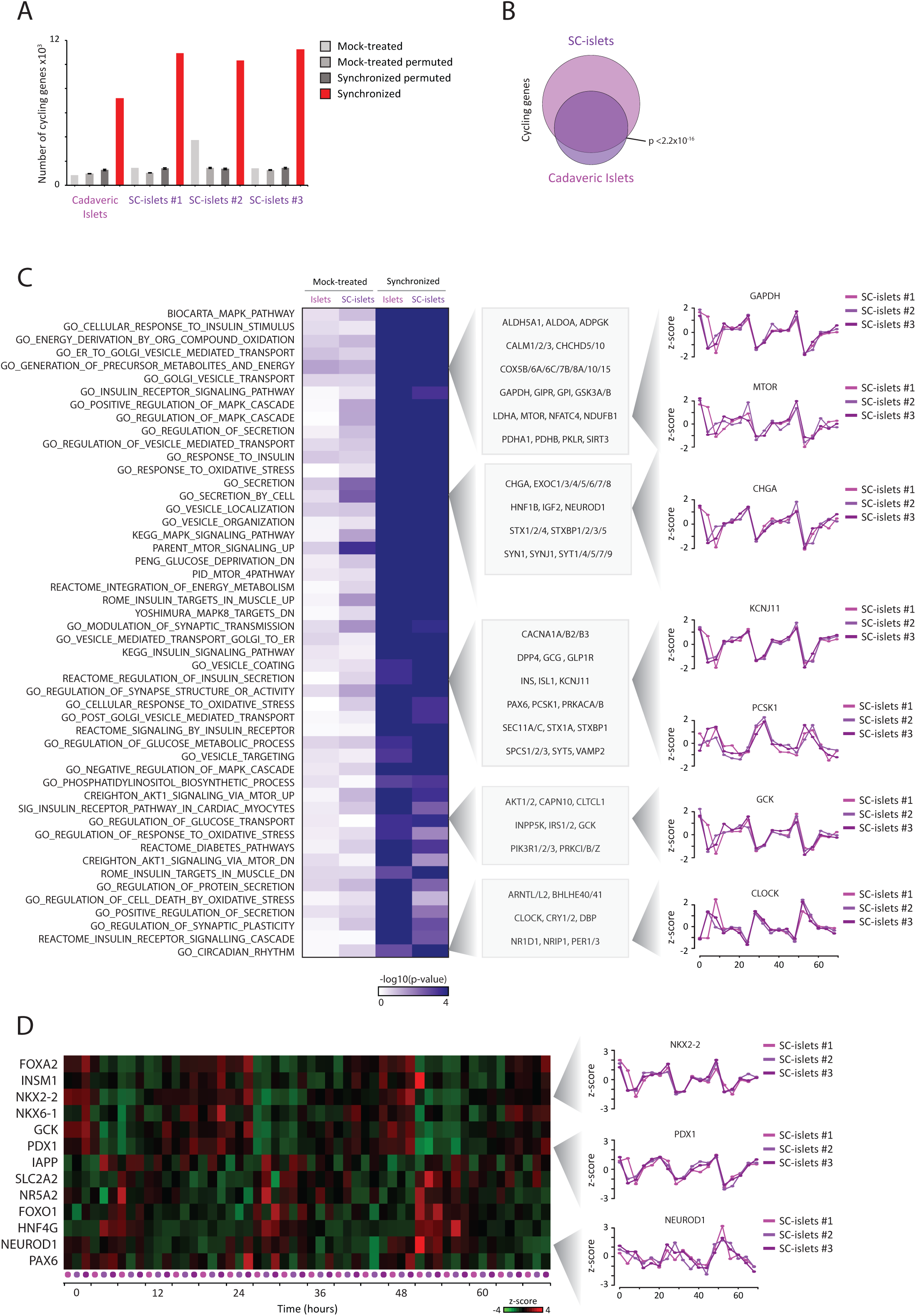
Transcriptional basis for coordinated metabolic and insulin cycles, related to Figure 6. (**A**) Identification of genes specifically induced to cycle upon synchronization of cadaveric/SC-islets. Shown is the number of cycling genes (q <0.05, harmonic regression test for rhythmicity) in synchronized vs. mock-treated cultures based on their ordered or randomly-permuted RNA-seq expression profiles. (**B**) Overlap between genes cycling in cadaveric and SC-islet cultures. (**C**) Selective enrichment of energy metabolism effectors and regulators as well as insulin processing/transport/secretion and signaling factors among cycling gene sets. Heatmap shows gene set enrichment p-values for mock-treated vs. synchronized cadaveric/SC-islet cultures, with profiles for select rhythmic mRNAs shown to the right. (**D**) Expression patterns for select pancreatic TFs and markers, with cycling profiles shown to the right.

## EXTENDED EXPERIMENTAL PROCEDURES

### Cell culture

The hPSC line HUES8 (NIH human embryonic stem cell registry #0021) was used for all experiments. Undifferentiated HUES8 cells were maintained in supplemented mTeSR1 medium (StemCell Technologies) in spinner flasks (Corning) set at a 70rpm rotation rate in a 37°C 5% CO_2_ incubator. Directed differentiation into islet cells was conducted as described previously (Pagliuca et al., 2014) with the modifications described in (Millman et al., 2016). Briefly, 150million cells were seeded in 300ml mTeSR1 +10μM ROCK Inhibitor Y27632, fed with mTeSR1 48h later, and 72h later stepwise differentiation stages were induced by the following treatments:

Stage 1: 24h in S1 medium +100ng/ml ActivinA +14μg/ml CHIR99021, followed by 48h in the same medium without CHIR99021.

Stage 2: 72h in S2 medium +50ng/ml KGF.

Stage 3: 24h in S3 medium +50ng/ml KGF +0.25μM Sant1 +2μM Retinoic acid (RA) +500nM PDBU +10μM Y27632 +200nM LDN193189, followed by 24h in the same medium without LDN193189.

Stage 4: 6 days in S3 medium +50ng/ml KGF +0.25uM Sant1 +0.1μM Retinoic acid (RA) +10μM Y27632 +5ng/ml ActivinA.

Stage 5: 4 days in BE5 medium +0.25μM Sant1 +0.1μM Retinoic acid (RA) +1μM XXI +10μM Alk5i II +1μM T3 +20ng/ml Betacellulin +10μM Y27632, followed by 3 days in BE5 medium +25nM Retinoic acid (RA) +1μM XXI +10μM Alk5i II +1μM T3 +20ng/ml Betacellulin +10μM Y27632.

Stage 6: 7-30 days in supplemented CRML-1066 medium +10% defined fetal bovine serum (FBS, Hyclone) +10μM Alk5i II +1μM T3.

Primary adult islets from cadaveric donors (Prodo laboratories) were cultured in supplemented CRML-1066 medium +10% FBS in low-attachment plates (Corning). All methods involving human cells were approved by the Harvard University IRB and ESCRO committees.

### Cell purification

Cell clusters sampled upon completion of each differentiation stage or primary islets were washed with PBS, incubated in PBS +50% Accutase (StemCell Technologies) for 7min, and dissociated mechanically by petting up and down in PBS +10μM Y27632. For fixed-cell purification, cells were cross-linked in PBS +1% paraformaldehyde for 10min, quenched with 125mM glycine for 5min, and washed with PBS. Next, cells were blocked in PBS +5% donkey serum (Jackson Immunoresearch) +0.1% saponin (Sigma) for 20min and subsequently stained with primary antibodies in blocking buffer for 1hr and fluorophore-conjugated secondary antibodies in blocking buffer for 30min. Stained cells were washed with and resuspended in PBS +5% donkey serum. For live-cell purification, single-cell suspensions were stained in PBS + primary fluorophore-conjugated antibodies for 30min on ice, washed with and resuspended in PBS. Stained fixed/live cells were filtered through a 40μm nylon mash into flow cytometry tubes (BD Falcon) and sorted using MoFlo flow cytometers (Beckman Coulter) into PBS +1% BSA (Sigma) on ice. The cell populations as defined by antibody combinations were:

Stage 1: SOX17+ (R& D Systems; AF1924)

Stage 3: PDX1+ (R& D Systems; AF2419)

Stage 4: PDX1+ NKX6+ (DSHB; F55A12) for fixed cells, and GP2+ (MBL International; D277-5) (Ameri et al., 2017; Cogger et al., 2017) for live cells.

Stage 5: NGN3+ (R& D Systems; AF3444) for fixed cells, and SUSD2+ (Milentyi Biotec; 130-106-401) (Liu et al., 2014) for live cells.

Stage 6 SC-β: C-peptide+ (DSHB; GN-ID4) Glucagon-like peptide-2 (GLP2)-(Santa Cruz; sc-7781) for fixed cells, and TSQ+ (Thermo Fisher Scientific; M688) DPP4-(Miltenyi Biotec; 130-093-441) for live cells.

Stage 6 PH: C-peptide+ GLP2+ for fixed cells, and TSQ+ DPP4+ for live cells.

Primary β cells: C-peptide+ GLP2-for fixed cells, and TSQ+ HPi2+ (Novus Biologicals; NBP1-18946AF488) TM4SF4-(Novus Biologicals; FAB7998R) (Muraro et al., 2016) for live cells.

Primary α cells: C-peptide+ GLP2+ for fixed cells, and TSQ+ HPi2+ (Novus Biologicals; NBP1-18946AF488) TM4SF4+ (Novus Biologicals; FAB7998R) (Muraro et al., 2016) for live cells.

Live cells were defined as Propidium Iodide (PI)-negative cells. Sorting strategies for each subpopulation are summarized in Figure S1.

We validated our strategy for sorting live SC-β from PH using DPP4, which inactivates glucagon-like peptide-1 (GLP1) (Mentlein et al., 1993), in conjugation with the TSQ zinc dye, by fixing sorted TSQ+ DPP4-and TSQ+ DPP4+ cells and verifying >90% C-peptide+ GLP2-and >85% C-peptide+ GLP2+ enrichment, respectively, via flow cytometry-quantified intracellular staining.

### ChIP-seq

FACS-purified fixed cells (typically ∼1million) were pelleted and flash-frozen. ChIP-seq was conducted as described previously (Gifford et al., 2013) with minor modifications. Briefly, cell pellets were thawed on ice for 30min, incubated in lysis buffer (0.5% NP-40 +85mM KCl +20mM Tris-HCl pH8.0 +protease inhibitor) for 10min on ice, and nuclei were pelleted and incubated in lysis buffer (1% NP-40 +0.5% sodium deoxycholate +0.1% SDS +10mM Tris-HCl pH7.5 +protease inhibitor) for 10min on ice. Chromatin was then sheared with a Branson Sonifier (model S-450D) at 4°C and incubated with 1μg/million cells H3K27Ac (Active Motif; 39133) or H3K4me1 (Millipore, 17-614) antibody overnight at 4°C. Next, antibody-protein complexes were isolated by incubation with Protein A/G beads (Life Technologies; 100-02D/100-07D) for 2h at 4°C. Samples were then sequentially washed twice with low-salt buffer (0.1% SDS +1% Triton X-100 +2mM EDTA +20mM Tris-HCl pH8.1 +150mM NaCl), twice with high-salt buffer (0.1% SDS +1% Triton X-100 +2mM EDTA +20mM Tris), twice with LiCl buffer (0.25M LiCl +1% NP-40 +1% deoxycholate +1mM EDTA +10mM Tris-HCl pH8.1), twice with TE (10mM Tris-HCl pH8.0 +1mM EDTA), and finally eluted in freshly-prepared elution buffer (1% SDS +0.1M NaHCO_3_) at 65°C for 30min. Eluates were then treated with reverse crosslinking salt mixture (250mM Tris-HCl pH6.5 +62.5mM EDTA pH8.0 +1.25M NaCl +5mg/ml Proteinase K + 62.5ng/μl RNase A) overnight at 65°C. DNA was then purified using AMPure XP magnetic beads (Beckman Coulter), and sequencing libraries were generated using the NEBNext Ultra II DNA Library Prep Kit (New England Biolabs; E7103), pooled, and sequenced on a HiSeq 2500 instrument (Illumina).

### ATAC-seq

FACS-purified live cells (typically ∼50,000) were pelleted and tagmentation was performed at 37°C for 30min as previously described (Buenrostro et al., 2013). DNA was isolated using the MinElute PCR purification kit (Qiagen), PCR-amplified for 10 cycles, and purified using AMPure XP magnetic beads (Beckman Coulter). Double-sided AMPure cleanup to remove high-molecular-weight fragments was conducted by incubation of double-concentrated AMPure beads added at 0.55x volume to the PCR reactions, followed by cleanup with a 1x AMPure volume. Sequencing libraries were pooled and sequenced on a HiSeq 2500 instrument.

### WGBS

FACS-purified live cells (typically ∼500,000) were pelleted and WGBS was performed as previously described (Donaghey et al., 2018). DNA was fragmented with a Covaris S2 sonicator for 6min, purified using the DNA Clean and Concentrator kit (Zymo Research), and bisulfite conversion was performed using the EZ DNA Methylation-Gold kit (Zymo Research). Sequencing libraries were generated using the Accel-NGS Methyl-Seq DNA library kit (Swift Biosciences), pooled, and sequenced on a HiSeq 4000 instrument.

### RNA-seq

FACS-purified live cells (typically ∼500,000) or whole islet/SC-islet preparations (typically ∼1million cells) were pelleted and RNA-seq was performed from total RNA depleted for rRNA as previously described (Alvarez-Dominguez et al., 2017a). Directional cDNA libraries were prepared using a stranded RNA-seq library preparation kit (KAPA Biosystems), pooled, and sequenced on a HiSeq 2500 instrument.

### ChIP-seq mapping

Sequencing reads were mapped to the human hg19 reference genome assembly using bowtie2 (Langmead and Salzberg, 2012) with default parameters. Peak calling was performed using MACS2 (Zhang et al., 2008) with default parameters and “--broad --qvalue 0.01” to composite broad regions of read enrichment over background with a minimum FDR q-value cutoff of 0.01, using whole-cell extract from each stage as the background control. H3K4me1 and H3K27ac peaks across all cell types were concatenated and merged whenever overlapping by BEDTools (Quinlan and Hall, 2010) to obtain a unified catalog of H3K27ac/H3K4me1 regions (Figure S4).

### ATAC-seq mapping

Sequencing reads were mapped to the human hg19 reference genome assembly using bowtie2 (Langmead and Salzberg, 2012) with default parameters. Peak calling was performed using MACS2 (Zhang et al., 2008) with default parameters, “--nomodel --shift 37 --extsize 73” to bypass building a ChIP-based shifting model and instead shift read 5’-ends and extend in the 5’->3’ direction by a half-nucleosome size (73bp), and “--broad --qvalue 0.01” to composite broad regions of read enrichment with a minimum FDR q-value cutoff of 0.01.

### WGBS mapping

Sequencing reads were mapped to the human hg19 reference genome assembly using BSMap (Xi and Li, 2009) in bisulfite mode with default parameters. CpG methylation was called as described previously (Ziller et al., 2013), excluding low-quality, duplicate, and >10%-mismatched reads. Only CpGs with >5X coverage, which totaled 23-27million per cell stage, were considered for downstream analyses. Differentially methylated regions (DMRs) were determined using the DSS R package (Feng et al., 2014) with difference>0.2, P<0.05, minimum CpGs=4, and merge regions if closer than 500bp. DMR methylation specificity was quantified as described in (Ziller et al., 2013) based on the Jensen-Shannon divergence from the extreme case in which a region is completely methylated in only one sample and unmethylated in all others or vice versa.

### RNA-seq mapping

Sequencing reads were mapped to the human hg19 reference genome assembly using TopHat2 (Kim et al., 2013) with default and “--min-anchor 5” parameters. Differential gene expression based on GENCODE v19 annotations (Harrow et al., 2012) was determined using HTSeq (Anders et al. 2015) and DESeq (Anders and Huber 2010) as described in (Alvarez-Dominguez et al., 2017a). Expression counts were normalized as counts per million mapped reads (CPM) and only genes with CPM>1 were considered expressed. Differential splicing/promoter/CDS usage was determined using Cuffdiff2 (Trapnell et al., 2013) with default parameters.

### Coverage quantification

ChIP-seq/ATAC-seq/RNA-seq coverage across a region of interest (e.g. DMRs, enhancers) was summarized as the cumulative read pileup across the region, counted using HTSeq (Anders et al., 2015) in “-m union” mode, and normalized as counts per million mapped reads (CPM). WGBS coverage across a region of interest was summarized as the mean CpG methylation across the region; this yielded intermediate methylation levels within active chromatin regions, since hypomethylated CpGs within active chromatin regions are sparse (Ziller et al., 2013).

### Identification of enhancers and super-enhancers

Enhancers were defined as H3K27ac-enriched regions identified by MACS2 that do not overlap gene promoter-proximal (TSS ±2kb) regions. RefSeq annotations obtained from the UCSC genome browser were used to extract TSS ±2kb regions, which were then intersected with H3K27ac broad regions using BEDTools.

Enhancer domains were defined as previously described (Whyte et al., 2013), by linking enhancer sites within 12.5kb of each other and ranking the resulting domains by increasing H3K27ac enrichment. The point for which a line with slope of 1 is tangent to the curve of region H3K27ac enrichment versus region ranking was then used to distinguish super-enhancer vs. typical enhancer domains, as shown in Figure S13A. H3K27ac enrichment was calculated based on the normalized, background-subtracted H3K27ac coverage within stitched-enhancer domains.

A unified enhancer catalog was constructed by merging enhancer domains from all differentiation stages using BEDTools. ATAC, H3K4me1, H3K27ac, RNA and WGBS coverage were quantified at each stage for each enhancer in the catalog, and only enhancers for which the cumulative coverage across all stages was >0 for each of the datasets were used for downstream analyses.

### Enhancer state dynamics

Enhancer state was defined based on H3K4me1 enrichment. For each cell type, enhancer domains where statistically significant H3K4me1 enrichment over background was identified by MACS2 were considered H3K4me1-marked in that cell type. Enhancer activity was analogously defined based on H3K27ac enrichment.

Enhancer gain/loss was determined based on gain/loss of H3K4me1 enrichment supported by significant gain/loss of H3K4me1 coverage as determined by DESeq. Enhancer activation/deactivation was analogously determined based on differential H3K27ac marking.

To investigate enhancer turnover along the differentiation path, we determined, for enhancers gained/lost and activated/deactivated at each differentiation stage, the stage with the maximal change in H3K4me1 and H3K27ac levels, respectively, as shown in Figures S7D and S7E.

Enhancers activated between two differentiation stages that were H3K4me1-marked in the first stage were deemed to undergo a primed-to-active transition. To investigate turnover of enhancer priming along the differentiation path, we determined, for enhancers activated at each stage that were H3K4me1-marked in some previous stage, the stage with the maximal change in H3K4me1 levels, as shown in Figure 3A.

Alternative approaches for defining H3K4me1-and H3K27ac-marking (e.g. by using arbitrary read coverage cutoffs) altered the ratio of enhancers gained vs. lost, active vs. inactive, and H3K4me1-premarked vs. previously unmarked within each differentiation stage, but did not change the relative trends across stages and thus did not change our conclusions.

### Clustering analyses

Correlation studies (e.g. pairwise enhancer catalog H3K4me1 correlations between all cell types and replicates) were generated in R (http://www.Rproject.org/; cor function) with default parameters and “method="spearman"” to use the Spearman’s rank-order implementation.

Principal component analysis was implemented in R (prcomp function) with default parameters and “scale=T” to scale the data to have unit variance.

K-means clustering analyses (e.g. enhancer coverage, pioneer TF enrichment) were conducted in R (kmeans function) with default parameters and “algorithm="Lloyd", nstart=100, iter.max=100000” to use the Lloyd–Forgy implementation (LLoyd, 1982) with 100 random starts and maximum 100000 iterations. Data was standardized prior to clustering by using z-scores to compute the number of standard deviations from the mean across conditions for each data point. To determine the appropriate number of clusters (K), we plotted the cumulative within-cluster sum of squared error (WSSE) for a sequence of cluster solutions (K=1 to K=30), and defined the point at which the reduction in WSSE slows dramatically as the optimal solution.

### Transplantation studies

Animal studies were performed in accordance with the NIH Guide for the Care and Use of Laboratory Animals recommendations. All animals were handled according to the Harvard University Institutional Animal Care and Use Committee, approved by the Committee on the Use of Animals in Research and Teaching of Harvard University Faculty of Arts & Sciences, which is AAALAC International accredited, has a PHS Assurance on file with the NIH Office of Laboratory Animal Welfare (A3593-01), and is registered with the USDA (14-R-0128).

Cell transplantations into immunodeficient SCID-Beige mice (Jackson Laboratory) or Rowett Nude rats (Charles River) were conducted as described previously (Pagliuca et al., 2014) with minor modifications: sorted and unsorted cells were re-aggregated in CRML-1066 medium +10% FBS over 2-4 days with feeding every second day. The re-aggregated cell clusters were then resuspended in RPMI-1640 medium (Life Technologies; 11875-093) and kept on ice until loading into a catheter for cell delivery under the mouse kidney capsule.

To retrieve grafts, engrafted kidneys were dissected from freshly euthanized transplanted mice as described previously (Pagliuca et al., 2014). For histological analysis, grafts were fixed in PBS +4% paraformaldehyde overnight, embedded in paraffin, and sectioned at 100um. For ATAC-seq, grafts were washed in saline and dissociated mechanically by trituration with scissors in PBS +50% Accutase for 7min followed by petting up and down with a 16.5gauge needle. Single-cell suspensions were washed with and resuspended in PBS, stained with TSQ at 37°C for 10min, filtered through a 40μm nylon mash into flow cytometry tubes (BD Falcon), and PI-TSQ+ cells were sorted using MoFlo flow cytometers (Beckman Coulter) into PBS +1% BSA (Sigma) on ice.

### Glucose-stimulated insulin secretion assays

Cell clusters sampled from cadaveric or stem cell-derived islet preparations were divided into four parts for triplicate GSIS assays and insulin content determination. Clusters were washed twice with Krebs buffer containing 2.8mM glucose and loaded into 24-well plate inserts (Millicell Cell Culture Insert; PIXP01250), followed by pre-incubation in Krebs buffer containing 2.8mM glucose for 1h to remove residual insulin. Subsequently, clusters were washed with Krebs buffer containing 2.8mM glucose and sequentially challenged with Krebs buffer containing 2.8mM, 20mM, 2.8mM, and 20mM glucose, with a 1h incubation for each concentration and an additional wash between 20mM and 2.8mM to remove residual glucose, followed by incubation with 2.8mM glucose + 30mM KCl for 1h. Clusters were then dissociated using TrypLE Express (Life Technologies) and cells were counted by an automated Vi-Cell (Beckman Coulter). All incubations were carried at 37°C, with supernatant samples collected at the end of each incubation.

Detection of serum human insulin following a glucose challenge in immunocompromised transplanted animals was conducted as described previously (Pagliuca et al., 2014). Animals were fasted for 16h overnight, and the glucose challenge was performed by intraperitoneal injection of 2g D-(+)-glucose/1 kg body weight. Serum was collected both before and 30min after injection through mandibular bleeding using a lancet (Feather; 2017-01). Serum was separated out using Microvettes (Sarstedt; 16.443.100) and stored at −80°C until ELISA analysis.

GSIS dynamics of handpicked cadaveric/SC-islets were assayed in triplicate on an automated Perifusion System (BioRep) as described previously (Blum et al., 2012). Chambers were first perifused with Krebs buffer containing 2.8mM glucose for 1h at a flow rate of 100ul/min and sequentially perifused with 2.8mM, 20mM, and 2.8mM glucose for 15min, 30min, and 15min, respectively, followed by perifusion with 2.8mM glucose + 30mM KCL for 15min.

Insulin levels in supernatant/serum samples containing secreted insulin were processed using a Human Ultrasensitive Insulin ELISA kit (ALPCO Diagnostics; 80-INSHUU-E10) according to the manufacturer’s instructions.

### Immunohistochemistry

Immunohistochemistry of kidney graft sections was performed as described previously (Pagliuca et al., 2014). Sections were subjected to deparrafinization using Histoclear (Thermoscientific; C78-2-G) and rehydrated, them emerged in 0.1M EDTA (Ambion; AM9261) for antigen retrieval, and placed in a pressure cooker (Proteogenix; 2100 Retriever) for 2h. Slides were blocked in PBS +0.1% Triton X-100 (VWR; EM-9400) +5% donkey serum (Jackson Immunoresearch; 017-000-121) for 1h, followed by incubation in blocking solution with primary antibodies overnight at 4°C. The next day, cells were washed twice in PBS, followed by secondary antibody incubation for 2h protected from light. For imaging, secondary antibodies were washed twice with PBS, and slides were mounted in Vectashield mounting medium with DAPI (Vector Laboratories; H-1200), covered with coverslips and sealed with nail polish. Images were taken using a Zeiss LSM 510 microscope (Carl Zeiss).

### Pioneer factor identification

To study pioneer factors across developmental transitions, sites of newly-gained ATAC signal were identified by running MACS2 as detailed above but using a given stage as the treatment and the known/inferred previous stage as the background control:

Treatment=DE, control= Hpsc

Treatment=PP1, control=DE

Treatment=PP2, control=PP1

Treatment=EN, control=PP2

Treatment=SC-β, control=EN

Treatment=PH, control=EN

Treatment=α-cell, control=PH

Treatment=β-cell, control=SC-β

To assess the pioneer ability of TFs highlighted by motif enrichment analysis as detailed below, we downloaded pioneer index and chromatin log odds scores from (Sherwood et al., 2014), as well as chromatin accessibility regulator ranks from (Lamparter et al., 2017). Given motif redundancy between TFs of the same evolutionarily related subfamily (Jolma et al., 2010), we assigned the same motif score to no more than one TF subfamily member (e.g. the RFX subfamily score was only assigned to RFX1).

To examine DNAme dynamics at the binding sites of known pioneer TFs highlighted by our analysis, we downloaded binding peaks determined by ChIP-seq for FOXA1/FOXA2/GATA4/GATA6 in DE cells (Tsankov et al., 2015) and quantified the relative change in mean methylation between the hPSC and DE ±2.5kb from the center of each TF peak using a fixed number of equal-sized bins.

### Core regulatory circuit identification

Core regulatory circuits (CRCs), defined as groups of transcription factors (TFs) encoded by genes associated with super-enhancers that bind the SE associated with their own and each other’s SE to form fully interconnected autoregulatory loops, were identified using CRCmapper.py as described in (Saint-Andre et al., 2016) with minor modifications. Briefly, the set of expressed genes that encode TFs overlapping/proximal to SEs was first identified, then the set of SE-assigned TF genes whose products bind enhancer constituents within their own SE domain, as predicted by FIMO (Grant et al., 2011) based on a database of TF recognition motifs (Saint-Andre et al., 2016), was determined, and lastly groups of autoregulated SE-driven TFs that are similarly predicted to bind each other’s SE were identified recursively.

We verified that motif-based binding predictions are effectively captured by ChIP-seq for OCT4/SOX2/NANOG/MYC in hPSC and SOX17/FOXA2/OTX2/EOMES in DE (Tsankov et al., 2015), as well as for PDX1/ONECUT1/FOXA1/FOXA2 in PP2 (Cebola et al., 2015) (Wang et al., 2015) and FOXA2/NKX2-2/NKX6-1/PDX1 in cadaveric islets (Pasquali et al., 2014), as shown in Figure S14. Many possible CRCs are identified for each differentiation stage (Table S5), and the circuit containing the set of TFs most often represented across the set of possible circuits is selected as the representative CRC model. Of note, TFs without a known recognition motif (e.g. NGN3) are accordingly missing in CRC models.

Testing different parameters for CRC identification (e.g. restricting motif analysis to ATAC-seq peaks within SEs instead of SE constituents, using a different TF motif database (Weirauch et al., 2014), changing the criteria for identifying expressed genes, and changing SE-gene association rules) led to highly similar lists that capture known master regulators of each pancreatic differentiation stage and thus did not change our conclusions.

The expression specificity of CRC TFs was scored by the fraction of the TF’s cumulative expression across all stages represented in that stage (i.e. its fractional expression level). Median specificity scores ranged from 0.06 to 0.10 across differentiation stages. We selected an empirical specificity cutoff of 0.30 that optimally distinguished between known stage-specific factors and known uniformly expressed ones.

To quantify regulatory connectivity, for each SE-driven TF we defined the inward binding of other TFs to its SE as its regulatory IN degree, and the outward binding of the TF to other TFs’ SEs as its regulatory OUT degree. Binding was predicted as before via TF recognition motif scanning within SE domain enhancer constituents using FIMO ad detailed below, and total connectivity was calculated as the sum of the regulatory IN and OUT degrees.

### shRNA studies

Lentiviral vectors carrying independent *LMX1B*-targeting or control shRNAs (GE Healthcare Dharmacon, Inc.) were transfected into mitotic LentiX-293T (Takara Bio; 632180) cells maintained in DMEM medium (Life Technologies) using TransIT transfection reagent (Mirus). Viruses were concentrated 48h and 72h post-transfection by precipitation with PEG-it virus precipitation solution (System Biosciences) overnight at 4°C followed by centrifugation at 3000g for 30min at 4°C, and stored at −80°C until infection.

For lentiviral infection, cell clusters sampled from suspension cultures the day of EN stage induction were dissociated by incubation in PBS +50% Accutase (StemCell Technologies) for 7min, followed by mechanical dissociation by petting up and down in PBS +10μM Y27632. Cells were resuspended at a density of 5-10million cells/mL in EN-stage day 1 medium (BE5 medium +0.25μM Sant1 +0.1μM Retinoic acid (RA) +1μM XXI +10μM Alk5i II +1μM T3 +20ng/ml Betacellulin +10μM Y27632), and Polybrene infection reagent (Santa Cruz) was added at 8μg/mL. Cells were then plated into 6-well plates pre-coated with BD Matrigel Matrix High Concentration (BD Biosciences) and viruses were added dropwise to each well. Cells were incubated at 37°C and fed daily.

For infected cell collection, the day of change to EN-stage day 5 medium cells were dissociated by incubation with TrypLE Express (Gibco) for 7min at 37°C and quenched in BE5 medium +10μM Y27632. For RNA studies, single-cell suspensions were filtered through a 40μm nylon mash into flow cytometry tubes (BD Falcon), and live infected (PI-GFP+) cells were sorted using MoFlo flow cytometers (Beckman Coulter) into PBS +1% BSA (Sigma) on ice. For FACS immunostaining studies, single-cell suspensions were fixed in PBS +4% paraformaldehyde for 10min, washed with PBS, and blocked in PBS +5% donkey serum (Jackson Immunoresearch) +0.1% saponin (Sigma) for 20min. Cells were then stained in blocking buffer +NGN3 primary antibody for 1hr and in blocking buffer +fluorophore-conjugated secondary antibody for 30min, then washed with and resuspended in PBS +5% donkey serum. Stained cells were filtered through a 40μm nylon mash into flow cytometry tubes (BD Falcon), and infected (GFP+) cells were analyzed for NGN3 levels using MoFlo flow cytometers (Beckman Coulter).

### NanoString and qPCR studies

Total RNA from sorted cells was isolated using a miRNeasy Mini Kit (Qiagen), and RNA was stored at −80°C until downstream studies. For Nanostring profiling, 100-300ng RNA was hybridized to a custom nCounter XT probe set, processed and imaged using the Nanostring prep station and nCounter (NanoString Technologies), and analyzed using the Nanostring nSolver software with default parameters and with the geometric mean expression of five housekeeping genes (RPL15, RPL19, UBE2D3, ITCH, and TCEB1) as an internal normalization control. Only samples with sufficient readout complexity (normal distribution of expression estimates) were quantified. For real-time quantitative PCR, cDNA was synthesized with the SuperScriptIII firststrand synthesis kit (Life Technologies) using random hexamers (Thermo Fisher Scientific). PowerUp SYBR Green-based real-time PCR (Life Technologies) was performed in a 7900HT Fast Real-time PCR System (Applied Biosystems) with 18S rRNA as an internal normalization control.

### In vitro synchronization

Cadaveric islets (typically 5×10^3^ IEQ per condition) and SC-islets (typically 5×10^6^ cells per condition) cultured in ABLE 30ml disposable bioreactors (Reprocell; ABBVS03A) underwent the following treatments (individually or in combination) for 1-4 days: 10uM Forskolin (Stemgent; 04-0025) for 1h followed by 23h recovery; 20mM of Glucose (Sigma; G7528) for 12h followed by 12h recovery; 37.5mg/L Insulin (Sigma; I9278) for 12h followed by 12h recovery; 210mg/L Arginine (Sigma; A6969) for 12h followed by 12h recovery. Stringent washes were implemented before change to media without factors to remove residual factors. The same wash/incubation times were used for mock-treated cultures. At the end of the synchronization/recovery cycles, cultures were kept in constant media and samples were taken every 4h for RNA profiling and insulin secretion/calcium influx assays. Supernatant samples were also taken during this period and profiled for dissolved gases using a Stat Profile Prime instrument (Nova Biomedical).

### Calcium influx assays

Calcium influx dynamics of cadaveric or stem cell-derived islet preparations in response to a glucose challenge were assayed as described previously (Kenty and Melton, 2015; Pagliuca et al., 2014). Cell cluster samples embedded in hESC-qualified Matrigel (VWR; 47743-722) were stained with Fluo4-AM (Life Technologies; F-14217) for 45min, incubated for 15min without the dye, and imaged on an AxioZoom V16 microscope (Carl Zeiss) during sequential static incubations with Krebs buffer containing 2.8mM, 20mM, and 2.8mM glucose for 5min, 10min, and 5min, respectively, followed by incubation with 2.8mM glucose + 30mM KCL for 5min. Images were acquired every 17s throughout incubations, and the mean fluorescence intensity of each cluster across the time series was quantified using ImageJ (Schneider et al., 2012).

### RNA rhythmicity analysis

RNA-seq data from synchronized cadaveric/SC-islet cultures were normalized across libraries using the DESeq2 scaling function to estimate library size factors. Only genes with a normalized count >10 across all samples were retained for downstream analyses. Rhythmicity was evaluated using harmonic regression by fitting a truncated Fourier series to z-score standardized gene expression timecourses with the HarmonicRegression R library (Luck et al., 2014), which uses a Gaussian error assumption to calculate a p-value for each gene.

An empirical alternative for establishing statistical significance, implemented by comparing rhythmicity F-statistics under the Gaussian error assumption for the original vs. 100 randomly-shuffled expression profiles for each gene, led to comparable/greater number of cycling genes enriched for the same ontology terms and thus did not change our conclusions.

### Region-based annotations

Genomic regions of interest (e.g. DMRs, enhancers) were associated with nearby genes and analyzed for distance to their TSS, gene ontology (Ashburner et al., 2000) and pathway annotations from the Molecular signatures database (Liberzon et al., 2011) with GREAT (McLean et al., 2010) using the “basal plus extension” setting, which associates regions with the gene regulatory domains they overlap, defined as zones -1kb to +5kb of a gene TSS (regardless of other nearby genes) that are extended to the nearest gene’s regulatory domain but no more than 1Mb. Only annotations meeting a >2 region-based fold enrichment and an FDR q-value <0.05 by both a binomial test over genomic regions and a hypergeometric test over associated genes were considered significantly enriched.

To investigate whether a defined set of regions (e.g. H3K4me1-premarked enhancers) shows statistically significant, concordant differences between two states of interest (e.g. H3K27ac levels in α vs. β cells) we used GSEA (Subramanian et al., 2005) with default parameters and “-metric log2_Ratio_of_Classes”.

To assess the genomic distribution of regions of interest (e.g. H3K4me1/H3K27ac peaks, DMRs), a custom script was used to overlap their coordinates with a database of genic/intergenic regions as described in (Alvarez-Dominguez et al., 2017a).

### Gene set and pathway enrichment analysis

Gene lists ranked by a feature of interest (e.g. expression change, −log10 p-value of rhythmicity) were analyzed for enrichment of genes grouped by biological process ontology or by curated annotations from the Molecular signatures database with GSEA using default parameters and “-metric log2_Ratio_of_Classes”. Pathway annotations from the Kyoto Encyclopedia of Genes and Genomes and the Reactome database were extracted to investigate overrepresentation of pre-defined pathways.

### Motif enrichment analysis

Genomic sequences from regions of interest (e.g. DMRs, de novo ATAC peaks) were searched for matches to a database of TF recognition sites (Saint-Andre et al., 2016) for TFs expressed in the relevant cell type using FIMO (Grant et al., 2011) as described in (Alvarez-Dominguez et al., 2017b) with minor modifications: a Markov model of sequence nucleotide composition was used as the background model for motif matching (to normalize for biased distribution of individual letters in the examined sequences), and motifs with an odds ratio>2 and q-value<0.05 (Fisher’s exact test) relative to 10 randomly-shuffled controls were considered significantly enriched.

### Statistical analyses

No statistical methods were used to predetermine sample size or remove outliers. The statistical difference between two sets of paired count data (e.g. motif matches in test vs. randomly-shuffled sequences) was assessed by a Fisher’s exact test using the fisher.test R implementation with default parameters. For unpaired data, a Shapiro-Wilk normality test was first performed using the shapiro.test R implementation with default parameters; for normally distributed data (e.g. *LMX1B* levels in test vs. control treatments) we then used a two-sided t-test (t.test R implementation with default parameters) to assess confidence on the measured difference of their mean values. For unpaired data that don’t follow a normal distribution, we used a non-parametric Wilcoxon rank sum test to determine if they belong to the same distribution. FDR q-values were obtained by correcting p-values for multiple hypothesis testing by the false discovery rate method using the p.adjust R implementation with default parameters.

### Additional bioinformatics methods

All sequencing reads were quality-checked with FastQC (http://www.bioinformatics.babraham.ac.uk/projects/fastqc/). Genome-wide read density maps were generated by MACS2 using the “--bdg” option, normalized by RSeQC (Wang et al., 2012a) using the “normalize_bigwig.py” function, and visualized using BEDTools and the UCSC genome browser. Signal coverage and signal change surrounding regions of interest (e.g. DMRs, enhancer sites) were visualized using the ngs.plot R package (Shen et al., 2014). Data heatmaps were generated using the heatmap.2 function of the gplots R package (http://CRAN.R-project.org/package=gplots). Network diagrams were generated using Gephi (Bastian M., 2009).

## REFERENCES

Adam, R.C., Yang, H., Rockowitz, S., Larsen, S.B., Nikolova, M., Oristian, D.S., Polak, L., Kadaja, M., Asare, A., Zheng, D., et al. (2015). Pioneer factors govern super-enhancer dynamics in stem cell plasticity and lineage choice. Nature 521, 366–370.

Aguayo-Mazzucato, C., Sanchez-Soto, C., Godinez-Puig, V., Gutierrez-Ospina, G., and Hiriart, M. (2006). Restructuring of pancreatic islets and insulin secretion in a postnatal critical window. PLoS One 1, e35.

Alvarez-Dominguez, J.R., Knoll, M., Gromatzky, A.A., and Lodish, H.F. (2017). The Super-Enhancer-Derived alncRNA-EC7/Bloodlinc Potentiates Red Blood Cell Development in trans. Cell Rep 19, 2503–2514.

Ashcroft, F.M., and Rorsman, P. (2012). Diabetes mellitus and the beta cell: the last ten years. Cell 148, 1160–1171.

Bass, J., and Takahashi, J.S. (2010). Circadian integration of metabolism and energetics. Science 330, 1349–1354.

Blodgett, D.M., Nowosielska, A., Afik, S., Pechhold, S., Cura, A.J., Kennedy, N.J., Kim, S., Kucukural, A., Davis, R.J., Kent, S.C., et al. (2015). Novel Observations From Next-Generation RNA Sequencing of Highly Purified Human Adult and Fetal Islet Cell Subsets. Diabetes 64, 3172– 3181.

Blum, B., Hrvatin, S., Schuetz, C., Bonal, C., Rezania, A., and Melton, D.A. (2012). Functional beta-cell maturation is marked by an increased glucose threshold and by expression of urocortin 3. Nature biotechnology 30, 261–264.

Boyer, L.A., Lee, T.I., Cole, M.F., Johnstone, S.E., Levine, S.S., Zucker, J.P., Guenther, M.G., Kumar, R.M., Murray, H.L., Jenner, R.G., et al. (2005). Core transcriptional regulatory circuitry in human embryonic stem cells. Cell 122, 947–956.

Buenrostro, J.D., Giresi, P.G., Zaba, L.C., Chang, H.Y., and Greenleaf, W.J. (2013). Transposition of native chromatin for fast and sensitive epigenomic profiling of open chromatin, DNA-binding proteins and nucleosome position. Nat Methods 10, 1213–1218.

Cebola, I., Rodriguez-Segui, S.A., Cho, C.H., Bessa, J., Rovira, M., Luengo, M., Chhatriwala, M., Berry, A., Ponsa-Cobas, J., Maestro, M.A., et al. (2015). TEAD and YAP regulate the enhancer network of human embryonic pancreatic progenitors. Nat Cell Biol 17, 615–626.

Chen, T., Ueda, Y., Xie, S., and Li, E. (2002). A novel Dnmt3a isoform produced from an alternative promoter localizes to euchromatin and its expression correlates with active de novo methylation. J Biol Chem 277, 38746–38754.

Cheng, L., Chen, C.L., Luo, P., Tan, M., Qiu, M., Johnson, R., and Ma, Q. (2003). Lmx1b, Pet-1, and Nkx2.2 coordinately specify serotonergic neurotransmitter phenotype. J Neurosci 23, 9961–9967.

Choukrallah, M.A., Song, S., Rolink, A.G., Burger, L., and Matthias, P. (2015). Enhancer repertoires are reshaped independently of early priming and heterochromatin dynamics during B cell differentiation. Nat Commun 6, 8324.

Creyghton, M.P., Cheng, A.W., Welstead, G.G., Kooistra, T., Carey, B.W., Steine, E.J., Hanna, J., Lodato, M.A., Frampton, G.M., Sharp, P.A., et al. (2010). Histone H3K27ac separates active from poised enhancers and predicts developmental state. Proc Natl Acad Sci U S A 107, 21931–21936.

Donaghey, J., Thakurela, S., Charlton, J., Chen, J.S., Smith, Z.D., Gu, H., Pop, R., Clement, K., Stamenova, E.K., Karnik, R., et al. (2018). Genetic determinants and epigenetic effects of pioneer-factor occupancy. Nat Genet 50, 250–258.

Felsenfeld, G., Boyes, J., Chung, J., Clark, D., and Studitsky, V. (1996). Chromatin structure and gene expression. Proc Natl Acad Sci U S A 93, 9384–9388.

Flynn, R.A., and Chang, H.Y. (2014). Long noncoding RNAs in cell-fate programming and reprogramming. Cell Stem Cell 14, 752–761.

Gamble, K.L., Berry, R., Frank, S.J., and Young, M.E. (2014). Circadian clock control of endocrine factors. Nat Rev Endocrinol 10, 466–475.

Gaulton, K.J., Nammo, T., Pasquali, L., Simon, J.M., Giresi, P.G., Fogarty, M.P., Panhuis, T.M., Mieczkowski, P., Secchi, A., Bosco, D., et al. (2010). A map of open chromatin in human pancreatic islets. Nat Genet 42, 255–259.

German, M.S., Wang, J., Chadwick, R.B., and Rutter, W.J. (1992). Synergistic activation of the insulin gene by a LIM-homeo domain protein and a basic helix-loop-helix protein: building a functional insulin minienhancer complex. Genes Dev 6, 2165–2176.

Gifford, C.A., Ziller, M.J., Gu, H., Trapnell, C., Donaghey, J., Tsankov, A., Shalek, A.K., Kelley, D.R., Shishkin, A.A., Issner, R., et al. (2013). Transcriptional and epigenetic dynamics during specification of human embryonic stem cells. Cell 153, 1149–1163.

Gonzalez, A.J., Setty, M., and Leslie, C.S. (2015). Early enhancer establishment and regulatory locus complexity shape transcriptional programs in hematopoietic differentiation. Nat Genet 47, 1249–1259.

Goode, D.K., Obier, N., Vijayabaskar, M.S., Lie, A.L.M., Lilly, A.J., Hannah, R., Lichtinger, M., Batta, K., Florkowska, M., Patel, R., et al. (2016). Dynamic Gene Regulatory Networks Drive Hematopoietic Specification and Differentiation. Dev Cell 36, 572–587.

Gromada, J., Chabosseau, P., and Rutter, G.A. (2018). The alpha-cell in diabetes mellitus. Nat Rev Endocrinol.

Gu, T., Lin, X., Cullen, S.M., Luo, M., Jeong, M., Estecio, M., Shen, J., Hardikar, S., Sun, D., Su, J., et al. (2018). DNMT3A and TET1 cooperate to regulate promoter epigenetic landscapes in mouse embryonic stem cells. Genome Biol 19, 88.

Heintzman, N.D., Hon, G.C., Hawkins, R.D., Kheradpour, P., Stark, A., Harp, L.F., Ye, Z., Lee, L.K., Stuart, R.K., Ching, C.W., et al. (2009). Histone modifications at human enhancers reflect global cell-type-specific gene expression. Nature 459, 108–112.

Hu, W., Alvarez-Dominguez, J.R., and Lodish, H.F. (2012). Regulation of mammalian cell differentiation by long non-coding RNAs. EMBO Rep 13, 971–983.

Iwafuchi-Doi, M., and Zaret, K.S. (2014). Pioneer transcription factors in cell reprogramming. Genes Dev 28, 2679–2692.

Jindal, R.M., Taylor, R.P., Gray, D.W., Esmeraldo, R., and Morris, P.J. (1992). A new method for quantification of islets by measurement of zinc content. Diabetes 41, 1056–1062.

Kim, D., Pertea, G., Trapnell, C., Pimentel, H., Kelley, R., and Salzberg, S.L. (2013). TopHat2: accurate alignment of transcriptomes in the presence of insertions, deletions and gene fusions. Genome Biol 14, R36.

Langmead, B., and Salzberg, S.L. (2012). Fast gapped-read alignment with Bowtie 2. Nat Methods 9, 357–359.

Lara-Astiaso, D., Weiner, A., Lorenzo-Vivas, E., Zaretsky, I., Jaitin, D.A., David, E., Keren-Shaul, H., Mildner, A., Winter, D., Jung, S., et al. (2014). Immunogenetics. Chromatin state dynamics during blood formation. Science 345, 943–949.

Lin, C.Y., Erkek, S., Tong, Y., Yin, L., Federation, A.J., Zapatka, M., Haldipur, P., Kawauchi, D., Risch, T., Warnatz, H.J., et al. (2016). Active medulloblastoma enhancers reveal subgroupspecific cellular origins. Nature 530, 57–62.

Lister, R., Pelizzola, M., Kida, Y.S., Hawkins, R.D., Nery, J.R., Hon, G., Antosiewicz-Bourget, J., O’Malley, R., Castanon, R., Klugman, S., et al. (2011). Hotspots of aberrant epigenomic reprogramming in human induced pluripotent stem cells. Nature 471, 68–73.

Liu, J.S., and Hebrok, M. (2017). All mixed up: defining roles for beta-cell subtypes in mature islets. Genes Dev 31, 228–240.

Luyten, A., Zang, C., Liu, X.S., and Shivdasani, R.A. (2014). Active enhancers are delineated de novo during hematopoiesis, with limited lineage fidelity among specified primary blood cells. Genes Dev 28, 1827–1839.

Marcheva, B., Ramsey, K.M., Buhr, E.D., Kobayashi, Y., Su, H., Ko, C.H., Ivanova, G., Omura, C., Mo, S., Vitaterna, M.H., et al. (2010). Disruption of the clock components CLOCK and BMAL1 leads to hypoinsulinaemia and diabetes. Nature 466, 627–631.

McCall, M., and Shapiro, A.M. (2012). Update on islet transplantation. Cold Spring Harb Perspect Med 2, a007823.

Millman, J.R., Xie, C., Van Dervort, A., Gurtler, M., Pagliuca, F.W., and Melton, D.A. (2016). Generation of stem cell-derived beta-cells from patients with type 1 diabetes. Nat Commun 7, 11463.

Neph, S., Stergachis, A.B., Reynolds, A., Sandstrom, R., Borenstein, E., and Stamatoyannopoulos, J.A. (2012). Circuitry and dynamics of human transcription factor regulatory networks. Cell 150, 1274–1286.

Oliver-Krasinski, J.M., and Stoffers, D.A. (2008). On the origin of the beta cell. Genes Dev 22, 1998–2021.

Organization, W.H. (2016). Global Report on Diabetes. In World Health Organization, pp. 83.

Otonkoski, T., Andersson, S., Knip, M., and Simell, O. (1988). Maturation of insulin response to glucose during human fetal and neonatal development. Studies with perifusion of pancreatic isletlike cell clusters. Diabetes 37, 286–291.

Pagliuca, F.W., and Melton, D.A. (2013). How to make a functional beta-cell. Development 140, 2472–2483.

Pagliuca, F.W., Millman, J.R., Gurtler, M., Segel, M., Van Dervort, A., Ryu, J.H., Peterson, Q.P., Greiner, D., and Melton, D.A. (2014). Generation of functional human pancreatic beta cells in vitro. Cell 159, 428–439.

Paige, S.L., Thomas, S., Stoick-Cooper, C.L., Wang, H., Maves, L., Sandstrom, R., Pabon, L., Reinecke, H., Pratt, G., Keller, G., et al. (2012). A temporal chromatin signature in human embryonic stem cells identifies regulators of cardiac development. Cell 151, 221–232.

Parker, S.C., Stitzel, M.L., Taylor, D.L., Orozco, J.M., Erdos, M.R., Akiyama, J.A., van Bueren, K.L., Chines, P.S., Narisu, N., Program, N.C.S., et al. (2013). Chromatin stretch enhancer states drive cell-specific gene regulation and harbor human disease risk variants. Proc Natl Acad Sci U S A 110, 17921–17926.

Perelis, M., Marcheva, B., Ramsey, K.M., Schipma, M.J., Hutchison, A.L., Taguchi, A., Peek, C.B., Hong, H., Huang, W., Omura, C., et al. (2015). Pancreatic beta cell enhancers regulate rhythmic transcription of genes controlling insulin secretion. Science 350, aac4250.

Peschke, E., and Peschke, D. (1998). Evidence for a circadian rhythm of insulin release from perifused rat pancreatic islets. Diabetologia 41, 1085–1092.

Polonsky, K.S., Given, B.D., and Van Cauter, E. (1988). Twenty-four-hour profiles and pulsatile patterns of insulin secretion in normal and obese subjects. J Clin Invest 81, 442–448.

Rada-Iglesias, A., Bajpai, R., Prescott, S., Brugmann, S.A., Swigut, T., and Wysocka, J. (2012). Epigenomic annotation of enhancers predicts transcriptional regulators of human neural crest. Cell Stem Cell 11, 633–648.

Rada-Iglesias, A., Bajpai, R., Swigut, T., Brugmann, S.A., Flynn, R.A., and Wysocka, J. (2011). A unique chromatin signature uncovers early developmental enhancers in humans. Nature 470, 279–283.

Rakshit, K., Qian, J., Gaonkar, K.S., Dhawan, S., Colwell, C.S., and Matveyenko, A.V. (2018). Postnatal Ontogenesis of the Islet Circadian Clock Plays a Contributory Role in beta-Cell Maturation Process. Diabetes 67, 911–922.

Rezania, A., Bruin, J.E., Arora, P., Rubin, A., Batushansky, I., Asadi, A., O’Dwyer, S., Quiskamp, N., Mojibian, M., Albrecht, T., et al. (2014). Reversal of diabetes with insulin-producing cells derived in vitro from human pluripotent stem cells. Nature biotechnology 32, 1121–1133.

Rezania, A., Riedel, M.J., Wideman, R.D., Karanu, F., Ao, Z., Warnock, G.L., and Kieffer, T.J. (2011). Production of functional glucagon-secreting alpha-cells from human embryonic stem cells. Diabetes 60, 239–247.

Rinn, J.L., and Chang, H.Y. (2012). Genome regulation by long noncoding RNAs. Annu Rev Biochem 81, 145–166.

Rorsman, P., Arkhammar, P., Bokvist, K., Hellerstrom, C., Nilsson, T., Welsh, M., Welsh, N., and Berggren, P.O. (1989). Failure of glucose to elicit a normal secretory response in fetal pancreatic beta cells results from glucose insensitivity of the ATP-regulated K+ channels. Proc Natl Acad Sci U S A 86, 4505–4509.

Rorsman, P., and Huising, M.O. (2018). The somatostatin-secreting pancreatic delta-cell in health and disease. Nat Rev Endocrinol 14, 404–414.

Russ, H.A., Parent, A.V., Ringler, J.J., Hennings, T.G., Nair, G.G., Shveygert, M., Guo, T., Puri, S., Haataja, L., Cirulli, V., et al. (2015). Controlled induction of human pancreatic progenitors produces functional beta-like cells in vitro. EMBO J 34, 1759–1772.

Saint-Andre, V., Federation, A.J., Lin, C.Y., Abraham, B.J., Reddy, J., Lee, T.I., Bradner, J.E., and Young, R.A. (2016). Models of human core transcriptional regulatory circuitries. Genome research 26, 385–396.

Sherwood, R.I., Hashimoto, T., O’Donnell, C.W., Lewis, S., Barkal, A.A., van Hoff, J.P., Karun, V., Jaakkola, T., and Gifford, D.K. (2014). Discovery of directional and nondirectional pioneer transcription factors by modeling DNase profile magnitude and shape. Nature biotechnology 32, 171–178.

Shih, H.P., Wang, A., and Sander, M. (2013). Pancreas organogenesis: from lineage determination to morphogenesis. Annu Rev Cell Dev Biol 29, 81–105.

Singer, R.A., and Sussel, L. (2018). Islet Long Noncoding RNAs: A Playbook for Discovery and Characterization. Diabetes 67, 1461–1470.

Smidt, M.P., Asbreuk, C.H., Cox, J.J., Chen, H., Johnson, R.L., and Burbach, J.P. (2000). A second independent pathway for development of mesencephalic dopaminergic neurons requires Lmx1b. Nat Neurosci 3, 337–341.

Stadler, M.B., Murr, R., Burger, L., Ivanek, R., Lienert, F., Scholer, A., van Nimwegen, E., Wirbelauer, C., Oakeley, E.J., Gaidatzis, D., et al. (2011). DNA-binding factors shape the mouse methylome at distal regulatory regions. Nature 480, 490–495.

Stolovich-Rain, M., Enk, J., Vikesa, J., Nielsen, F.C., Saada, A., Glaser, B., and Dor, Y. (2015). Weaning triggers a maturation step of pancreatic beta cells. Dev Cell 32, 535–545.

Tsankov, A.M., Gu, H., Akopian, V., Ziller, M.J., Donaghey, J., Amit, I., Gnirke, A., and Meissner, A. (2015). Transcription factor binding dynamics during human ES cell differentiation. Nature 518, 344–349.

Wamstad, J.A., Alexander, J.M., Truty, R.M., Shrikumar, A., Li, F., Eilertson, K.E., Ding, H., Wylie, J.N., Pico, A.R., Capra, J.A., et al. (2012). Dynamic and coordinated epigenetic regulation of developmental transitions in the cardiac lineage. Cell 151, 206–220.

Wang, A., Yue, F., Li, Y., Xie, R., Harper, T., Patel, N.A., Muth, K., Palmer, J., Qiu, Y., Wang, J., et al. (2015). Epigenetic priming of enhancers predicts developmental competence of hESC-derived endodermal lineage intermediates. Cell Stem Cell 16, 386–399.

Whyte, W.A., Orlando, D.A., Hnisz, D., Abraham, B.J., Lin, C.Y., Kagey, M.H., Rahl, P.B., Lee, T.I., and Young, R.A. (2013). Master transcription factors and mediator establish super-enhancers at key cell identity genes. Cell 153, 307–319.

Xi, Y., and Li, W. (2009). BSMAP: whole genome bisulfite sequence MAPping program. BMC Bioinformatics 10, 232.

Xie, R., Everett, L.J., Lim, H.W., Patel, N.A., Schug, J., Kroon, E., Kelly, O.G., Wang, A., D’Amour, K.A., Robins, A.J., et al. (2013). Dynamic chromatin remodeling mediated by polycomb proteins orchestrates pancreatic differentiation of human embryonic stem cells. Cell Stem Cell 12, 224– 237.

Xu, C.R., Li, L.C., Donahue, G., Ying, L., Zhang, Y.W., Gadue, P., and Zaret, K.S. (2014). Dynamics of genomic H3K27me3 domains and role of EZH2 during pancreatic endocrine specification. EMBO J 33, 2157–2170.

Zhang, J.A., Mortazavi, A., Williams, B.A., Wold, B.J., and Rothenberg, E.V. (2012). Dynamic transformations of genome-wide epigenetic marking and transcriptional control establish T cell identity. Cell 149, 467–482.

Zhang, W., Xia, W., Wang, Q., Towers, A.J., Chen, J., Gao, R., Zhang, Y., Yen, C.A., Lee, A.Y., Li, Y., et al. (2016). Isoform Switch of TET1 Regulates DNA Demethylation and Mouse Development. Mol Cell 64, 1062–1073.

Ziller, M.J., Edri, R., Yaffe, Y., Donaghey, J., Pop, R., Mallard, W., Issner, R., Gifford, C.A., Goren, A., Xing, J., et al. (2015). Dissecting neural differentiation regulatory networks through epigenetic footprinting. Nature 518, 355–359.

Ziller, M.J., Gu, H., Muller, F., Donaghey, J., Tsai, L.T., Kohlbacher, O., De Jager, P.L., Rosen, E.D., Bennett, D.A., Bernstein, B.E., et al. (2013). Charting a dynamic DNA methylation landscape of the human genome. Nature 500, 477–481.

## SUPPLEMENTAL REFERENCES

Alvarez-Dominguez, J.R., Knoll, M., Gromatzky, A.A., and Lodish, H.F. (2017a). The Super-Enhancer-Derived alncRNA-EC7/Bloodlinc Potentiates Red Blood Cell Development in trans. Cell Rep 19, 2503–2514.

Alvarez-Dominguez, J.R., Zhang, X., and Hu, W. (2017b). Widespread and dynamic translational control of red blood cell development. Blood 129, 619–629.

Ameri, J., Borup, R., Prawiro, C., Ramond, C., Schachter, K.A., Scharfmann, R., and Semb, H. (2017). Efficient Generation of Glucose-Responsive Beta Cells from Isolated GP2(+) Human Pancreatic Progenitors. Cell Rep 19, 36–49.

Anders, S., Pyl, P.T., and Huber, W. (2015). HTSeq--a Python framework to work with highthroughput sequencing data. Bioinformatics 31, 166–169.

Ashburner, M., Ball, C.A., Blake, J.A., Botstein, D., Butler, H., Cherry, J.M., Davis, A.P., Dolinski, K., Dwight, S.S., Eppig, J.T., et al. (2000). Gene ontology: tool for the unification of biology. The Gene Ontology Consortium. Nat Genet 25, 25–29.

Bastian M. H.S., Jacomy M. (2009). Gephi: an open source software for exploring and manipulating networks. International AAAI Conference on Weblogs and Social Media.

Cogger, K.F., Sinha, A., Sarangi, F., McGaugh, E.C., Saunders, D., Dorrell, C., Mejia-Guerrero, S., Aghazadeh, Y., Rourke, J.L., Screaton, R.A., et al. (2017). Glycoprotein 2 is a specific cell surface marker of human pancreatic progenitors. Nat Commun 8, 331.

Feng, H., Conneely, K.N., and Wu, H. (2014). A Bayesian hierarchical model to detect differentially methylated loci from single nucleotide resolution sequencing data. Nucleic Acids Res 42, e69.

Grant, C.E., Bailey, T.L., and Noble, W.S. (2011). FIMO: scanning for occurrences of a given motif. Bioinformatics 27, 1017–1018.

Gu, G., Wells, J.M., Dombkowski, D., Preffer, F., Aronow, B., and Melton, D.A. (2004). Global expression analysis of gene regulatory pathways during endocrine pancreatic development. Development 131, 165–179.

Gutierrez, G.D., Bender, A.S., Cirulli, V., Mastracci, T.L., Kelly, S.M., Tsirigos, A., Kaestner, K.H., and Sussel, L. (2017). Pancreatic beta cell identity requires continual repression of non-beta cell programs. J Clin Invest 127, 244–259.

Harrow, J., Frankish, A., Gonzalez, J.M., Tapanari, E., Diekhans, M., Kokocinski, F., Aken, B.L., Barrell, D., Zadissa, A., Searle, S., et al. (2012). GENCODE: the reference human genome annotation for The ENCODE Project. Genome research 22, 1760–1774.

Jolma, A., Kivioja, T., Toivonen, J., Cheng, L., Wei, G., Enge, M., Taipale, M., Vaquerizas, J.M., Yan, J., Sillanpaa, M.J., et al. (2010). Multiplexed massively parallel SELEX for characterization of human transcription factor binding specificities. Genome research 20, 861–873.

Kenty, J.H., and Melton, D.A. (2015). Testing pancreatic islet function at the single cell level by calcium influx with associated marker expression. PLoS One 10, e0122044.

Lamparter, D., Marbach, D., Rueedi, R., Bergmann, S., and Kutalik, Z. (2017). Genome-Wide Association between Transcription Factor Expression and Chromatin Accessibility Reveals Regulators of Chromatin Accessibility. PLoS Comput Biol 13, e1005311.

Liberzon, A., Subramanian, A., Pinchback, R., Thorvaldsdottir, H., Tamayo, P., and Mesirov, J.P. (2011). Molecular signatures database (MSigDB) 3.0. Bioinformatics 27, 1739–1740.

Liu, H., Yang, H., Zhu, D., Sui, X., Li, J., Liang, Z., Xu, L., Chen, Z., Yao, A., Zhang, L., et al. (2014). Systematically labeling developmental stage-specific genes for the study of pancreatic beta-cell differentiation from human embryonic stem cells. Cell Res 24, 1181–1200.

LLoyd, S.P. (1982). Least squares quantization in PCM. IEEE Transactions on Information Theory 28, 128–137.

Luck, S., Thurley, K., Thaben, P.F., and Westermark, P.O. (2014). Rhythmic degradation explains and unifies circadian transcriptome and proteome data. Cell Rep 9, 741–751.

McLean, C.Y., Bristor, D., Hiller, M., Clarke, S.L., Schaar, B.T., Lowe, C.B., Wenger, A.M., and Bejerano, G. (2010). GREAT improves functional interpretation of cis-regulatory regions. Nat Biotechnol 28, 495–501.

Mentlein, R., Gallwitz, B., and Schmidt, W.E. (1993). Dipeptidyl-peptidase IV hydrolyses gastric inhibitory polypeptide, glucagon-like peptide-1(7-36)amide, peptide histidine methionine and is responsible for their degradation in human serum. Eur J Biochem 214, 829–835.

Muraro, M.J., Dharmadhikari, G., Grun, D., Groen, N., Dielen, T., Jansen, E., van Gurp, L., Engelse, M.A., Carlotti, F., de Koning, E.J., et al. (2016). A Single-Cell Transcriptome Atlas of the Human Pancreas. Cell Syst 3, 385–394 e383.

Pasquali, L., Gaulton, K.J., Rodriguez-Segui, S.A., Mularoni, L., Miguel-Escalada, I., Akerman, I., Tena, J.J., Moran, I., Gomez-Marin, C., van de Bunt, M., et al. (2014). Pancreatic islet enhancer clusters enriched in type 2 diabetes risk-associated variants. Nat Genet 46, 136–143.

Quinlan, A.R., and Hall, I.M. (2010). BEDTools: a flexible suite of utilities for comparing genomic features. Bioinformatics 26, 841–842.

Richardson, L., Venkataraman, S., Stevenson, P., Yang, Y., Moss, J., Graham, L., Burton, N., Hill, B., Rao, J., Baldock, R.A., et al. (2014). EMAGE mouse embryo spatial gene expression database: 2014 update. Nucleic Acids Res 42, D835–844.

Roadmap Epigenomics, C., Kundaje, A., Meuleman, W., Ernst, J., Bilenky, M., Yen, A., Heravi-Moussavi, A., Kheradpour, P., Zhang, Z., Wang, J., et al. (2015). Integrative analysis of 111 reference human epigenomes. Nature 518, 317–330.

Schneider, C.A., Rasband, W.S., and Eliceiri, K.W. (2012). NIH Image to ImageJ: 25 years of image analysis. Nat Methods 9, 671–675.

Shen, L., Shao, N., Liu, X., and Nestler, E. (2014). ngs.plot: Quick mining and visualization of next-generation sequencing data by integrating genomic databases. BMC Genomics 15, 284.

Subramanian, A., Tamayo, P., Mootha, V.K., Mukherjee, S., Ebert, B.L., Gillette, M.A., Paulovich, A., Pomeroy, S.L., Golub, T.R., Lander, E.S., et al. (2005). Gene set enrichment analysis: a knowledge-based approach for interpreting genome-wide expression profiles. Proc Natl Acad Sci U S A 102, 15545–15550.

Teo, A.K., Arnold, S.J., Trotter, M.W., Brown, S., Ang, L.T., Chng, Z., Robertson, E.J., Dunn, N.R., and Vallier, L. (2011). Pluripotency factors regulate definitive endoderm specification through eomesodermin. Genes Dev 25, 238–250.

Trapnell, C., Hendrickson, D.G., Sauvageau, M., Goff, L., Rinn, J.L., and Pachter, L. (2013). Differential analysis of gene regulation at transcript resolution with RNA-seq. Nature biotechnology 31, 46–53.

Wang, L., Wang, S., and Li, W. (2012a). RSeQC: quality control of RNA-seq experiments. Bioinformatics 28, 2184–2185.

Wang, Z., Oron, E., Nelson, B., Razis, S., and Ivanova, N. (2012b). Distinct lineage specification roles for NANOG, OCT4, and SOX2 in human embryonic stem cells. Cell Stem Cell 10, 440-454.

Weirauch, M.T., Yang, A., Albu, M., Cote, A.G., Montenegro-Montero, A., Drewe, P., Najafabadi, H.S., Lambert, S.A., Mann, I., Cook, K., et al. (2014). Determination and inference of eukaryotic transcription factor sequence specificity. Cell 158, 1431–1443.

Zhang, Y., Liu, T., Meyer, C.A., Eeckhoute, J., Johnson, D.S., Bernstein, B.E., Nusbaum, C., Myers, R.M., Brown, M., Li, W., et al. (2008). Model-based analysis of ChIP-Seq (MACS). Genome Biol 9, R137.

